# Inkube: An all-in-one solution for neuron culturing, electrophysiology, and fluidic exchange

**DOI:** 10.1101/2024.12.06.627248

**Authors:** Benedikt Maurer, Selina Fassbind, Tobias Ruff, Jens Duru, Giusy Spacone, Theo Rodde, János Vörös, Stephan J. Ihle

## Abstract

Culturing neuronal networks *in vitro* is a tedious and time-consuming endeavor. In addition, how the composition of the culture medium and environmental variables such as temperature, osmolarity, and pH affect the spiking behavior of neuronal cultures is difficult to study using electrophysiology. In this work, we present “inkube”, an incubation system that has been combined with an electrophysiology setup and a fully automatic perfusion system. This setup allows for the precise measurement and control of the temperature of up to 4 microelectrode arrays (MEAs) in parallel. In addition, neuronal activity can be electrically induced and recorded from the MEAs. inkube can continuously monitor the medium level to automatically readjust osmolarity. Using inkube’s unique capability to precisely control the environmental variables of a neural culture, we found that medium evaporation influences the spiking response. Moreover, decreasing medium temperature by only 1.5°C significantly affected spike latency, a measure commonly used to show plasticity in *in vitro* experiments. We finally provide a proof-of-concept experiment for drug screening applications, where inkube automatically and precisely varies the concentration of magnesium ions in the medium. Given its high level of autonomy, the system can record, stimulate, and control the medium continuously without user intervention. Both the hardware and the software of inkube are completely open-source.

**Highlights:**

- Low-cost, open-hardware/open-software electrophysiology setup
- Full incubation solution: Temperature, CO_2_, and humidity control
- Perfusion system for automatic fluidic exchange and drug testing with volume feedback

## Introduction

The overarching goal of neuroscience is to understand how the brain works. In order to make scientific progress, reliable and reproducible experiments are of utmost importance (1–3). Given the high level of complexity of the brain, it can be difficult to control extraneous variables. Parameters such as attention (4), sleep (5), and nutrients (6) all fall within this regime and affect reproducibility. Instead of studying the whole brain, these effects can be mitigated by working with dissociated cultures, which decrease system complexity. Thereby tracking and controlling extraneous variables becomes easier. In such *in vitro* model systems, the focus of study ranges from single channels (7) to brain slices (8).

*In vitro* neuroscience can study multiple objectives, such as drug discovery (9–11), the study of neurological diseases (12), neurodevelopmental studies (13), and plasticity research (14–18). For all of these experiments, experimental reproducibility is important, as without it, generally applicable conclusions cannot be drawn. However, just like the field of top-down neuroscience, bottom-up neuroscience struggles with reproducibility (19–24).

One source of activity changes lies in cell migration and network changes, which can be reduced by applying patterning techniques such as polydimethylsiloxane (PDMS) microstructures. Microstructures allow for directional axonal guidance by physically constraining neuronal growth (25, 26). In addition, the microstructures increase the chances of recording neuronal activity from an extracellular electrode, as they improve signal-to-noise ratio (SNR) and guide the neurites right on top of the electrodes (27).

Another potential source of activity changes lies in both temperature and osmolarity fluctuations, which affects the spiking behavior of neuronal networks. In the past, studies have been performed to stabilize the changes in environment (28). Yet, it is currently not clear, by how much fluctuations in the culture environment affect the neuronal activity.

The neuronal activity can be recorded extracellularly by using active or passive microelectrode arrays (MEAs). Active MEAs are complementary metal-oxide semiconductor (CMOS). The amplification and digitization circuitry is located on the same chip as the electrode array, which enables parallel recording at high spatial resolution from up to over 10,000 electrodes (29). On the other hand, passive MEAs separate the electrode array from the recording electronics. This enables usage of transparent materials and facilitate temperature control, as power dissipating electronics are not in contact with the culture. In order to record and also stimulate neuronal cultures on passive MEAs, specific hardware is required. Commercial solutions, however, tend to be rigid, as the hardware and firmware are not directly available to the researchers. This introduces limitations to the types of experiments that are possible. In addition, commercial solutions are costly and tend to be difficult to combine with other custom-made equipment required for experiments. Furthermore, they often still require additional incubation solutions and only have limited or indirect temperature control.

Recently, the open hardware movement is winning in popularity due to its accessibility, affordability, and customizability (30). This holds true particularly for biomedical science and neuroscience. Open-source multifunctional acquisition systems were already shown in the literature (31–34) that mitigate some of the shortcomings of commercial solutions. Some of them were also turned into commercial products (35).

There is a multitude of wet lab equipment, that comprises open hardware, ranging from the aforementioned electrophysiology setups to incubators (36–38) and perfusion systems, as well as microscopes (39), pipettes (40) and even pipetting robots (41). While many of these approaches focus on one modality, the customizability and modularity allows such open hardware to be combined. For example, O’Leary *et al*. combined a perfusion system and an electrophysiology setup (42). Yet, depending on the complexity of the laboratory equipment, it can be difficult to coordinate multiple components; a critical necessity, when wanting to study the effects in changes in environment on the spiking activity of a neuronal culture. Furthermore, most systems do not easily scale to be operable with multiple MEAs in parallel.

In this work we present “inkube”, which is a low-cost, modular open hardware all-in-one solution for measuring neuronal activity of up to 4 neuronal cultures in parallel. It is built around a system-on-a-chip (SoC) containing a field-programmable gate array (FPGA) and can stimulate and record electrically from 240 electrodes in parallel. inkube contains 4 controllers and actuators to precisely control the temperature of the 4 neuronal cultures independently. In addition, it has an integrated waterbath for active humidity control and a CO_2_ controller. Finally, a perfusion system allows to add and remove 4 different liquids to each of the 4 neuronal cultures. For each neuronal culture active feedback of the culture medium volume is incorporated. inkube has been programmed in VHDL, C, Cython, as well as Python. All controllable parameters of the system can be set through Python for ease of use and maximal integrability of the parameters in experimental paradigms. We present how inkube can be build as well as its operation ranges and stability. We show, how the volume of a neuronal culture can be measured and controlled to a precision of roughly 4 µL through a novel, low-cost system based on a laser and a camera sensor. We demonstrate the effect of culture temperature and osmolarity changes on spiking behavior and illustrate the performance of the perfusion system including its potential for drug screening. The here presented hardware solution addresses and resolves the above mentioned shortcomings. The system has been tested using commercially available MEAs. However, it is also compatible with open-source MEAs (43–45). To our knowledge, inkube is the first incubation solution, which combines with a perfusion system as well as an electrophysiology setup, where all components can be easily controlled through a central control unit.

## Materials and Methods

### A. Hardware description

inkube consists of an electrophysiology setup, temperature controllers, a humidity controller, a CO_2_ controller, a perfusion system, and a volumetric sensor to measure the volume of the culture medium. In total, inkube can measure and control up to 4 MEAs in parallel. The block diagram of the system is presented in Fig. 1A. The system is completely open-source. The assembly instructions as well as the parts-list can be found in Supplementary Information A of the supplementary information (SI). Pictures are shown that help assemble inkube (Fig. S1 - Fig. S39). The total cost of the system is approximately $12,000 USD, of which approximately $4,000 USD are required for the perfusion system. Due to the system’s modularity, the end user does not have to assemble the whole system but can instead only rebuild the required subparts to minimize the cost of inkube even further.

**Fig. 1.**
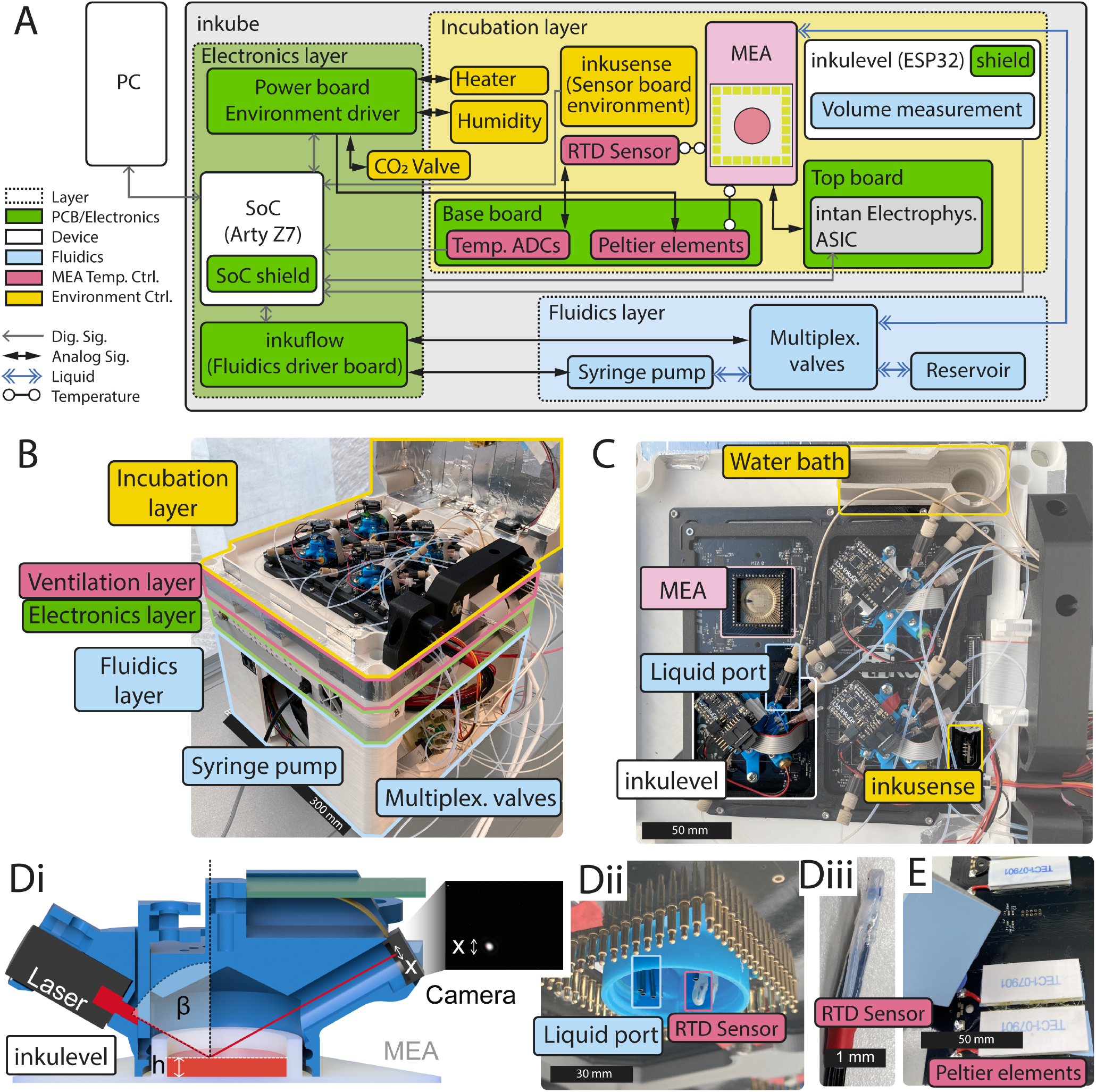
Block diagram and system overview. **(A)** The block diagram of inkube. The different layers of inkube are color-coded. **(B)** Picture of a 3D-printed inkube with an open lid (top-right) showing the different layers. **(C)** Zoom in of the incubation chamber, with 3 mounted inkulevels and one open MEA. Each of the inkulevels is connected to 4 liquid ports. **(D)** inkulevel can be used to measure the medium height in the well. It is a proxy for the overall culture medium volume. The cross section (i) shows the working principle: a laser is reflected by the liquid surface and projected onto a camera. The change of medium height is then extracted from the white dot in the image shown on the right. (ii) shows inkulevel from below without its top board. In (iii), the RTD sensor in its PTFE tube casing is shown. **E** Bottom view of two Peltier elements (white) below the base board are shown, with removed heat sink (left).

inkube uses the Arty Z7 (Digilent, Pullman, WA, USA) development board, which contains a Zynq-7000 SoC (XC7Z020-1CLG400C, Xilinx, San Jose, CA). It is communicating with a GNU/Linux personal computer (PC) via both an Ethernet connection and a USB connection. The Ethernet connection is used mostly to send data from inkube to the PC, while the PC predominantly sends commands to inkube via the USB connection. The Arty Z7 has been programmed in VHDL and C (C11), while the software on the PC side has been written in Python 3 as well as Cython 3. The software layout is presented in Fig. S40. inkube comes with a graphical user interface, which can plot neuronal activity in real time Fig. S43, the status of the system Fig. S44, environment parameters Fig. S45, raster plots Fig. S46, and spike shapes Fig. S47. Python was chosen due to its widespread use in scientific environments as well as its relative ease of use.

All components of inkube fit into a 3D-printed cube. The components are subdivided into 4 layers, where each layer has a unique function. We termed these layers (top to bottom): incubation, ventilation, electronics, and fluidics layer. An overview of the assembled 3D-printed inkube is given in Fig. 1B. The incubation layer contains the 4 MEAs, the sensors, and the electrophysiology components. The ventilation layer is cooling the electronics and is required for temperature regulation. The electronics layer houses most of the electronics. Finally, the components of the perfusion system are located in the fluidics layer, which is the largest and bottom most layer. In the following, each layer is discussed in more detail.

#### A.1 Incubation layer

The incubation layer is the top most layer and houses up to 4 MEAs, sensors, and the electrophysiology setup. The performance of which are described in Section A.4. A picture of the incubation layer is given in Fig. 1C.

The layer can be closed with a 3D-printed lid Fig. S36, which has been electrically shielded using aluminum tape to reduce electromagnetic interference on the electrophysiology setup.

inkube has multiple sensors, which are all placed in the incubation layer. These sensors are **1)** a temperature sensor for the air reservoir inside of the incubation layer **2)** a temperature sensor for each of the 4 MEAs, which measures the temperature of the culture medium in its designated MEA **3)** a humidity sensor, **4)** a CO_2_ sensor of the air reservoir **5)**, and a surface reflection based height sensor we termed inkulevel, which measures the volume of the culture medium of each of the 4 MEAs. inkulevel is shown in Fig. 1D and is discussed further in Section A.5.

The temperature and humidity of the air reservoir is measured with a SHTC3 sensor (Sensirion, Stäfa, Switzerland), while the CO_2_ concentration is measured using a STC31 sensor (Sensirion, Stäfa, Switzerland). In Section A.1, we present the capabilities of the temperature and humidity control. The SHTC3 cannot be used to measure the temperature of the medium reservoirs, as it communicates via digital data protocols which would interfere with the electrophysiology setup. Instead, an analog resistance temperature detector (RTD, PT1000) is being used here in a 4-wire configuration with an impedance measurement IC (MAX31865, Analog Devices, Inc., Wilmington, MA, USA) on the base board. The RTD is wrapped in biocompatible polytetrafluoroethylene (PTFE) tubing to make the sensor water tight. The RTD is mounted inside of inkulevel (see Fig. 1Dii). A disassembled RTD is shown in Fig. 1Diii, while the assembly instructions are given in Fig. S24. In Section A.3 we discuss the dynamics and performance of this assembly.

inkube has multiple actuators to control temperature, CO_2_ concentration, and humidity. Each MEA is located on top of 2 thermoelectric Peltier devices (see Fig. 1E), which can control the temperature of the MEAs independent of each other. This way, MEAs can be both heated and cooled down with respect to the reservoir temperature. The thermoelectric devices are mounted from below on the base board PCB. The reservoir temperature is controlled through resistive elements mounted inside of the lid (see Fig. S36B). In addition, the lid houses fans to circulate airflow inside the reservoir (see Fig. S39 for assembly instructions). The humidity can be controlled through a waterbath, which is heated with a 3D printer cartridge heater (see Fig. S38). How accurately the actuators and sensors can control the reservoir parameters is described in Section A.1.

The incubation layer also houses the electrophysiology system of inkube. In order to record activity from 4 MEAs with each having 59 electrodes and 1 reference electrode, a setup compatible for a total of 240 electrodes is required. In inkube this is achieved by utilizing 16 Intan RHS2116 ICs (Intan Technologies, Los Angeles CA, USA), as Intan ICs are established for recording neuronal activity from *in vitro* cultures (33). The ICs communicate through an SPI bus with the SoC, where the clock and chip select signals are shared between all 16 ICs. The clock frequency of the bus is 12.5 MHz, which enables sampling each electrode at a frequency of 17,361 Hz. For stimulating neuronal cultures, supply voltages of +7 V and -7 V are provided from the electronics layer. The recording and logic chip supply of 3.3 V is generated from a 5 V supply with an additional low-dropout regulator (LDO) in order to decrease the noise affecting the recording electronics. The RHS2116 ICs as well as the LDO are mounted on the PCB termed top board. When inkube is running, the MEAs are sandwiched between the top board and the base board, which houses the thermoelectric actuators.

#### A.2 Ventilation layer

The ventilation layer allows for air flow through inkube. It is required to heat and cool the other layers. Air, sucked in through 4 slits on two opposite sides of inkube, flows past the heat sinks of the Peltier elements (see Fig. S23A for assembly). The flow is powered by a fan at the bottom of inkube (see Fig. S25, which blows the air out through the bottom. To enable undisturbed air flow, inkube is mounted on small rubber feet (see GX5_121_feet in table 1, RND 455-00523, Distrelec Group AG, Nänikon, Switzerland).

#### A.3 Electronics layer

The electronics layer houses most of the electronics of inkube including the SoC that controls the whole system, which is part of the Arty Z7 development board. The onboard ethernet and USB ports are used for communication between PC and inkube. The peripheries of inkube are also connected to the development board. The electronics layer has an additional fan for cooling, which is shown in Fig. S26. On the wall of the electronics layers LEDs are mounted, which visualize the state of the valves of the perfusion system (see Section A.4).

The CO_2_ concentration of the air inside the reservoir is controlled through a valve (PVQ13-6M-06-M5-A, SMC Pneumatics, Tokyo, Japan) placed inside the electronics layer (see Section A.1 for performance). CO_2_ is provided through an external high pressure CO_2_ source at approximately 1.5 bar. The assembly as well as tubing for the CO_2_ is shown in Fig. S37.

There are three PCBs in this layer: **1)** the power board (see Fig. S13), **2)** the perfusion board (see Fig. S17), and **3)** the Arty Z7 shield (see Fig. S12). The power board contains all the electronic components that require high amounts of power, except for the perfusion system. These are the operational amplifiers (PA75CC, Apex Microtechnology, Tucson AZ, USA), that drive the Peltier elements for MEA heating and cooling as well as the reservoir and humidity heater. It also houses the driver of the CO_2_ valve and provides stable power sources to the other PCBs. The voltage levels provided are -7, 3.3, 5, and 7 V. The perfusion board contains the drivers for the syringe pump and the valves required for the liquid multiplexing in the fluidics layer. The syringe pump is controlled by a stepper motor driver (A4988, Pololu Robotics and Electronics, Las Vegas NV, USA), while the valves are controlled by switchable current sources (STP04CM05, STMicroelectronics, Plan-les-Ouates, Switzerland). All of these components are controlled by the SoC via a shift register. Finally, the Arty Z7 shield acts as an interface between the SoC and the top board. Its main task is to transform the CMOS logic signals from the SoC into the low-voltage differential signals that the top board requires for the electrophysiology chips. In addition, it acts as an interface between the SoC and the sensor board, the inkulevels, and the 4 medium-temperature sensors.

#### A.4 Fluidics layer

The fluidics layer is the bottom most layer of inkube and contains a dry bath to chill a 50 mL centrifuge tube through two additional Peltier elements (see Fig. S33). The dry bath is not actively controlled, and once stable, its temperature fluctuates between 4.5 °C and 6.5 °C (see Fig. S59). It further contains a custom perfusion system consisting of a 3D-printed linear perfusion pump designed specifically for inkube and 4 liquid multiplexers. The syringe pump controls up to 4 syringes (1 mL) in parallel and is built on top of a motorized linear stage (DG281-100, RobotDigg Shanghai, Shanghai, China). The assembly of the pump is given in Fig. S28. The perfusion system is capable of delivering 3 different media or drugs to each of the 4 MEAs independently. Simultaneously, it can aspirate culture medium from the MEAs. Therefore, 4 different liquids can be pumped in total.

The liquid path of the perfusion system is shown in Fig. S29. Resin printed multiplexers are used to distribute the 4 different liquids to the relevant MEA. They are printed with biocompatible resin (POWERRESINS SG CLEAR RESIN, 3BFAB BV, Amsterdam, Netherlands). Each of the 4 syringes is connected to a liquid multiplexer. The liquid multiplexer determines which fluidic link is selected. It can either connect the syringe to one of the 4 MEAs or to the corresponding liquid reservoir to refill the syringe. The switching is done through 5 normally-closed 2-way valves (MVL-22-NC-08-14P-PEEK-SIL or for negative pressure MVL-22-NC-08-03P-PEEK-SIL, Memetis GmbH, Karlsruhe, Germany). Since all 4 syringes are controlled by the same syringe pump, one of the 5 paths of the multiplexer needs to be open for all 4 liquids at any given point. Otherwise, pressure will build up in the perfusion system. Hence, each liquid that is not supposed to be pumped into a culture must be connected to its corresponding reservoir. As there are 5 valves per multiplexer and 4 multiplexers, a total of 20 valves is required. For visual feedback, LEDs on the side of the electronics layer indicate which of the valves is currently open (see Fig. S32). Before an experiment, the tubes and multiplexers of all liquids are flushed for cleaning with first 1.4 % sodium hypochlorite in aqueous solution (7681-52-9, VWR International, Radnor, PA, USA) diluted with ultrapure water, followed by a flush with ultrapure water. Afterwards, the tubes and multiplexers were flushed a final time with the type of liquid to be pumped during the experiment, except for the path for liquid retrieval.

#### A.5 Medium level measurement

The medium level measurement system, named inkulevel, provides feedback about the liquid level in the measurement wells with help of a laser (PICO 70125658, Reichelt electronics GmbH, Sande, Germany). We discuss the performance of inkulevel in Section A.2. A cross section of inkulevel is shown in Fig. 1Di. Its PCB is shown in Fig. S20. The height h of the medium defined in this figure is a proxy of the volume of the medium inside the well. While the absolute value is not exact as the surface tension of the medium with regard to the rim of the MEA may have inconsistent curvatures, relative changes can be tracked. The height can be calculated using a camera to capture the reflection point of a laser beam on the water surface. For that, the brightest pixel of the camera image could be used. However, sub-pixel resolution is possible by utilizing the entire shape of the measured reflection and deriving sub-pixel shifts.

To calculate the changes in signal for inkulevel, let *x* be the measured location of the reflection and *β* be the angle of incidence. For inkulevel, the angle of incidence is 60 °. Ignoring surface tension effects, we can calculate the changes in medium level height Δ*x* as:

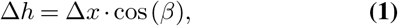

where Δ*h* is the change of the measured reflection dot on the camera sensor. Using an angle of incidence of 60 °we get that changes of the medium height are half as big as the changes on the camera sensor. As a camera, the ESP32-CAM board (Ai-Thinker, Shenzhen, Guangdong, China) with the OV2640 camera sensor (OmniVision Technologies, Santa Clara, CA, USA) is used without a lens. This board is also programmed to perform the image analysis to extract the location of the laser beam (based on (46)), which it then sends to the SoC through a universal asynchronous receiver-transmitter (UART) connection. The laser and the ESP32-CAM board are mounted together with the liquid inlets and outlets on a 3D-printed cap, that can be placed on top of the MEA. The 3D-printed parts are made of Phrozen Aqua Resin and printed on a Sonic Mini 4K (Phrozen, Hsinchu, Taiwan). The mounting steps for inkulevel are given in Fig. S35. A spring loaded screw is used to adjust the pinhole height for the laser in order to compensate for small variations caused by the 3D-printing. When an image is captured by the camera the location of the laser reflection is determined by integrating the pixel values in each row of the sensor. The maximal integral is considered as the center of the laser reflection if it is larger than a threshold (standard value set to 10 *×* 255) and the brightest pixel has a value larger than 200. For subpixel resolution, a weighted average is then taken of the 20 rows around the maximum value, multiplied by 10 and rounded to an integer. Therefore, a difference Δ*x* of 4.5 µm on the sensor translates to 10 bit. The value is then transmitted to the SoC. For an invalid measurement the maximum (6000 as the image resolution is set to 800 by 600 pixels) is transmitted to the SoC.

### B. Cell culturing

Primary hippocampal rat cells or neurons differentiated from human induced pluripotent stem cells (iNeurons) were used in this work. The type of neuron used is specified for each experiment.

MEAs were prepared as described in previous work (47). In short, 60 electrode MEAs were used for this work (60MEA500/30iR-Ti-gr, Multi Channel Systems MCS GmbH, Reutlingen, Germany). MEAs were surface coated with 750 µL of poly-D-lysine (PDL, P6407, Sigma Aldrich, St. Louis, MO, USA) in phosphate buffered saline (PBS, 10010023, Thermo Fisher Scientific, Waltham, MA, USA) at a concentration of 0.1 mg/mL for 45 to 60 min, unless stated otherwise. Afterwards, PDMS microstructures (26) were aligned on top of the MEA using 70 % ethanol in ultrapure water. Each microstructure had 15 independent networks, covering 4 electrodes each in a circular fashion.

Primary hippocampal cells from E18 embryos (Sprague-Dawley rats, Janvier Labs, France)) were dissociated and seeded (seeding density: 150k cells per MEA) using protocols described in previous work (47). Animal experiments were approved by the Cantonal Veterinary Office Zurich, Switzerland. After seeding, cells were cultured in neurobasal™ NB medium. To obtain the complete NB medium, 500 mL of NB (21103049, Thermo Fisher Scientific, Waltham, MA, USA) was supplemented with 5 mL of GlutaMax™ (35050038, Thermo Fisher Scientific, Waltham, MA, USA) and 5 mL of Penicillin Streptomycin (15140122, Thermo Fisher Scientific, Waltham, MA, USA) and stored at 4 °C. Shortly before an experiment, the medium was heated to 37 °C and 2 % v/v B27 supplement™ (17504001, Thermo Fisher Scientific, Waltham, MA, USA) was added.

Human induced pluripotent stem cells (iPSCs), kindly provided by Novartis (Novartis AG, Basel, Switzerland), were differentiated as described in (48). Neurobasal differentiation medium (NBD) was prepared exactly as described in (48) (magnesium concentration of 0.81 mM). The seeding protocol was adapted from (48) by using NBD supplemented with laminin (L2020, Sigma Aldrich, St. Louis, MO, USA) to a final concentration of 5 µg/mL and 2 µg/mL doxycycline (631311, Takara Bio USA, Inc., Mountain View, CA, USA) for the first 7 days after seeding. MEAs were coated with 400 µL of 0.05 mg/mL PDL. Neurons were seeded at a density of 250 k/MEA.

For both types of cell cultures, the medium was exchanged every 3 to 4 days by replacing half of the volume with fresh culture medium.

### C. Signal processing and spike detection

The recorded voltage traces were processed as follows: First, the traces are high-pass filtered with a third order Butterworth infinite impulse response (IIR) filter with a cut-off frequency of 300 Hz (49). Subsequently, high frequency fluctuations are removed with a 5 element Savitzky–Golay filter (50). inkube is utilizing a threshold-based spike detection. The threshold is always computed with a factor X multiplied with the median absolute deviation (MAD) of the electrode specific data. Once a trace surpasses the threshold, a spike event is triggered. For a period of 1 ms the maximum absolute value is being detected. After the period is over, the electrode is blocked for detection for 2 ms and the package ID of the maximum is stored. The detection is performed either directly on the filtered data (with X *∈* {5, 6}) or on data additionally processed with a smoothed non-linear energy operator (with X = 12.5) (51). The non-linear energy operator is computed with a spacing of k_*neo*_ *∈* {3, 4} and smoothed with a Bartlett window of length 4k + 1 (52). Example spike shapes can be found in Fig. S60.

### D. Mg^2+^ Perfusion

To show the capabilities of the perfusion system as well as the potential of inkube to be used for drug testing, different amounts of Mg^2+^-ions where added to the cell cultures, as the effect of Mg^2+^-ions on neuronal activity in culture is well described (53–55). As such, the two different fluids to be added in inkube were chosen to be the NBD medium (Mg^2+^-ions concentration of 0.81 mM) and NBD medium with added Mg^2+^-ions at an increased concentration of 5.81 mM. The third liquid path of inkube was used for culture medium removal. The fourth liquid path remained unused for this experiment. The NBD medium with a magnesium concentration of 5.81 mM was created by additionally adding 0.5 % v/v magnesium chloride (1 M in H_2_O, 63069, Sigma Aldrich, St. Louis, MO, USA).

## Results

In this section, we investigate the electrophysiology performance of inkube while controlling different aspects of the culture environment. For that, we first present the capabilities of inkube to control the culture environment in Section A. Then, we investigate the effect the culture environment has on the neuronal activity in Section B.

### A. Performance validation

In this section we look at how well inkube can control the culture environment.

#### A.1 Reservoir control

inkube can control the temperature, CO_2_ concentration, and relative humidity of the air inside of the incubation layer, the components of which are described in Section A.1 and Section A.3. It takes less than 12 min for the reservoir temperature (see Fig. S49) and CO_2_ (see Fig. S53) concentration to reach a new threshold, once inkube has stabilized. Controlling the humidity of the reservoir takes longer, with a rise time of approximately 1 % per minute (see Fig. S51). Since the air reservoir can only be heated, the temperature range of the air reservoir is between ambient temperature and 17 °C above ambient (at 22 °C ambient temperature, see Fig. S50). The CO_2_ has a range of 0 - 28 % (see Fig. S54), while the humidity can usually be in the range of ambient - 72 % (see Fig. S52). In the rest of this section, we first present how the medium volume can be controlled, followed by the medium temperature.

#### A.2 Medium volume control

inkulevel can be used to measure the level height of the medium inside of the MEA. This is done by imaging a laser point reflected from the water surface by a camera (see Section A.5). We first investigated, how well the location of the impinging point correlates with the volume of the medium. An experiment was performed, where the MEA was filled with ultrapure water and the water volume was controlled directly with the syringe pump. No valve multiplexers were used. The linear behavior of the syringe pump was successfully confirmed beforehand (see Fig. S57). In addition, the hysteresis of the pump was determined to be less than 10 µL (see Fig. S58). The medium volume was then changed in steps of 10 µL within 30 s between 940 and 1000 µL with always returning to baseline for a total of 10 cycles, where the position of the laser point was measured at every step. Constant evaporation was assumed. Hence, the measurement was adjusted by a linear fit. The results of these measurements are presented in Fig. 2A. The relationship between the volume and the measured height is not perfectly linear over the whole measured range. However, it is linear around 1000 µL. The relation is shown by fitting a linear curve on the range from 970 - 1000 µL (R^2^ = 0.99, slope = 6.74 µg/µL).

**Fig. 2.**
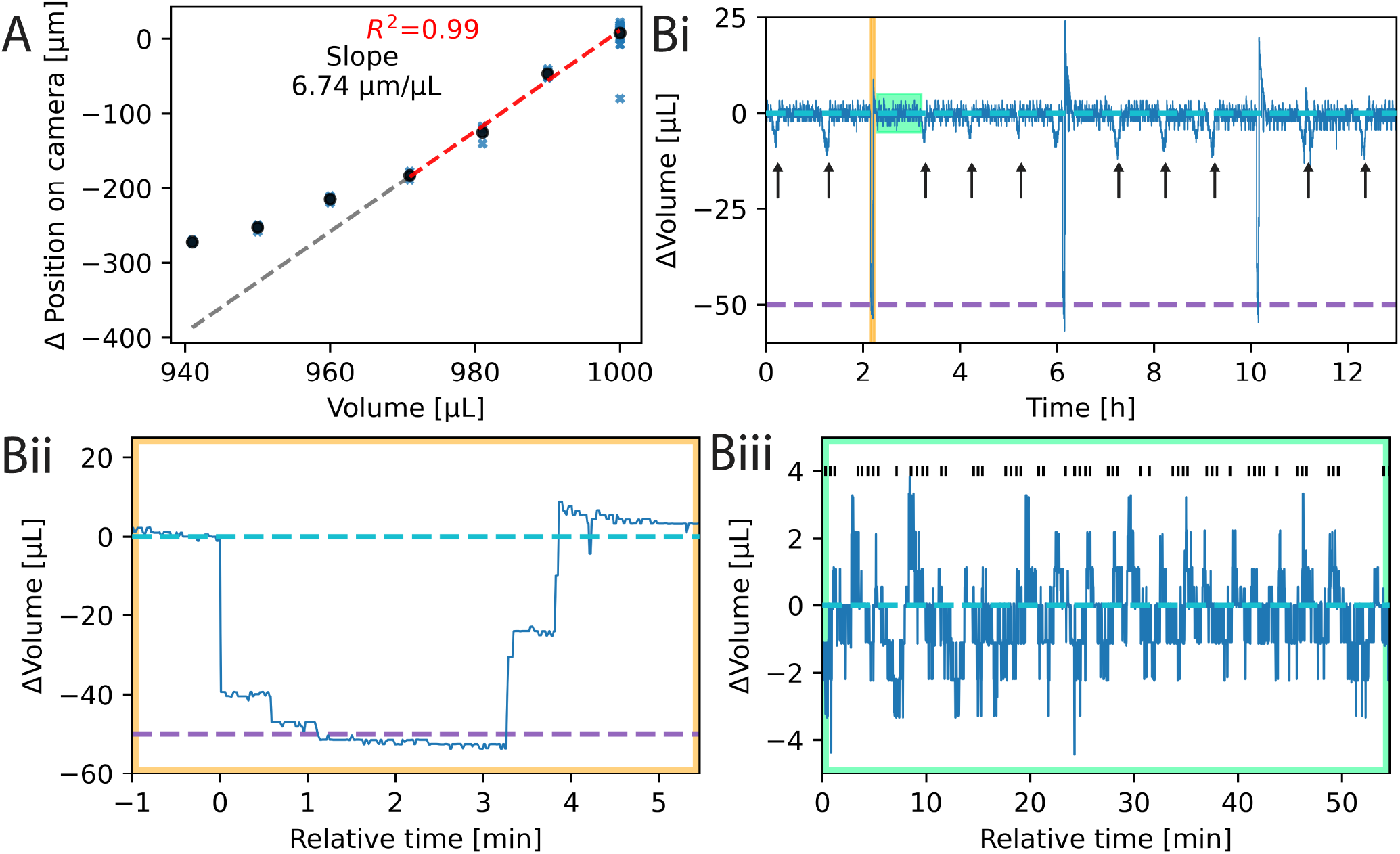
Measuring the volume of the culture medium. **(A)** Relationship between the measured reflection of the laser point (vertical axis) and the volume of the MEA (horizontal axis). A linear fit was applied from 970 to 1000 µL (dashed red line) and extrapolated for the whole range (dashed gray line). **(B)** The measured volume of inkulevel when controlling for evaporation and periodically exchanging medium (i). Evaporation is counteracted by adding ultrapure water, once the signal falls below a threshold (dashed cyan line at 0 µL). Medium is exchanged on other MEAs in the system at the timepoints of the black arrows. Medium for the plotted line was exchanged 3 times during the plotted time interval, which occurred at the timepoints where the measured signal dips below the second threshold line (dashed purple line), which has been calibrated to be 50 µL below the first threshold. Zoom-in of the medium exchange (ii) and the water pumping (iii). Black vertical lines at the top of the subfigure represent time points, when 2 µL of ultrapure water was added. The two zoom-ins of (ii) and (iii) are highlighted in (i) with the yellow and green bounding boxes, respectively.

The theoretical slope considering the MEA dimensions and the pixel size of 4.5 by 4.5 µm is calculated to be about 4.1 µm/µL. This corresponds to the slope of a fit through the section between 950 and 980 µL (4.0µm/µL with R^2^ = 0.98) and can be assumed to be caused by distortion through the surface curvature.

Though the volume-height relationship is not linear across the full range, it is linear for small, adjacent volume intervals and increases monotonically. We used this property to counteract medium evaporation by adding ultrapure water to the MEA whenever the medium height drops below the initial threshold. The resulting measurement can be seen in Fig. 2B, where we controlled the medium level of 4 MEAs in parallel for over 12 h. An exemplary trace of one of the MEAs is given in Fig. 2Bi, with a zoom in of the maintained culture volumes in Fig. 2Biii.

In addition to keeping the medium volume stable, parts of the culture medium were exchanged periodically. For that, 50 µL of the liquid was removed and replaced with fresh medium. For that, a second threshold was used. An exemplary curve of the medium exchange is given in Fig. 2Cii. The exchange of 50 µL takes approximately 4 min. Since inkulevel is sensitive and not linear over the whole measurement range, these two thresholds need to be calibrated every time the MEAs are being taken out of the device and remounted. The calibration consists of 3 cycles. In each cycle the syringe pump removes 50 µL of old medium with an experimentally determined translation factor from liquid volume to steps on the motor as shown in Fig. S57. After a pause of 20 s during which the liquid surface can stabilize, 30 measurements are obtained from inkulevel and the median is stored as a lower threshold. The same procedure is followed after adding 50 µL of fresh medium. Finally, the mean of the 3 cycles is taken as the upper and lower thresholds. The calibration procedure takes about 10 min.

Since there is only one syringe pump for all the liquids in inkube as well as 4 MEAs, only one liquid can be pumped for one MEA at any given time. Hence, while medium exchange is performed for one MEA, evaporation cannot be counteracted during that time for the other MEAs. This effect can be seen in the extended curve in Fig. 2Ci. 50 µL of medium were also exchanged for the other 3 MEAs with equidistant phase shifts. The time points of the exchanges on the other MEAs are highlighted with black arrows.

#### A.3 Medium temperature control

Besides the temperature of the reservoir, the temperature of the medium of each MEAs can be controlled separately. For that an RTD sensor inside of the culture medium measures the temperature, which is then used by a PI controller to heat or cool the MEAs using Peltier elements (see Section A.1). The RTD sensor can be read out as a changing impedance in a 4-point configuration, which is obtained with an ASIC (MAX31865, Analog Devices, Inc., Wilmington, MA, USA) containing an analog-to-digital converter limiting the resolution. The size of the least significant bit around 37 °C is approximately 32 mK. The RTD sensor is placed inside the medium of the culture, to measure the temperature of the culture directly. In many commercially available setups, the temperature of the neurons is either not measured at all, or indirectly, by placing an RTD sensor below. These approaches are less precise than measuring the temperature of the medium directly, as we have shown in the supplementary information, specifically in Fig. S62, Fig. S63, and Fig. S64. From Fig. S64, it is clear that there are considerable temperature discrepancies between the bottom side of the recording headstage and the temperature of the culture medium. In addition, we observed larger fluctuations in temperature (RMSE: 1.268 bits) in case of indirect control if the reservoir temperature is not controlled (see Supplementary Information E) compared to inkube’s approach (RMSE: 0.46 *±* 0.01 bits).

The histogram of the precision error is given in Fig. 3A when maintaining the temperature of the reservoir as well as of the MEAs at 37 °C. Approximately 80 % of all measurements for all 4 MEAs where exact to the least significant bit with respect to the set value, while the other 20 % were off by one bit (recording length 4.5 h). The average RMSE of the signal for all 4 MEAs was approximately 14.7 mK. As a consequence, the control loop of inkube does not introduce any additional measurement errors to the system besides the precision errors of the sensor.

**Fig. 3.**
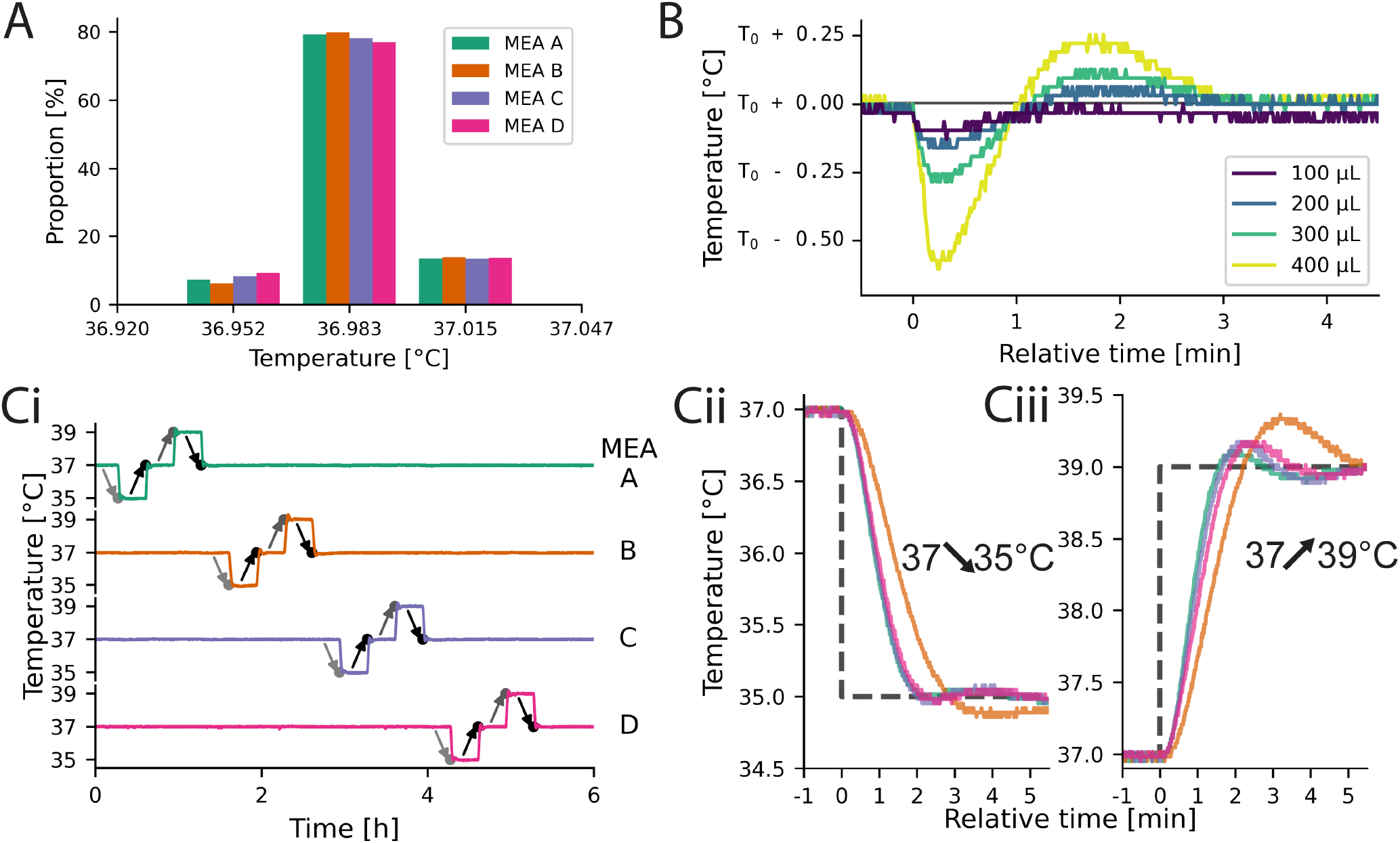
Temperature characterization of inkube. **(A)** Precision error of all 4 MEAs at a fixed threshold of 37 °C. **(B)** Changes in media temperature over time when adding different amounts of liquid at a set temperature of T_0_ =37.185 °C. **(C)** The temperature of all 4 MEAs can be controlled independently as shown by an exemplary square waveform. The measured temperatures of all MEAs are shown in (i), with a set value of 37 °C. The light gray arrows indicate a set value of 35 °C, the black of 37 °C and the dark gray of 39 °C. The falling edges are depicted in (ii), and the rising edges in (iii), each starting at 37 °C and changing by 2 °C as visualized by the dashed lines.

Next it was determined, how much the temperature changes, if medium is added to the MEA. A temperature change is expected, as the added medium is not temperature controlled. For that, 100, 200, 300, and 400 µL of medium were removed and added to an MEA containing 1000 µL total medium. The resulting temperature curves are shown in Fig. 3B. The depicted curves are aligned to the timepoint of added liquid at 0 min. Immediately after adding new medium, the temperature drops. The more liquid was added, the larger the drop (100 µL: -130 mK, 400 µL: -600 mK). Approximately 3 min after the liquid was added, the temperature recovered for all 4 volumes added. This experiment shows, that there is a trade-off between temperature stability and medium exchange speed. The PI controller of the temperature did not receive feedback about any changes in the medium volume. By giving the controller this information, we suspect it will be possible to decrease the size of the initial temperature drop. In addition, by tuning the PI controller, the overshoot and settling time can be tailored to the application needs.

To investigate how well the temperature of each of the MEAs could be controlled independently of the temperature of the other MEAs, step functions were applied to the set values of the temperatures. First, the temperature was decreased from 37 °C to 35 °C. 20 min later, the set value was changed back to 37 °C. Another 20 min later, this process was repeated, but this time with 39 °C instead of 35 °C. As a consequence, there are a total of two 2 K down steps and two 2 K up steps. These step functions were applied to all 4 MEAs sequentially. The total experiment took 5 h and 20 min and the measured temperatures of all 4 MEAs are shown in Fig. 3Ci. The plot does not only show, that the temperatures of the 4 MEAs can be controlled independently even for relatively large temperature differences of 2 K, but also how fast the temperatures of the MEAs adapt to a change of the set value. A zoom-in of the steps from 37 °C to 35 °C and 39 °C for all 4 MEAs are shown in Fig. 3Cii and Fig. 3Ciii, respectively. While the 4 MEAs have slightly different temporal dynamics, all settle within 5 min. Given a reservoir temperature of 37 °C, the limits of the medium temperature are slightly above *±*10 K in both directions (see Fig. S55 and Fig. S56).

#### A.4 Electrophysiology

The electrophysiology setup was introduced in Section A.1. The input referred noise of the electrophysiology chips for the raw signal is 3.21 µV_*rms*_ (median of all electrodes, minimum 2.22 µV_*rms*_), while for the filtered signal it is 2.41 µV_*rms*_ (median of all electrodes, minimum 1.54 µV_*rms*_). This noise performance is comparable to the expected performance of the RHS2116 ICs.

While closed-loop stimulation, meaning stimulating the culture dependent on its activity, is outside the scope of this work, inkube has this capability. The minimal tested round trip time for closed-loop stimulation periods was determined to be 10 ms, if the loop is closed through the host PC and Python interface. Faster round-trip times are achievable by closing the loop either in the Cython code for spike detection or directly inside the SoC.

### B. Effects of culture environment on neuronal activity

After characterizing the actuators of inkube in the previous section, we now present, how inkube can be used to investigate the effects of the culture environment on the neuronal spiking activity. For that, we present 3 different exemplary experiments, which show the capabilities of inkube. First, we look at temperature dependent changes in spike timing, followed by the effect of medium evaporation on the spiking activity. Finally, we end this section with a model drug testing experiment, during which we automatically change the concentration of a model compound (here Mg^2+^-ions) and investigate the effects on the spiking activity.

#### B.1 Relationship of temperature and spike timing

Endogenous neuronal activity was recorded at different temperatures to investigate the effect of culture temperature on spiking behavior. The microstructure used in this work together with the color-code of the corresponding electrode arrangement is presented in Fig. 4A (47). The spike timing was analyzed using raster plots, that are synchronized to neuronal activity recorded on a particular electrode. An example of this method is shown in Fig. 4B with synthetic data. Each network consists of 4-nodes with 4 corresponding electrodes aligned to the connecting microchannels. One of the 4 electrodes is chosen as the triggering electrode (here red). Then, the spike data is split into segments based on the spiking activity of the triggering electrode. A segment is defined as an area of a fixed length (*e.g*. 5 ms), which starts at the spike on the triggering electrode, provided that there are no other spikes recorded on the triggering electrode in a window of 10 ms before. We call such a spike a valid triggering spike. In the shown example, 3 valid triggering spikes with their corresponding segments (highlighted in yellow, orange, and pink) are shown. The spikes in each segment are then plotted in a horizontal line, where a spike recorded on each electrode has its unique color. The vertical location of the horizontal line depends on the start time of the segment, *i.e*. the timestamp of the trigger spike. We call the resulting raster plot a spike-time-triggered raster plot (STTRP). Similarly to stimulation triggered raster plots, if the networks have consistent spiking patterns, vertical bands will occur in the STTRP. Shifts in relative spike timing can be detected by shifts in the latency of the band.

**Fig. 4.**
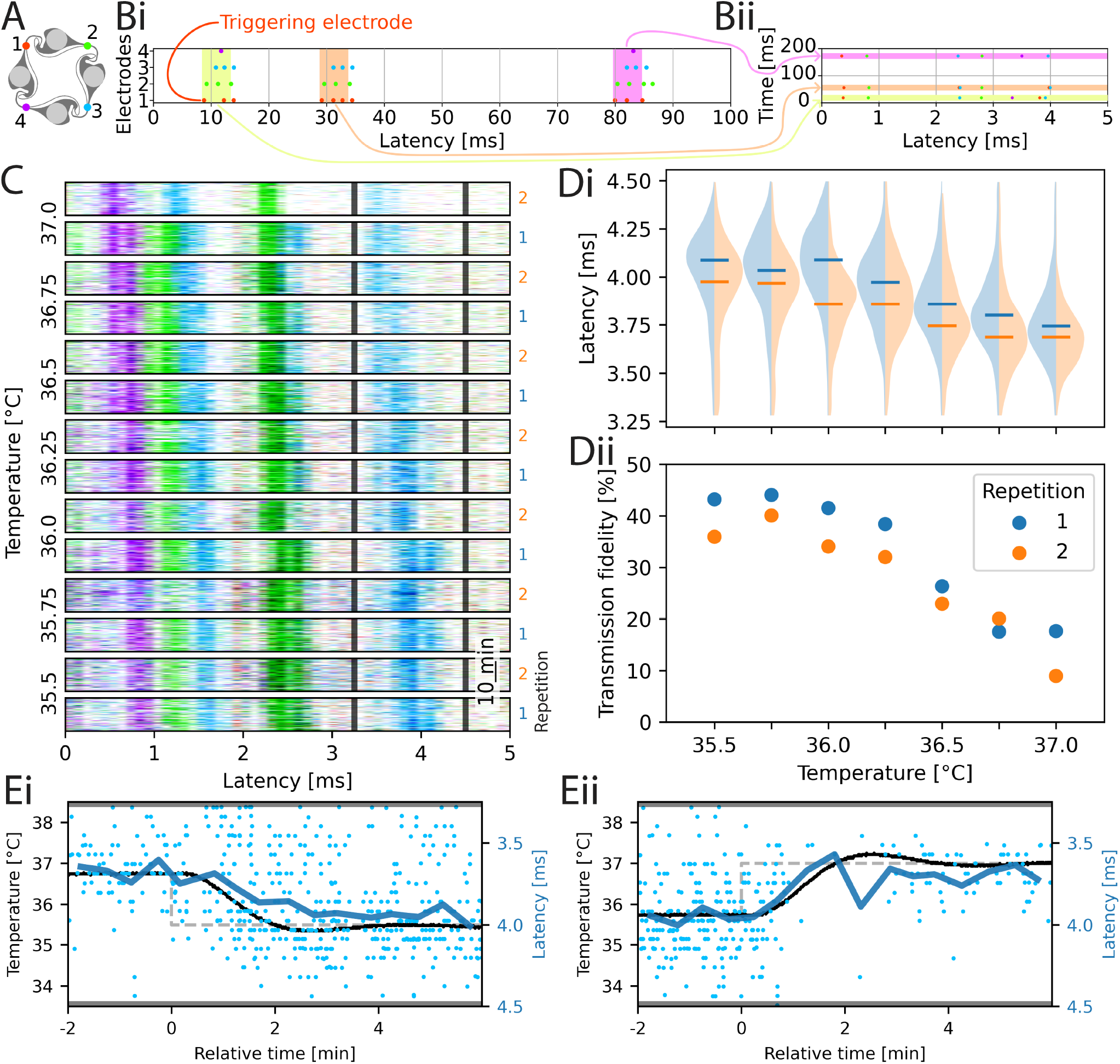
Spike-triggered Time Raster Plots at varying culture temperature. **(A)** The network architecture and the electrode colors of the 4 electrodes used in this work. **(B)** Graphical explanation of how raster plots are being created. Given a network with 4 electrodes (color-coded), consider (i) an exemplary spiking response of the network, where each line is one electrode. Then, a triggering electrode is chosen (here red). Each time this electrode spikes without there being spikes on other electrodes in a window before that, plot the spikes in a short time window after the triggering spike in a row at the vertical location of the time the spike occurred (ii). The result is a spike-time-triggered raster plot (STTRP). **(C)** The STTRP of the network for the culture temperature indicated on the left. To improve the quality of the STTRP, spikes on the triggering electrode where only considered, if there where no other spikes on any electrode in the previous 10 ms. The temperature was changed randomly during the experiment but it was reordered here for better interpretability. For unordered data see Fig. S65. Cell type: primary cortical rat cells (DIV 72). **(D)** Temperature dependence of the latency (i) and transmission fidelity (ii) of the blue band marked with the grey limits in (C). The violin plots in (i) show the distribution of the latency (blue: first repetition, orange: second repetition), with the median for each repetition marked by a horizontal line in the same color. The dots in (ii) show the average transmission fidelity, *i.e*. the probability of a spike occurring within the band limits after an eligible trigger spike, during the first (blue) and second (orange) cycle. **(E)** Correlation between temperature and latency over time. The measured temperature is plotted in black, while the set value for the temperature is plotted in gray (dashed). The latency has been binned (blue dots, binsize: 30 sec) and filtered (blue line).

STTRPs were used to investigate the effects of temperature on spike timing, by varying the temperature of the culture in 250 mK steps from 35.5 °C - 37 °C. The temperature of a culture of primary hippocampal neurons was changed in a random order such that biases due to overall drifts in the spike timing could be prevented. Each temperature level was held for 20 min. In addition, after each temperature level was reached once, the experiment was repeated a second time with a different random order of the levels. The STTRPs sorted by temperature are given in Fig. 4C. Only the last 17 min for each step with stable temperature are shown. The unordered activity is provided in Fig. S65. The red electrode was chosen as a triggering electrode. In Fig. 4D, the changes of latency of the blue band marked by the latency limits of 3.25 ms and 4.5 ms (black lines in Fig. 4C) as well as the transmission fidelity, which is defined as the probability of obtaining a spike on the blue electrode within the latency limits after a valid triggering spike, are shown. A similar analysis has been performed for different electrodes and band limits in the supplementary information (see Fig. S66).

In Fig. 4Di, the latency decreases as the temperature increases (p-value < 0.05, Mann-Kendall trend analysis (56)). The same significant effect could be observed on the other 3 bands investigated in the supplementary information (see Supplementary Information E). The transmission fidelity in Fig. 3Dii decreased, as the temperature increased (p-value < 0.05, Mann-Kendall trend analysis). By combining the p-values (Mann-Kendall trend analysis) of both repetitions with the Fisher method, the transmission fidelity of one band had no significant trends (Fig. S66A, p-value: 0.125), one was significantly increasing (Fig. S66B, p-value: 0.025), and one was significantly decreasing (Fig. S66A, p-value: 0.002) as the temperature increased. Consequently, the transmission fidelity can both correlate and inversely correlate with the temperature.

To investigate the relationship between the temperature and the shift in latency, we focus on two transient transitions within the data presented in Fig. 4C. The transitions are shown in Fig. 4E, where the set value of the temperature is plotted as a dashed gray line, while the actual temperature is plotted as a continuous black line. The temperature plot here is superimposed with the averaged latency (blue curve, mean of spikes binned to 30 s windows). The observed nearly perfect alignment between the temperature of the MEA and the latency indicate a direct causality correlation between these two parameters as opposed to indirect effects through spiking adaptations (*e.g*. plasticity) within the culture.

#### B.2 Relationship of evaporation and spiking activity

It is known that counteracting evaporation in cell cultures increases cell viability (28). Yet, it is not clear how evaporation affects neuronal activity. Furthermore, it is often implicitly assumed that evaporation, which influences the medium’s osmolarity, does not impact an experiment significantly. Hence, evaporation is usually not counteracted during an experiment. Here, we show how inkube can test the validity of this assumption.

We used a primary hippocampal culture to look at the effect of evaporation on neuronal activity. The activity was recorded for approximately 6.5 h, during which the medium volume was kept stable for the first 22 min by adding ultrapure water thereby avoiding any increase in the concentration of medium constituents through evaporation. Afterwards, no water was added for 4 h. The reservoir temperature was set to 34 °C, to speed up the evaporation process. The medium evaporated at a rate of approximately 0.18 µL/min during the experiment, as calculated from the relationship presented in Fig. 2A. Then, the controller that adds ultrapure water to the system was turned on and water was added to the medium again. After approximately 34 min, the medium volume was at its initial state and was maintained for the remainder of the experiment. The measured medium height is plotted in Fig. 5A.

**Fig. 5.**
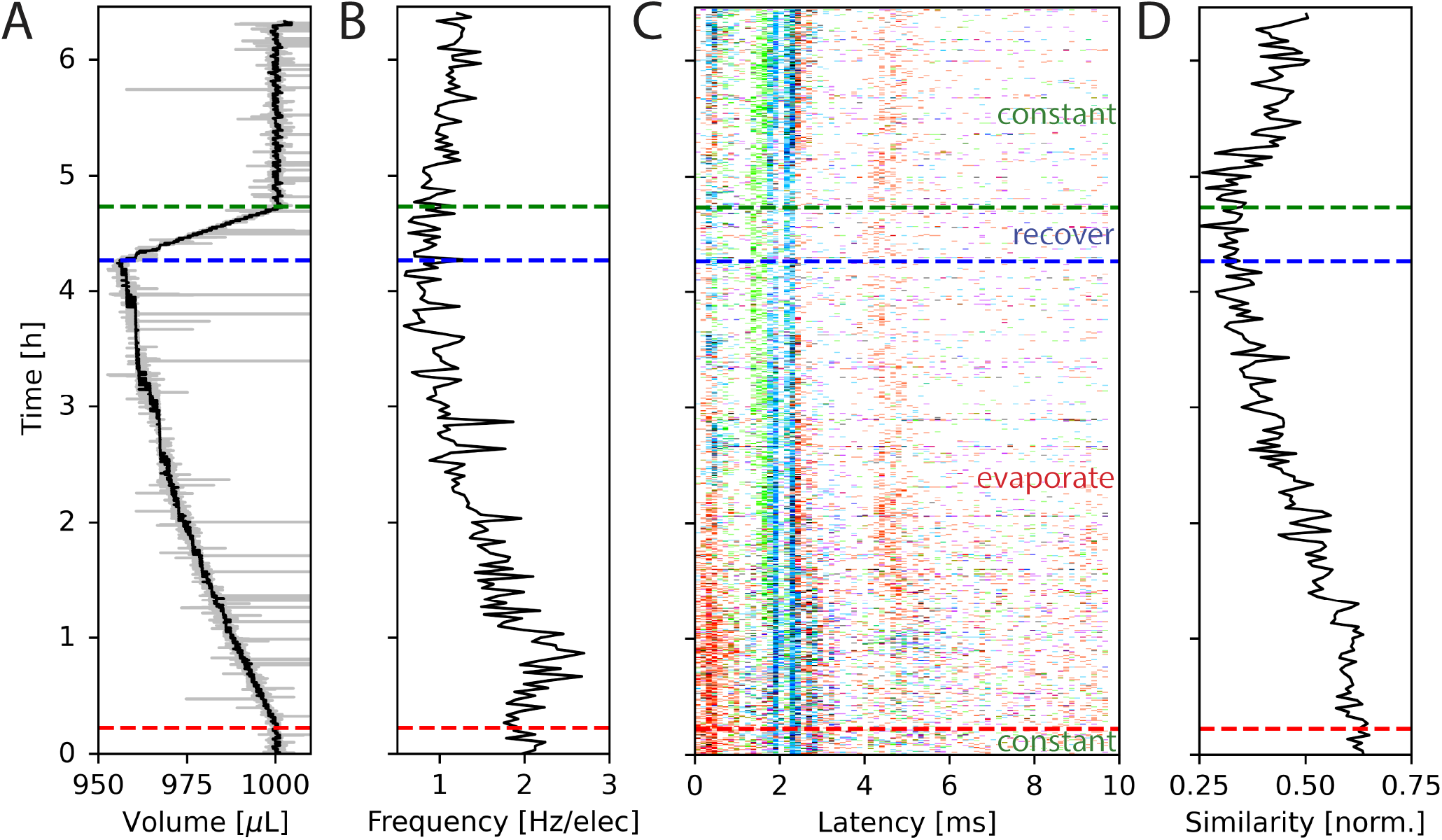
Changes in spontaneous network activity during medium evaporation. **(A)** The volume in the MEA well during the experiment as measured by inkulevel. The return value of inkulevel is translated according to the linear fit in Fig. 2A. **(B)** Averaged spike frequency of all electrodes during the experiment. **(C)** STTRP of a network triggered on the cyan electrode, where valid triggering spikes have no spikes on the same electrode 5 ms before. For the first approx. 20 min, the medium level was held constant, followed by approx. 4 h during which the medium could evaporate freely (red dashed line). Afterwards, the medium level was controlled again (blue dashed line), which took approx. 30 min to recover (green dashed line). **(D)** Similarity metric of the spike response over time. The similarity metric is defined in Equation 3. In short the similarity metric measures how closely the spiking patterns of two time periods match, with a value of 1 indicating identical patterns and lower values indicating less similarity. Cell type: primary cortical rat cells (DIV 206).

The average spike frequency of the circuit is plotted in Fig. 5B. The frequencies were binned into sections of size 2 min. Initially, the average spike frequency is approximately 2 Hz. Once the evaporation is not counteracted anymore, the average spiking frequency slightly increases at first, but then steadily decreases over a process of 4 h to approximately 1 Hz just before the medium level is being recovered to its initial state. For the rest of the experiment during which the medium level is held constant again, the spiking frequency slowly recovers but does not reach its original value anymore. It is not clear, why this is the case. Multiple possible explanations exist, such as long lasting changes in neuron and network dynamics caused by the changes in osmolarity, cell death, or insufficient recovery time. This experiment shows, that the spiking frequency can half over the progress of 4 h of recording, if evaporation is not being considered.

#### B.3 Model experiment for fully automated drug testing

Besides changes in spiking frequency, we were also interested in the changes in the spiking pattern caused by medium evaporation. For that, we focused on the STTRP, which is shown in Fig. 5C. The STTRP was plotted for a segment length of 10 ms after the triggering spike assuring no preceding spikes on the triggering electrode for 5 ms. Qualitatively, changes in the STTRP can be observed, where bands with latencies below 4 ms were fading out during uncontrolled evaporation, while the red band at a latency of around 4 ms has shifted to an earlier latency as the medium evaporated. Both effects were somewhat inverted as the water level was readjusted. For a more quantitative analysis, we looked at the 4 STTRPs of this network, which were created by choosing each of the 4 electrodes as the triggering electrode. Then, the STTRPs were binned for each electrode independently to 0.1 ms for the latency and 2 min for the trigger spike time. We define the resulting STTRP as the vector 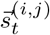, where *i*∈{1, 2, 3, 4} is the triggering electrode, j *∈* {1, 2, 3, 4} the observation electrode considered in the STTRP, and *t* ∈ {0, 1,…, 193} the index of the time bin. The similarity S^(*i,j*)^(t_0_, t_1_) between 2 STTRPs was then defined as:

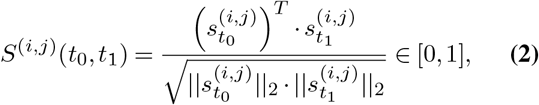

where ||·||_2_ is the L_2_-norm. The normalization term ensures the metric independent of the overall spiking frequency. The similarity is 1 if and only if the two binned STTRPs express an identical normalized spiking pattern. We then looked at the overall similarity 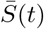 of the binned STTRPs with respect to the initial phase, during which the medium level was held constant:

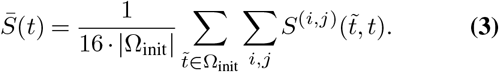

Here, Ω_init_ is the set of all bins within the first 22 min (initial constant phase). The number 16 is used to normalize over the number of contributing STTRPs, hence the 4 observation electrodes j and 4 triggering electrodes i. The overall similarity 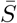 is plotted in Fig. 5D. As the entire initial phase is used for comparison with shorter time windows of the same period, it is expected that a value of 1 is only reached, if the spiking activity over all time bins is exactly identical. Due to uncorrelated spontaneous activity, this is unlikely. In fact, during the initial phase, the overall similarity was only approximately 0.63. The value then gradually decayed to approximately 0.33 within the following 4 h, during which the medium was evaporating. Once the medium level recovered, the overall similarity increased again to approximately 0.45. Just like the spiking frequency, the overall similarity did not fully recover. Furthermore, the overall recovery of the similarity metric did not instantaneously reflect the medium level change. Instead, recovery was delayed. Potential causes are similar to the ones discussed for the spiking frequency.

We performed a model experiment in the style of a recently published dose-response curve to illustrate the capabilities of inkube for drug testing. We automatically changed the concentration of Mg^2+^ in the culture medium of a human iNeuron culture. The starting concentration of 0.81 mM was consecutively increased up to 3.81 mM in steps of 1 mM. For each concentration level, the network was stimulated on the cyan electrode at 4 Hz for 10 min (bi-phasic rectangular, current peak-peak: 320 nA, stimulus halfwidth: 346 µs). Afterwards, 50 % of the medium was automatically exchanged 3 times with fresh medium at a magnesium concentration of 0.81 mM, resulting in a final concentration of 1.19 mM. Finally, the circuit was stimulated twice for 10 min each. In between each stimulus period and medium exchange, 5 min intervals with spontaneous recording were kept, for which the data is presented in the supplementary information (see Fig. S67). The experiment is sketched in Fig. 6Ai, while each of the raster plots of the stimulation response is given in Fig. 6Aii. Once the experiment was setup, it did not require user interaction and ran for approximately 2 h.

**Fig. 6.**
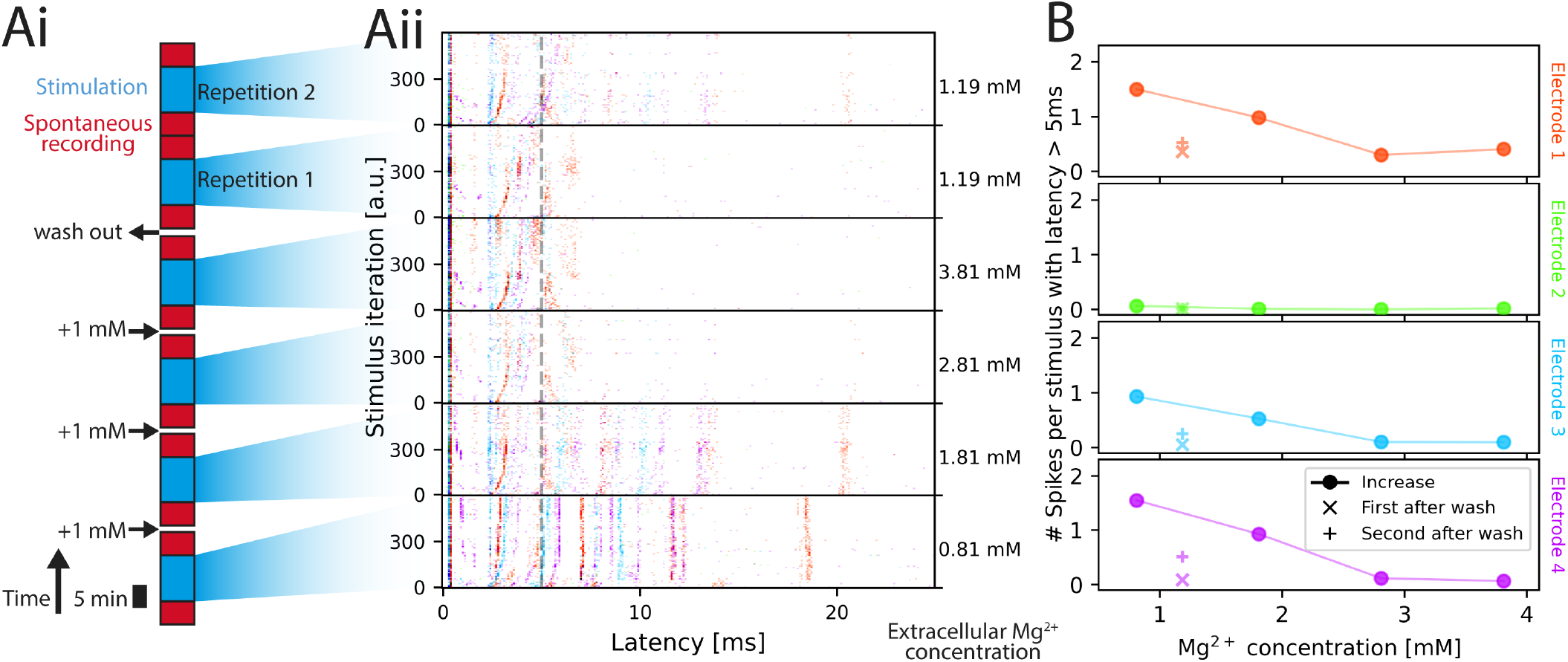
Network stimulation response at different magnesium concentrations. **(A)** Experimental setup and stimulation-induced responses. Magnesium concentration is increased in 3 steps (1 mM each) followed by a wash out (i). After each concentration change, 5 min of spontaneous recording was done, followed by 10 min of repeated stimulation (4 Hz). Afterwards, 5 min of spontaneous activity was recorded again. Post stimulus raster plots following the color code of Fig. 4 is shown in (ii). Stimulation with 160 nA amplitude is applied for 10 min at 4 Hz iterating over the 4 electrodes. The Mg^2+^ concentration in the medium is shown on the right. **(B)** Average spike count with respect to the Mg^2+^ concentration. The spike count of all spikes occurring with a latency of at least 5 ms after stimulus is plotted for all 4 electrodes (4 plots) color-coded by the color of the electrode (see Fig. 4A). The values corresponding to the increasing magnesium concentrations created during the initial 3 steps are plotted with full symbols, the average spike count after washout with + for the first wash, x for the second wash).

Qualitatively, the neuronal activity presented to an external stimulus in Fig. 6Aii decreased and the latency of bands increased as the Mg^2+^ concentration increased. As the ions were washed out, the spiking response recovered, albeit not to the previous levels. This behavioral response aligns with the one observed in previous work (53). To allow for quantitative comparison, the average number of spikes per stimulus occurring with a latency of more than 5 ms after the stimulus was plotted in Fig. 6B for each of the 4 electrodes. The cutoff of 5 ms was chosen to eliminate spike of directly responding neurons (57). The spikes that occur with delays longer than 5 ms are more likely caused by synaptic connections rather than directly by the stimulus. The post-stimulus time histogram of the 4 electrodes including the first 5 ms is presented in Fig. S68. The observed response, when removing the first 5 ms, is similar to previous work (53), where the number of spikes inversely correlates with the magnesium concentration prior to wash out. Post wash out, the second repetition has a higher spike count than in the first repetition and the maximum concentration, which makes it likely that neuronal cultures need time to adapt to the new environment.

## Discussions and outlook

In this work, we have presented inkube, an electrophysiology setup with an integrated incubator and perfusion system. inkube has control over the reservoir temperature, CO_2_ concentration, and relative humidity. In addition, it can independently control the medium temperature of 4 MEAs. We have shown that inkube is capable of maintaining the medium level in the wells of the MEAs. In addition, it can exchange the medium automatically.

inkube is of particular interest in experimental paradigms, where the focus lies on changes in spiking behavior. Particularly when studying plasticity, it is of utmost importance to have precise control over the culture environment, as otherwise any changes in behavior cannot be attributed to intrinsic changes of the culture itself, but potentially also to extrinsic properties such as osmolarity, which is directly affected by evaporation, or temperature fluctuations. Both have been shown to have a strong effect on spiking activity by our first two experiments. Another important feature of inkube is its scalability. By combining inkube with PDMS microstructures as shown in this work, a single inkube is capable of stimulating and recording neuronal activity from 60 independent neuronal circuits in parallel at reasonable cost. Finally, inkube is highly autonomous. Experimental paradigms do not require user interaction or can be controlled remotely, once the experiment has been setup, making inkube highly attractive for large scale drug screening experiments. One noteworthy inconvenience of inkube is the time it takes to set up experiments. Depending on the needs, each inkulevel together with its up to 4 liquid pathways needs to be set up, aligned, and washed, which introduces considerable over-head.

Besides using inkube for plasticity research and drug screening, future work will focus on allowing for closed-loop experiments. This requires that the applied environment parameters as well as potential electrical stimuli can be adapted according to the recorded neuronal activity. Combining environmental control with machine learning holds high potential (58). Furthermore, electrical stimulation of a neuronal network in a closed-loop manner has been shown to enable experiments that either react to network changes or are more efficient in investigating the network input-output relationship (59–61).

With the precise temperature control, volume feedback, and automatic medium exchange, the limiting factor for week-long experiments is contamination. A modularization of the electrophysiology board allows for easier assembly and operation in a clean environment such as a laminar flow cabinet. Alternatively, sealing inkulevel airtight or replacing it with a flow cell that is interfaced with filters and the existing fluidic setup has the potential to mitigate risks for contamination. Furthermore, the current version of the valve multiplexers contains a considerable dead volume. Minimizing the dead volume may further reduce the risk of contamination through the fluidic setup. Another possible avenue for mitigation of contamination risks is the use of more persistent antibiotics and antimycotics.

To make inkube more accessible, we are working on adapting it to other MEA layouts and commercial headstages, also featuring CMOS-based high-density MEAs. Furthermore, we are developing a system module that can be integrated into imaging setups to seamlessly combine electrophysiology with calcium imaging, optogenetics, and other imaging approaches.

## Conflicts of interest

There are no conflicts to declare.

## Acknowledgments

This research was supported by ETH Zürich, the Swiss National Science Foundation (SNSF) [project number 182779], the Human Frontiers Science Program (HFSP), and the Swiss Data Science Center (SDSC). S. J. Ihle thanks the Eric and Wendy Schmidt AI in Science Postdoctoral Fellowship (Schmidt Futures program). The authors also thank Peter Littlewood and Narayanan Bobby Kasthuri for their continued support.

## Data and material availability

Manufacturing files and experimental data are available in the ETH Research Collection under https://doi.org/10.3929/ethz-b-000705950. The software is available at https://github.com/maurer-lbb/inkube_software.git.

## CRediT author contributions

**Benedikt Maurer:** Conceptualization, Methodology, Software, Validation, Investigation, Data Curation, Writing - Original Draft, Visualization, Supervision.

**Selina Fassbind:** Methodology, Validation, Investigation.

**Tobias Ruff:** Validation, Investigation.

**Jens Duru:** Validation, Investigation.

**Giusy Spacone:** Methodology, Investigation.

**Theo Rodde:** Methodology, Investigation.

**János Vörös:** Supervision, Conceptualization, Funding acquisition, Project administration.

**Stephan J. Ihle:** Conceptualization, Methodology, Software, Validation, Investigation, Data Curation, Writing - Original Draft, Visualization, Supervision.

## Notes

https://doi.org/10.3929/ethz-b-000705950

https://github.com/maurer-lbb/inkube_software.git

### Competing Interest Statement

The authors have declared no competing interest.

