## Supplementary Information for "Inkube: An all-in-one solution for neuron culturing, electrophysiology, and fluidic exchange"

December 6, 2024

#### A Hardware components

This section describes the hardware components of inkube and contains instructions for the assembly.

##### A.1 Parts list

Table 1: Parts list

| File Name | Description | Linked assembly |
|---|---|---|
| FA5_001_dry_bath | dry bath for cooling | - |
| FA5_002_valve_mux | multiplexer with valves | - |
| FA5_003_pump | syringe pump with stepper motor | - |
| MA5_004_bot_fan | Fan for peltier cooling | - |
| FA5_005_inkuflow | Valve and pump driver | - |
| GA5_006_fpga | FPGA with shield | - |
| GA5_007_power | power management and drivers | - |
| EA5_008_co2 | CO2 valve and connectors | - |
| FA5_009_led | LED status for fluidics valve state | - |
| GA5_010_electronics_fan | fan to cool electronics and drivers | - |
| GA5_011_base | bottom board with MEA holder and ... | - |
| PA5_012_phys | electrophysiology board | - |
| VA5_013_inkulevel | volume sensor | - |
| EA5_014_inkusense | temperature, hum, co2 sensor | - |
| MA5_015_mea_sensor | RTD for MEA temperature | - |
| FA5_016_tube_fitting | fitting to seal the incubation ch... | - |
| EA5_017_humidity | water bath with heater for humidi... | - |
| EA5_018_lid_fan | fan in the lid for better control | - |
| EA5_019_res_heater | resistive reservoir heater | - |
| GD1_020_inkube_pwr_v4 | System power and environment drivers | GA5_007_power |
| GD1_021_inkube_fpga_v4 | FPGA shield | GA5_006_fpga |
| GD1_022_inkube_bot_v4 | MEA base board with temperature c... | GA5_011_base |
| PD1_023_inkube_top_v4 | MEA contacting and electrophysiology | PA5_012_phys |
| FD1_024_inkube_flw_v4 | Valve and pump driver | FA5_005_inkuflow |
| FD1_025_inkube_led_v4 | Valve multiplexer status LEDs | FA5_009_led |
| FD1_026_inkube_valve_v4 | Valve multiplexer connector | FA5_002_valve_mux |
| VD1_027_inkube_lvl_v4 | ESP32 shield | VA5_013_inkulevel |
| GD1_028_inkube_mea_v4 | Holder for MEA | GA5_011_base |
| ED1_029_sensor_board | sensor board | EA5_014_inkusense |
| GA5_030_housing | housing | - |
| FD2_031_housing_0 | fluidics layer | GA5_030_housing |
| GD2_032_housing_1 | electronics layer | GA5_030_housing |
| GD2_033_housing_2 | ventilation layer | GA5_030_housing |
| GD2_034_housing_3 | mea layer | GA5_030_housing |
| FD2_035_housing_4A | side access with tube openings | GA5_030_housing |

| | | |
|---|---|---|
| GD2_036_housing_4B | side access closed | GA5_030_housing |
| GD2_037_housing_5A | lid | GA5_030_housing |
| GD2_038_housing_5B1 | lid with open top | GA5_030_housing |
| GD2_039_housing_5B2 | acrylic glass for lid | GA5_030_housing |
| FA4_040_005_to_002 | 14-pin ribbon cable | - |
| FA4_041_005_to_009 | 20-pin ribbon cable | - |
| FA4_042_006_to_005 | 12-pin ribbon cable | - |
| GA4_043_006_to_007 | 8-pin ribbon cable | - |
| GA4_044_006_to_011 | 12-pin ribbon cable | - |
| GX4_045_007_to_006 | Molex cable (4-pin) Microfit 3.0 | - |
| PX4_046_007_to_012 | Molex cable (4-pin) Microfit 3.0 | - |
| FX4_047_007_to_005 | Molex cable (4-pin) Microfit 3.0 | - |
| GA4_048_007_to_011 | 14-pin Power cable | - |
| GX4_049 | Molex Microfit 3.0 Crimp | GA4_048_007_to_011 |
| GX4_050 | Molex Microfit 3.0 14CKT receptable | GA4_048_007_to_011 |
| PA4_051_006_to_012 | 68-pin ribbon cable | - |
| VA4_052_011_to_013 | 10-pin ribbon cable | - |
| PX4_053 | Ribbon cable (68-pin, 0.025') | PA4_051_006_to_012 |
| PX4_054 | 68-pin connector | PA4_051_006_to_012 |
| GX4_055 | 8-pin connector | GA4_043_006_to_007 |
| VX4_056 | 10-pin connector | VA4_052_011_to_013 |
| FX4_057 | 14-pin connector | FA4_040_005_to_002 |
| FX4_058 | 12-pin connector | FA4_042_006_to_005 |
| GX4_059 | 12-pin connector | GA4_044_006_to_011 |
| FX4_060 | 20-pin connector | FA4_041_005_to_009 |
| GX4_061 | ribbon cable 0.05', 30m | - |
| GX5_062 | cable for Molex | - |
| GX4_063_006_to_PC | USB-A to USB-A, 3.0, 1.8m | - |
| GX4_064_006_to_PC | Patch cable right angle | - |
| GX5_065_power_supply | Power supply 9V | - |
| GX3_066_soc | Zynq FPGA board | GA5_006_fpga |
| VX3_067_esp | ESP32-CAM AI Thinker Camera Module | VA5_013_inkulevel |
| FX3_068_motor_driver | pump driver AD4988 | FA5_005_inkuflow |
| MX5_069_sink | Heat Sink | GA5_011_base |
| MX5_070_epoxy | Araldite Instant Epoxy | GA5_011_base |
| MX5_071_pad | Thermal gap pad | GA5_011_base |
| MX5_072_pt | PT1000 2.0x4.0 B | MA5_015_mea_sensor |
| MX6_073_seal | PTFE heat shrink tube | MA5_015_mea_sensor |
| EX5_074_valve | CO2 Valve | EA5_008_co2 |
| EX6_075_tube | CO2 tubing, 50m | EA5_008_co2 |
| EX5_076_con | CO2 wall connector | EA5_008_co2 |
| FX5_077_valve | memetis microvalves | FA5_002_valve_mux |
| FX5_078_valve_ov | memetis microvalves negative pres... | FA5_002_valve_mux |
| EX5_079_hum_heater | Cartridge Heater Kit | EA5_017_humidity |
| MX5_080_bot_fan | Axial Fan peltiers | MA5_004_bot_fan |
| GX5_081_el_fan | Axial Fan electronics | GA5_010_electronics_fan |
| EX5_082_lid_fan | Axial Fan lid | EA5_018_lid_fan |
| FX5_083_sink_bath | heat sink for dry bath | FA5_001_dry_bath |
| FX5_084_pad_sticker | Thermal Gap Pad 0.13mm thickness | FA5_001_dry_bath |
| ED2_127_bath | bath for humidity | GA5_030_housing |
| PD2_085_spacer_r | spacer between base and e-phys. B... | PA5_012_phys |
| PD2_086_spacer_l | spacer between base and e-phys. B... | PA5_012_phys |
| PD2_087_spacer_r_flex | spacer between base and e-phys. B... | PA5_012_phys |
| PD2_088_spacer_l_flex | spacer between base and e-phys. B... | PA5_012_phys |
| PD2_089_spacer_bot | spacer between base and e-phys. B... | PA5_012_phys |
| PD2_090_spacer_bot_flex | spacer between base and e-phys. B... | PA5_012_phys |
| PX5_092_spring_contact | spring contact for MEA | PA5_012_phys |
| PD2_091_spacer_cover | spacer between base and e-phys. B... | PA5_012_phys |

| | | |
|---|---|---|
| VD2_094_bending_helper | to bend syringe needles | VA5_013_inkulevel |
| VX5_095_clamp | Clamp Syringe | VA5_013_inkulevel |
| VX5_096_laser | Laser module 650 nm, 6-12 VDC | VA5_013_inkulevel |
| FX5_097_needle | Blunt needle Sterican® 18G / 1,2 ... | VA5_013_inkulevel |
| FX5_098_syringe | Syringe 1mL | FA5_003_pump |
| FD2_099_pump_sledge | Sledge for syringe pump | FA5_003_pump |
| FD2_100_pump_back | Back part syringe pump | FA5_003_pump |
| FD2_101_pump_front | Front part syringe pump | FA5_003_pump |
| FD2_102_pump_base | Base plate syringe pump | FA5_003_pump |
| MX5_103_peltier | TEC device | GA5_011_base |
| FX5_104_peltier | TEC device | FA5_001_dry_bath |
| FX5_105_luer | Luer Fingertight | - |
| FX5_106_luer_f | Luer female | - |
| FX5_107_luer_m | Luer male | - |
| FX6_108_tube | PTFE Liquid tubing ID 1mm, x1m | - |
| PX5_109_tape | Aluminum adhesive tape for shielding | GA5_030_housing |
| FD2_110_valve_splitter | multiplexer with valves print | FA5_002_valve_mux |
| FX5_111_motor | Mini T6 LeadScrew Linear Motion S... | FA5_003_pump |
| FD2_112_holder | syringe holder counter | FA5_003_pump |
| FX5_113_switch | safety switch, Micro Switch, DG, ... | FA5_003_pump |
| FX5_114_bearing | 6 mm linear bearings | FA5_003_pump |
| FX5_115_shaft | 6 mm precision shafts | FA5_003_pump |
| FD2_116_dry_bath_back | dry bath | FA5_001_dry_bath |
| FD2_117_dry_bath_cap | dry bath | FA5_001_dry_bath |
| EX5_118_adapter | CO2 tube adapter | EA5_008_co2 |
| EX5_119_angle | CO2 tube angle | EA5_008_co2 |
| GX5_120_feet | rubber feet | GA5_030_housing |
| VX5_121_spring | spring for inkulevel pinhole | VA5_013_inkulevel |
| VD2_122_inkulevel_body | main part for volume sensor | VA5_013_inkulevel |
| VD2_123_inkulevel_pinhole | height adjustable pinhole for laser | VA5_013_inkulevel |
| VD2_124_inkulevel_top | counter for pinhole | VA5_013_inkulevel |
| PX5_125_mea | MCS MEA 60MEA500/10iR-Ti -gr | - |
| EX6_126_seal_hum | wrapping for humidity heater | EA5_017_humidity |
| PD2_127_mounting_helper | to mount spring contacts to e-phy... | PX4_046_007_to_012 |
| EX5_128_heater_sink | 4 Ohm resistor for heating with h... | EA5_019_res_heater |
| EX5_129_heater_tht | 4 Ohm resistor high power | EA5_019_res_heater |
| FD2_130_dry_bath_cover | dry bath cover to prevent condens... | FA5_001_dry_bath |
| FD2_131_tube_lid | cap for falcon tube with hole for... | FA5_001_dry_bath |
| GD2_132_lid_hook | holder for lid when open | GA5_030_housing |
| FD2_133_tube_fitting | fitting to seal the incubation ch... | FA5_016_tube_fitting |
| MX6_134_seal_small | PTFE heat shrink tube | MA5_015_mea_sensor |

The partslist lists all components required to build the inkube hardware. The filename serves as a unique identifier. The first letter indicates the functionality, i.e. for which feature of inkube the parts are required. The encoding is as follows:

- G (**G**eneral), these are essential components that are always required
- P (Electro**p**hysiology), required to record and stimulate neuronal cultures
- E (**E**nvironment control), required for temperature, humidity, and $CO_2$ control in the reservoir
- F (**F**luidics), required to pump and retrieve liquid into and from the MEA
- V (**V**olume feedback), required to obtain feedback about the volume in the MEA
- M (**M**EA Temperature control), required to individually control the medium temperature inside the MEA

The second letter indicates whether this is an external component to be bought (X), a design file for a CAD or PCB that is provided in the supplementary data for this paper (D), or an assembly of components (A). The third digit describes the component category listed below:

1 PCB

2 CAD

3 Electronic Devices

4 Cable

5 Component

6 Tube

A more detailed list containing quantity, suppliers, and cost can be found in the Supplementary Data (01_Hardware/01_BOM_syste[illegible]
The design files of the PCBs can be found in the Supplementary Data at `01_Hardware/02_PCB` and the design files for the CAD `01_Hardware/03_CAD`. The parts lists for assembly of the PCBs can be found in the same folder as the PCB design `BOM_PCBs.ods`.

#### A.2 Assembly overview

This section contains an overview of how the system is arranged with all components marked as assemblies. The first figure shows the CAD model in a side view. Next, the housing is depicted in top view from lowest layer (fluidics) to topmost layer (incubation) with annotations where to place other assemblies and cables connecting these. All parts are marked with the unique identifiers from the parts list. Finally, the same layers are shown with annotations for screws.

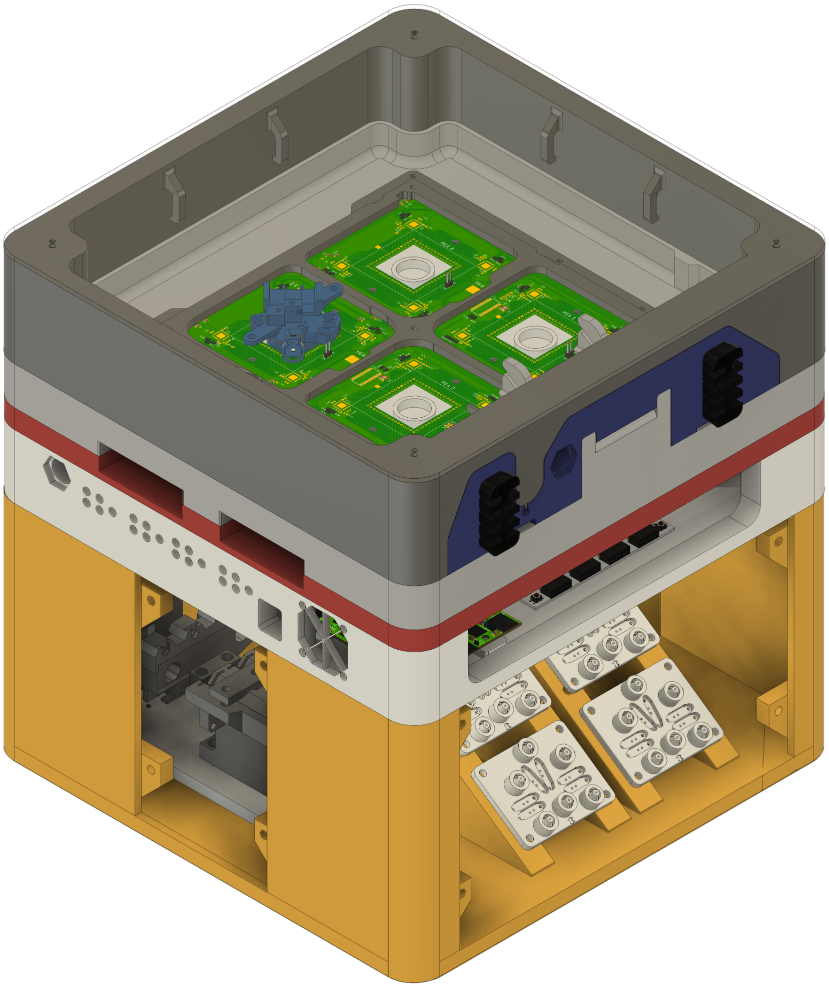

Figure S1: Side view of inkube layers in the CAD model

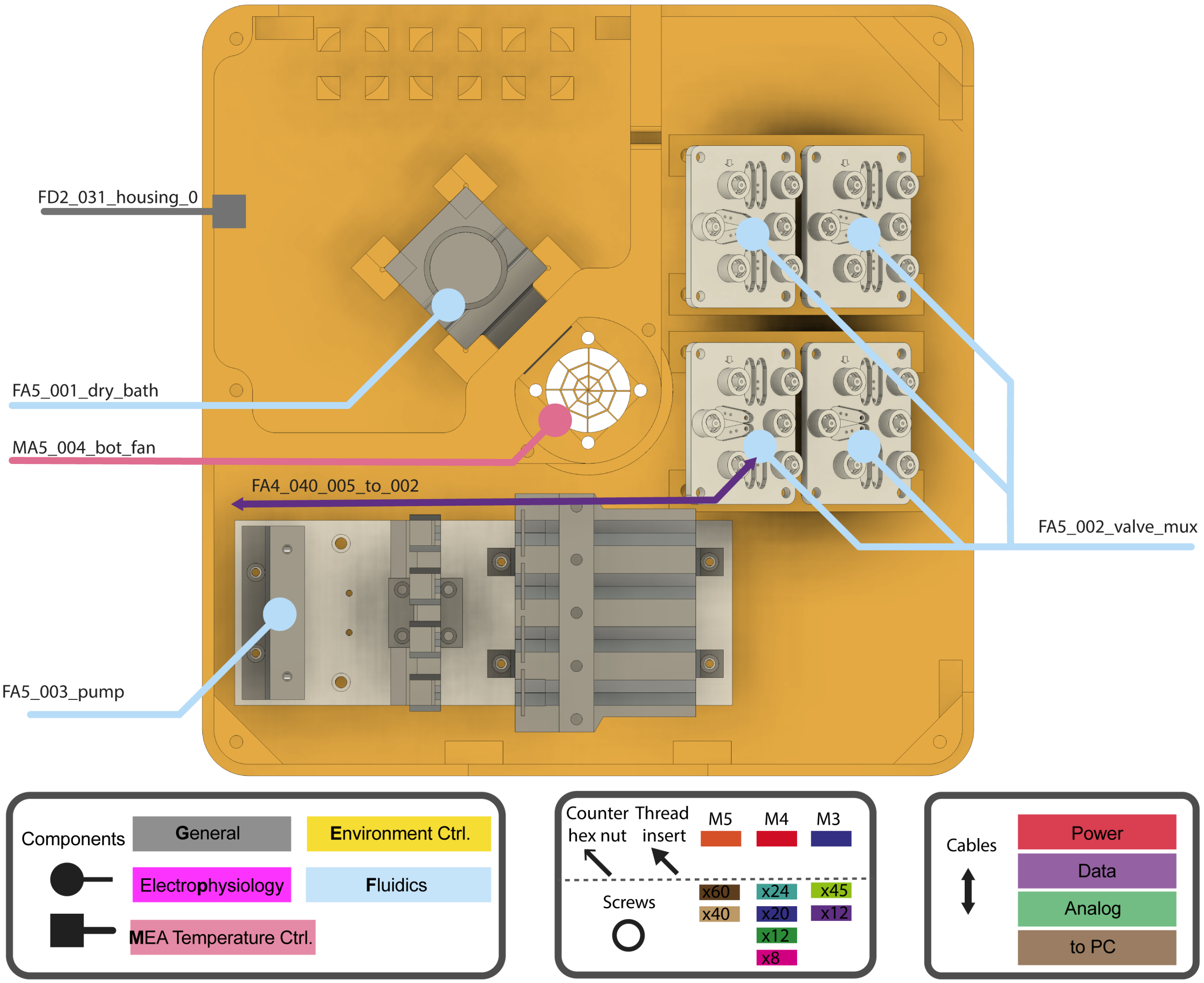


Figure S2: Fluidics layer (Layer 0) overview with components and cables

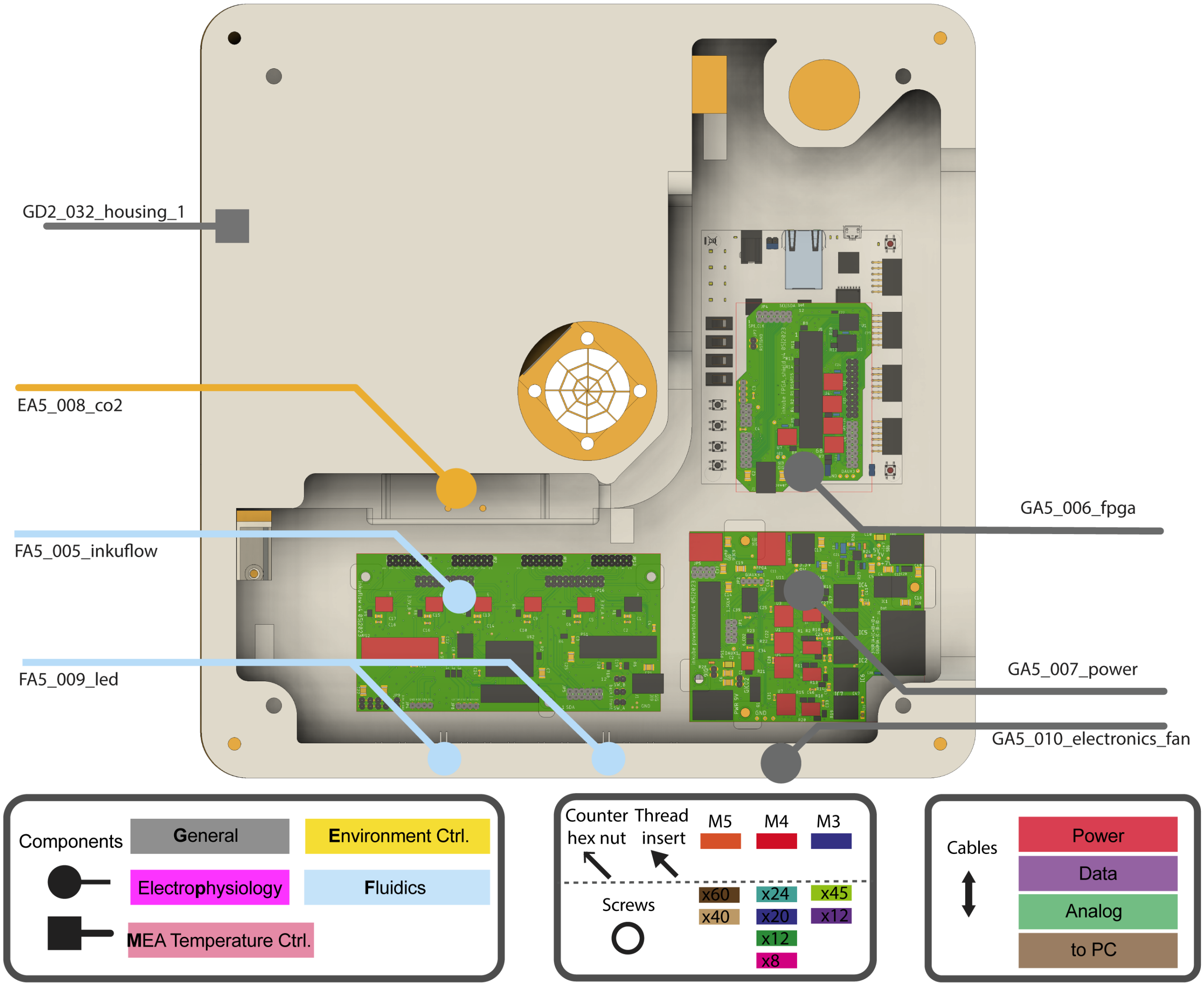


Figure S3: Electronics layer (Layer 1) overview with components

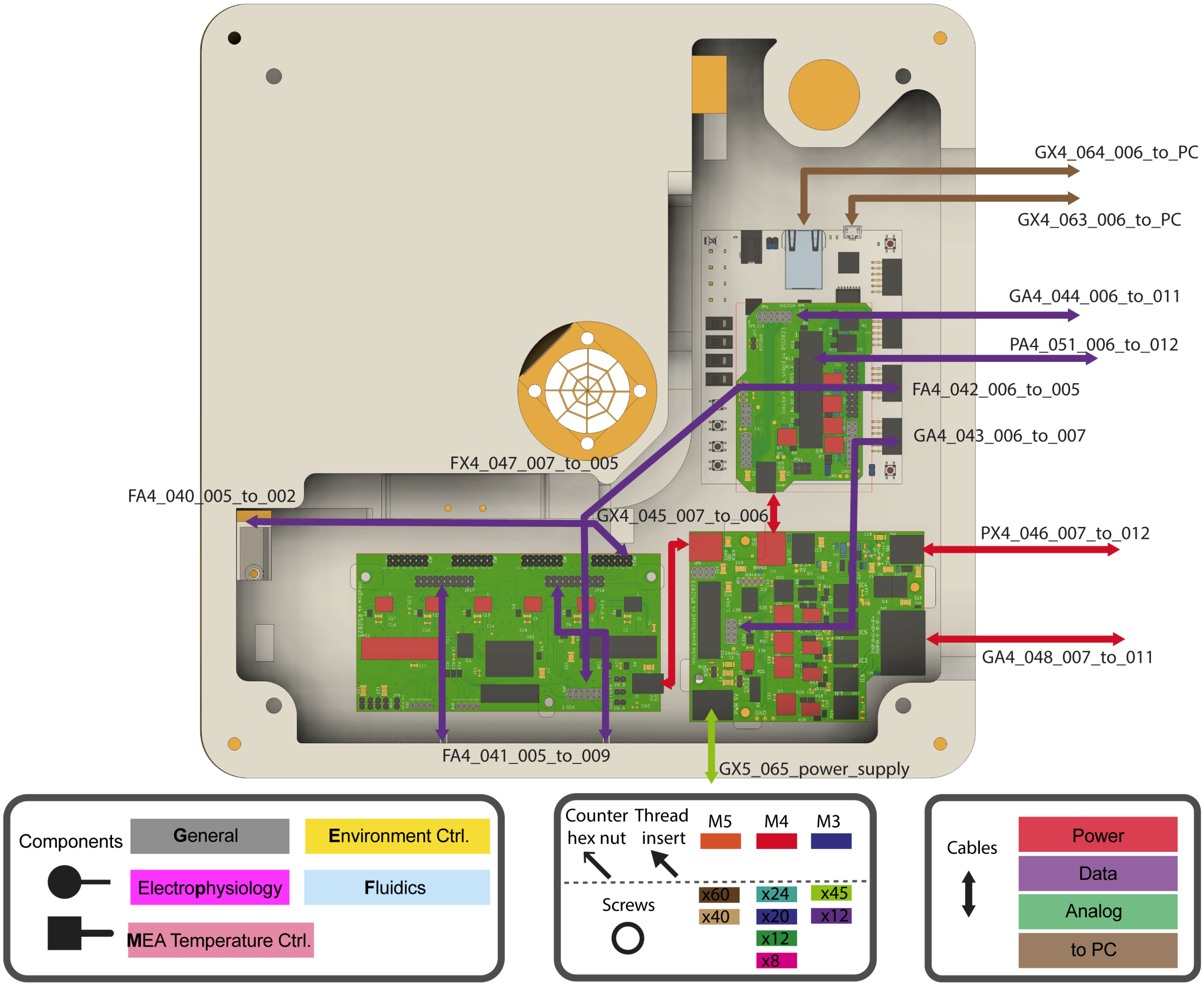


Figure S4: Electronics layer (Layer 1) overview with cables

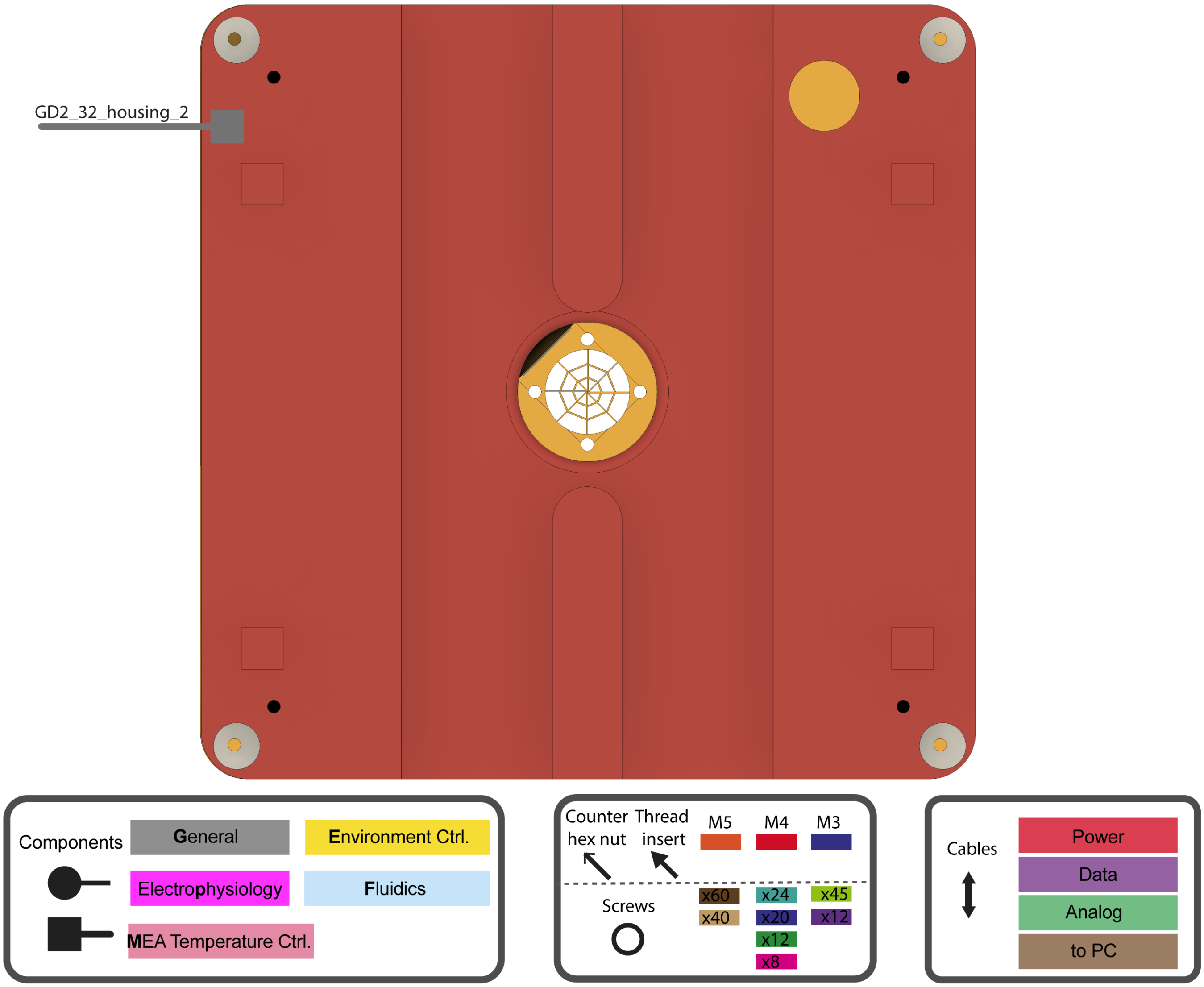


Figure S5: Ventilation layer (Layer 2) overview with components

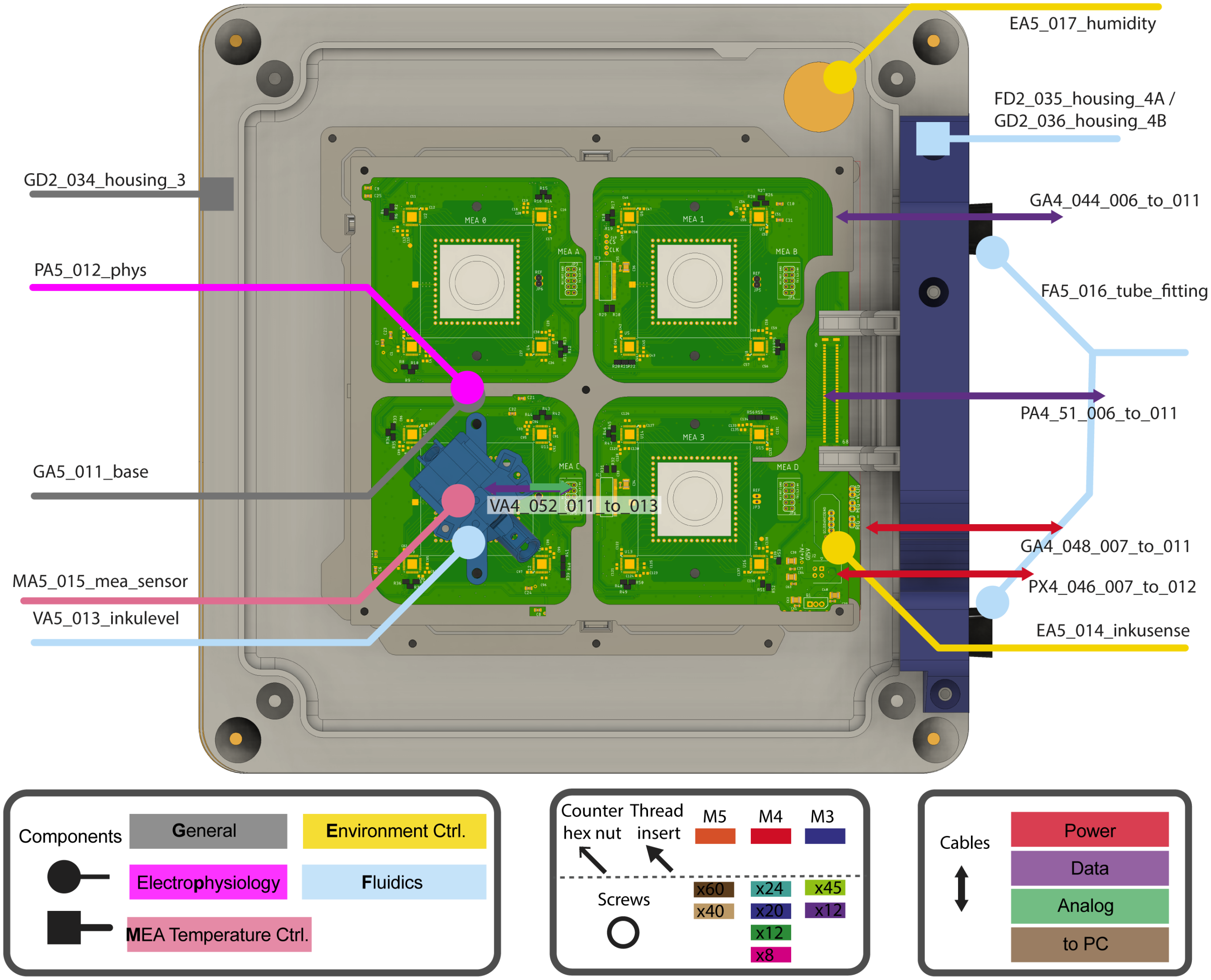


Figure S6: Incubation layer (Layer 3/4) overview with components and cables

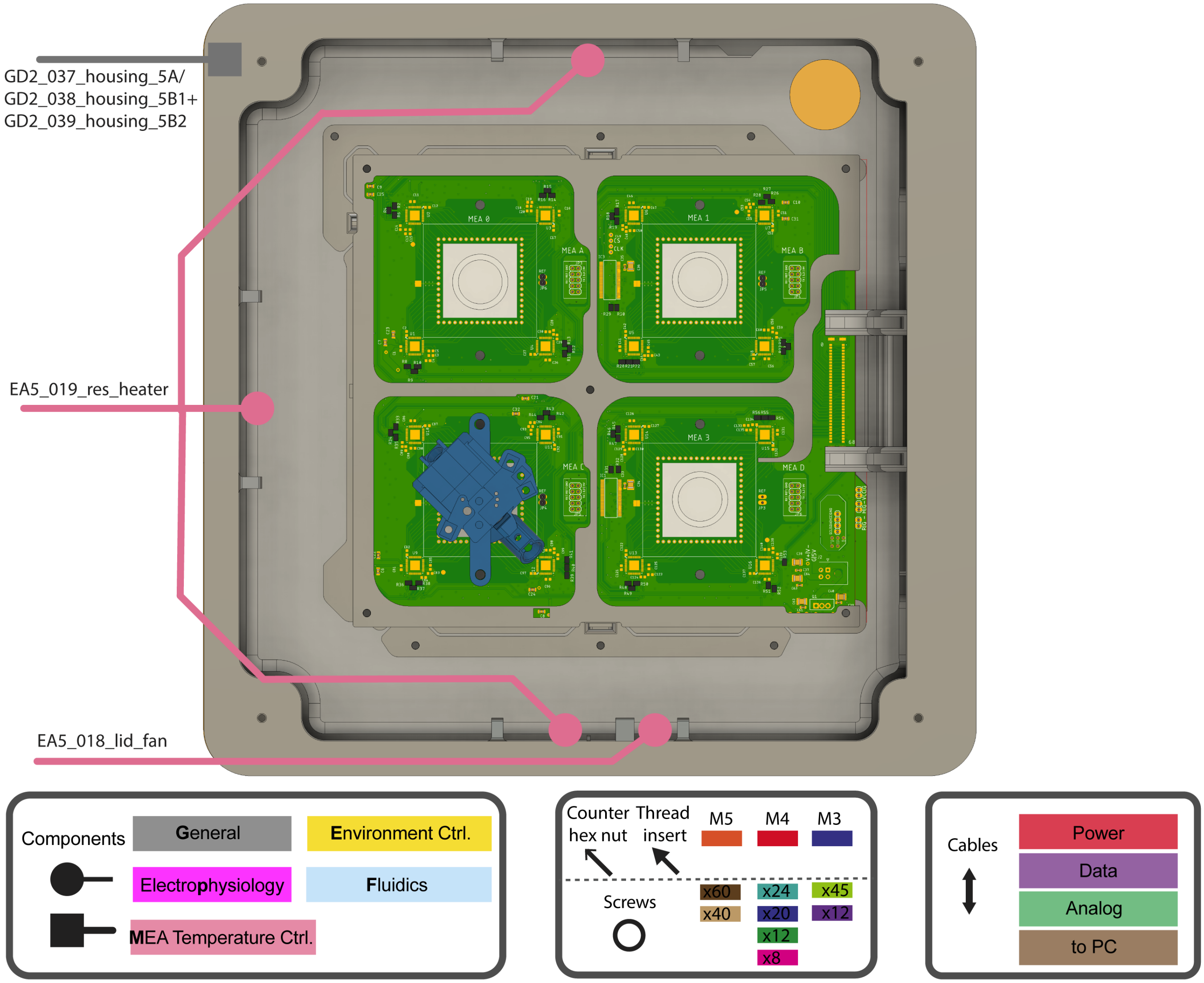


Figure S7: Lid overview with components

Next, the layers are shown with locations for hex nuts, thread inserts in the housing, and screws to attach assemblies to the housing.

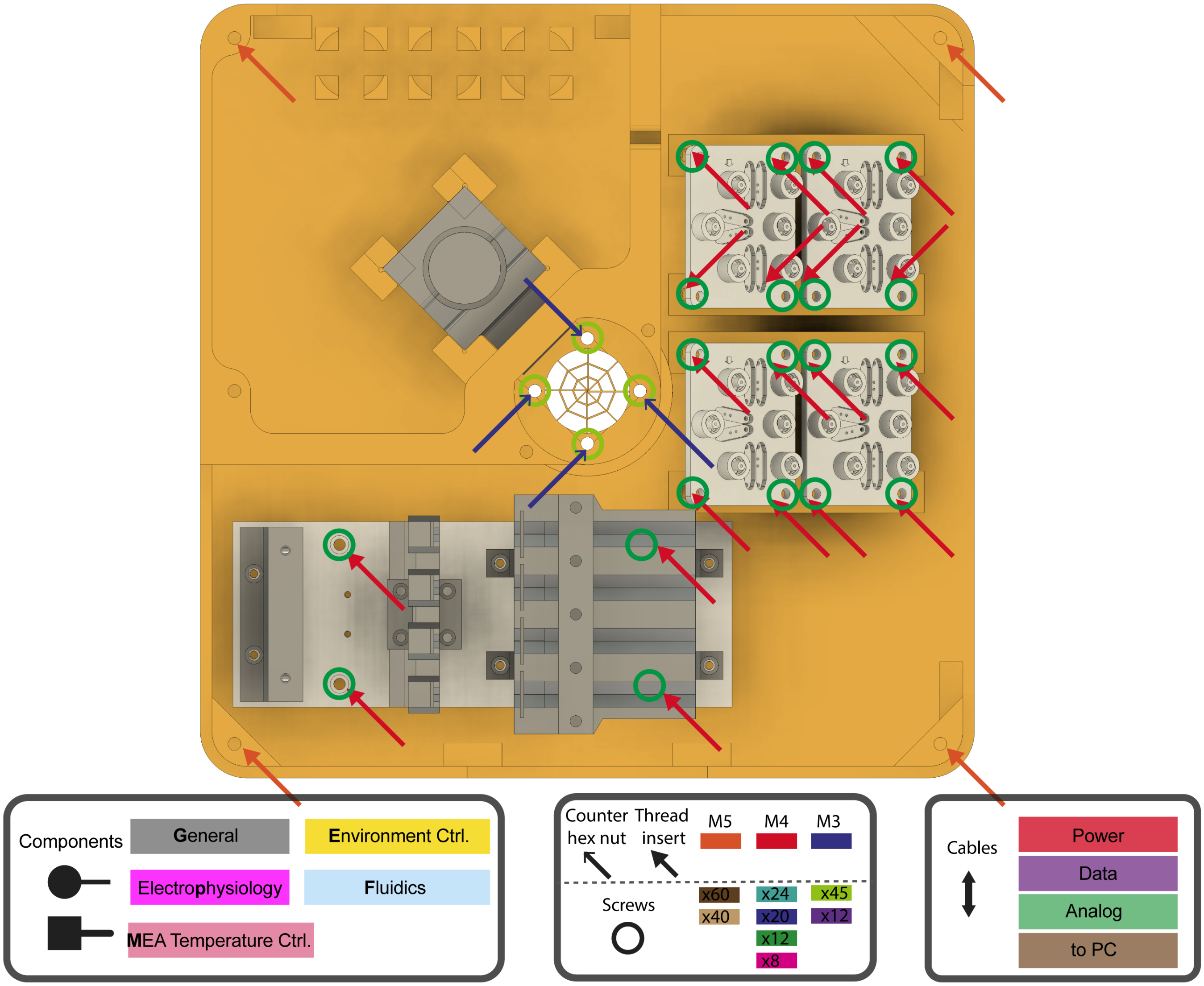


Figure S8: Fluidics layer (Layer 0) locations for screws

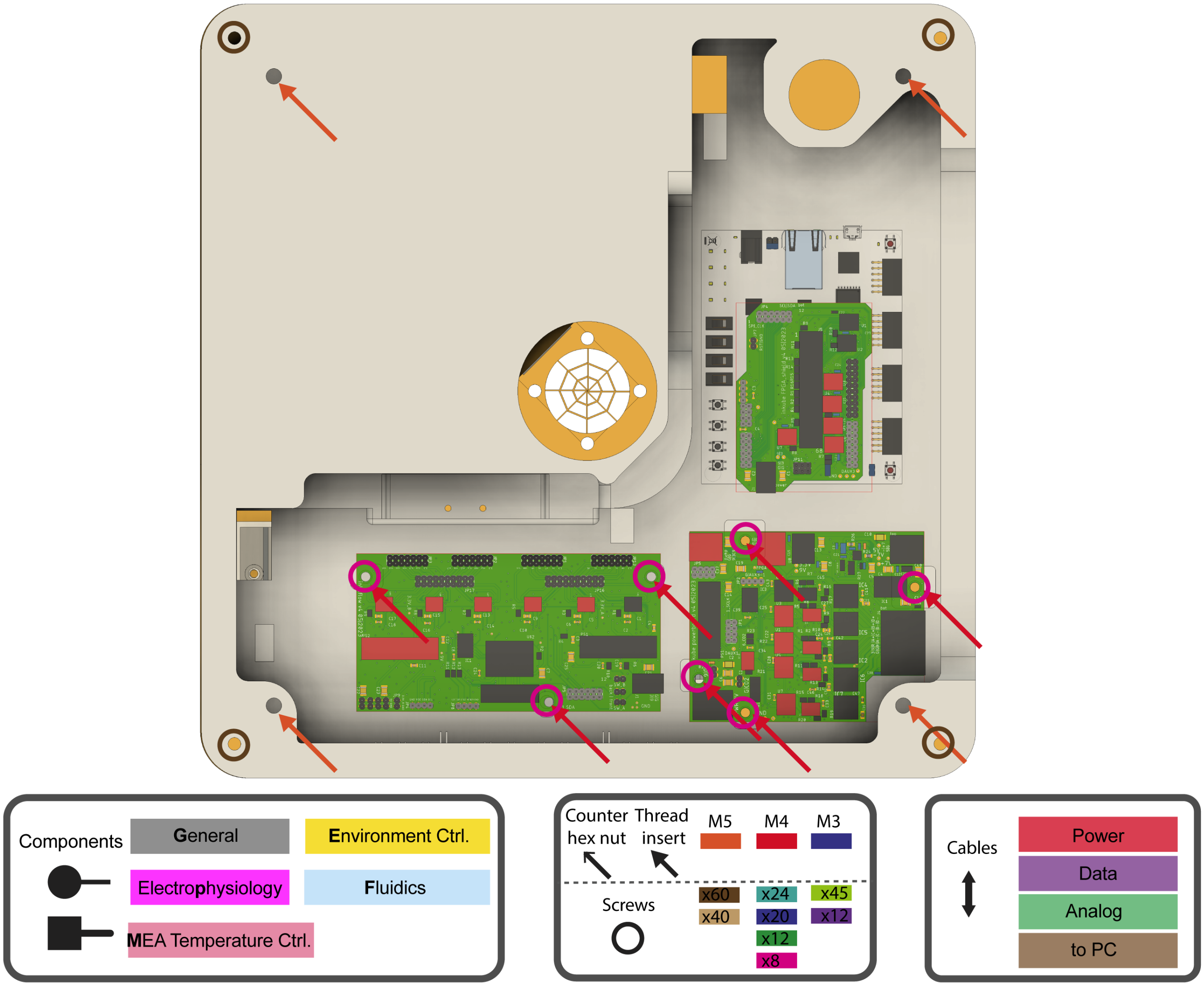


Figure S9: Electronics layer (Layer 1) locations for screws

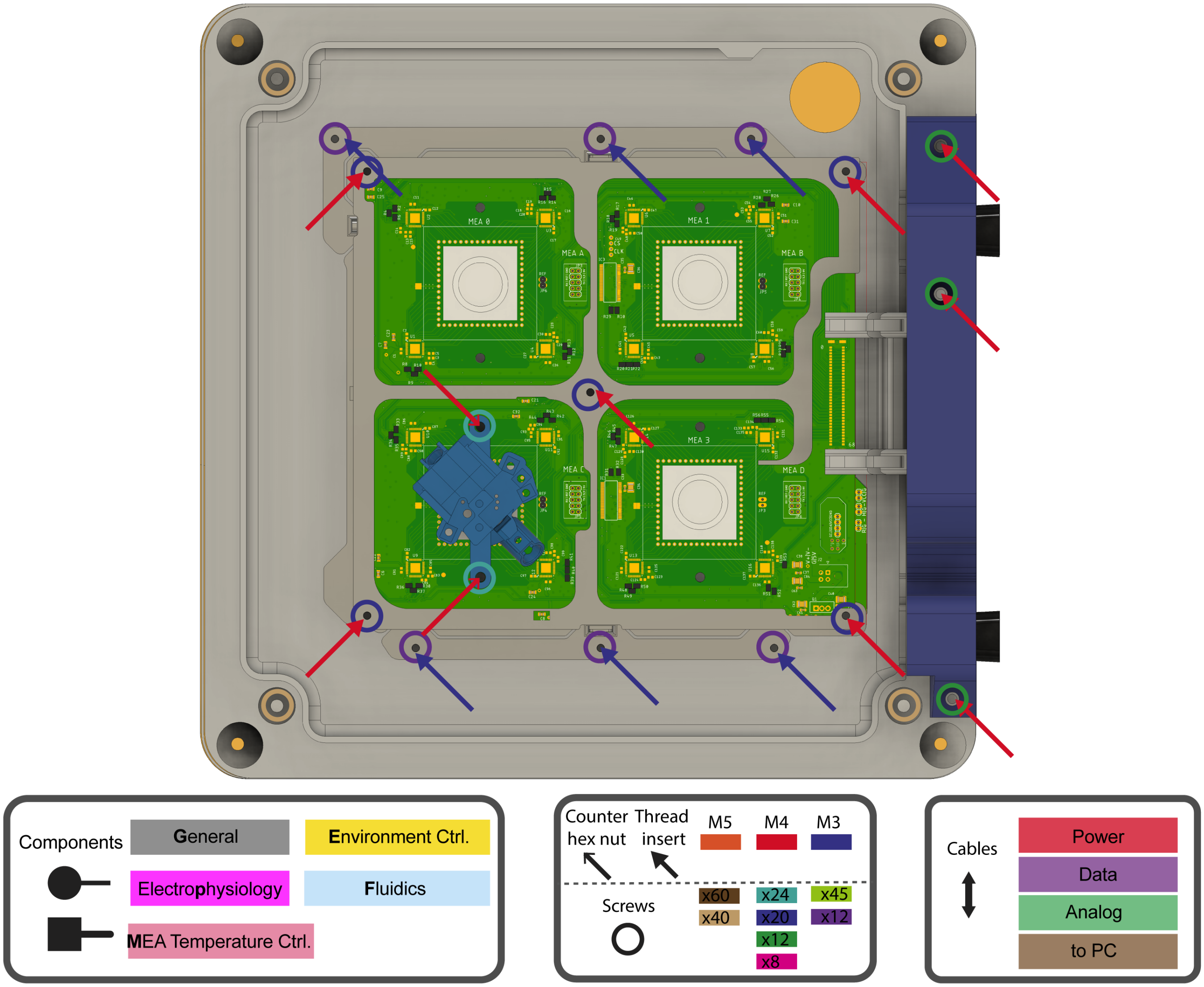


Figure S10: Incubation layer (Layer 3/4) locations for screws

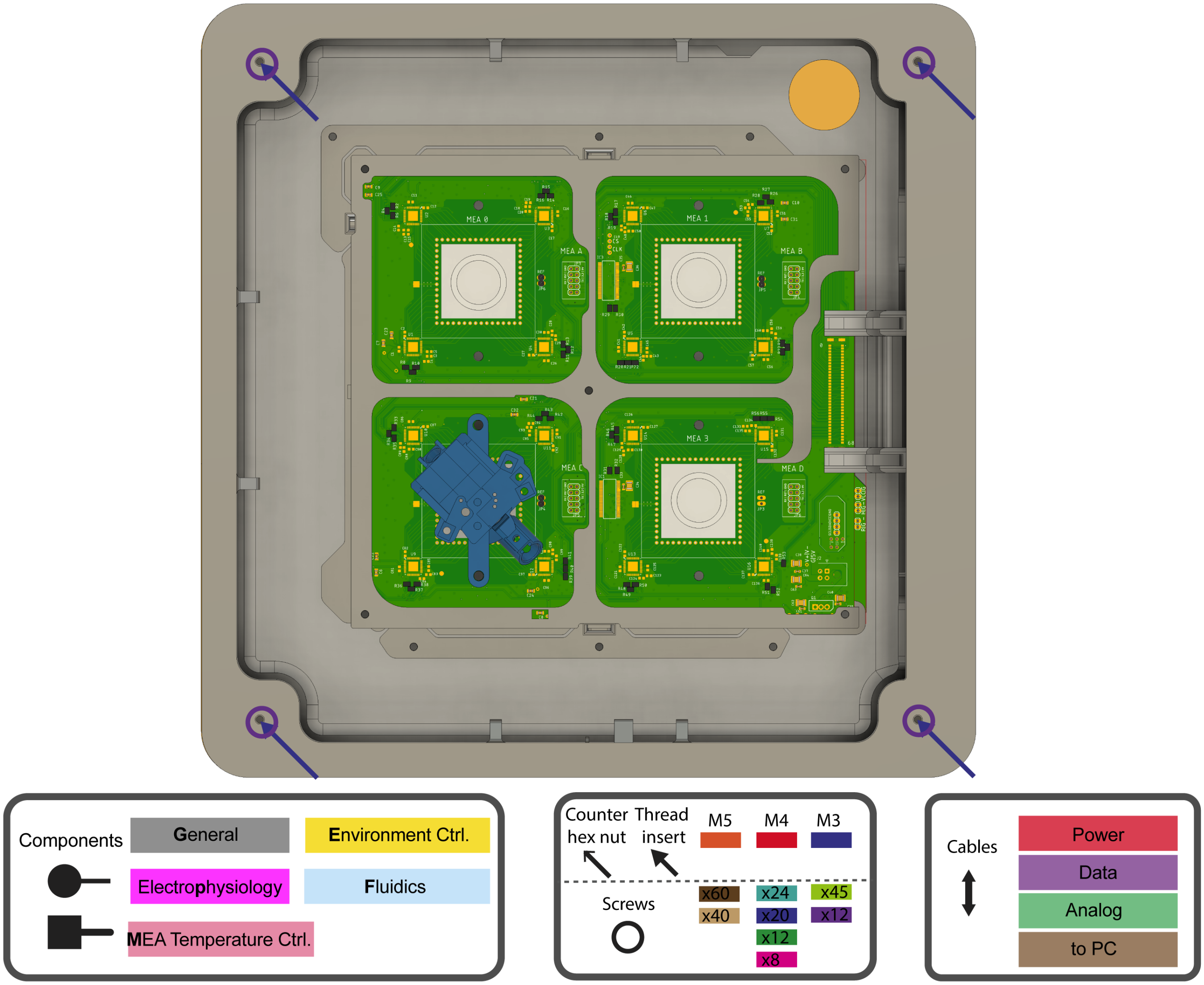


Figure S11: Lid locations for screws

#### A.3 PCBs

In this section all PCB designs are shown. All boards are provided in version 4 of inkube. The unique identifier used in the parts list can be seen in the figure caption.

First the boards required for the general functionality are provided.

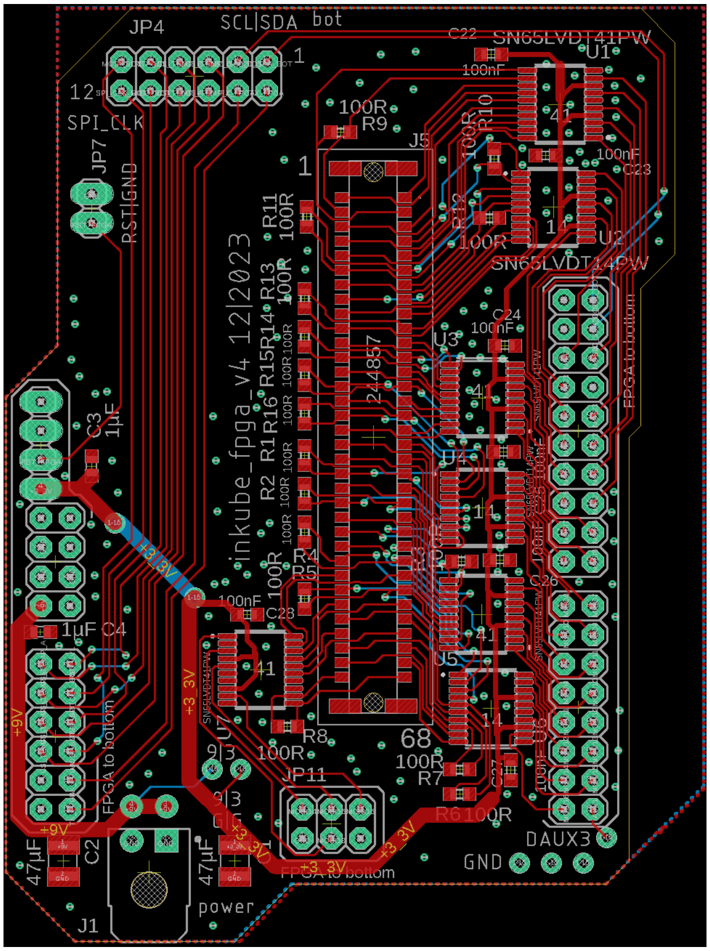


Figure S12: GD1_021_inkube_fpga_v4

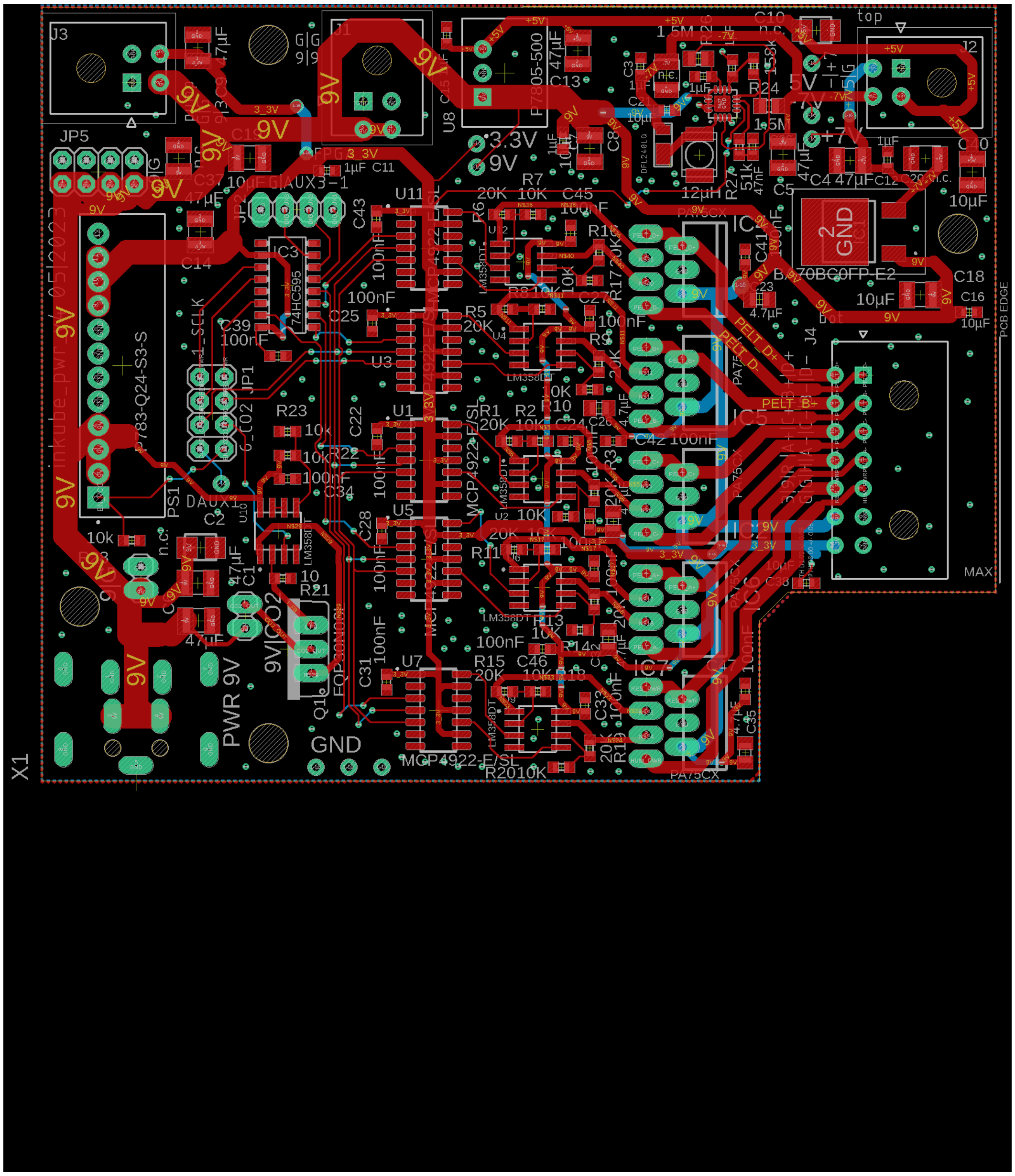

Figure S13: GD1_020_inkube_pwr_v4

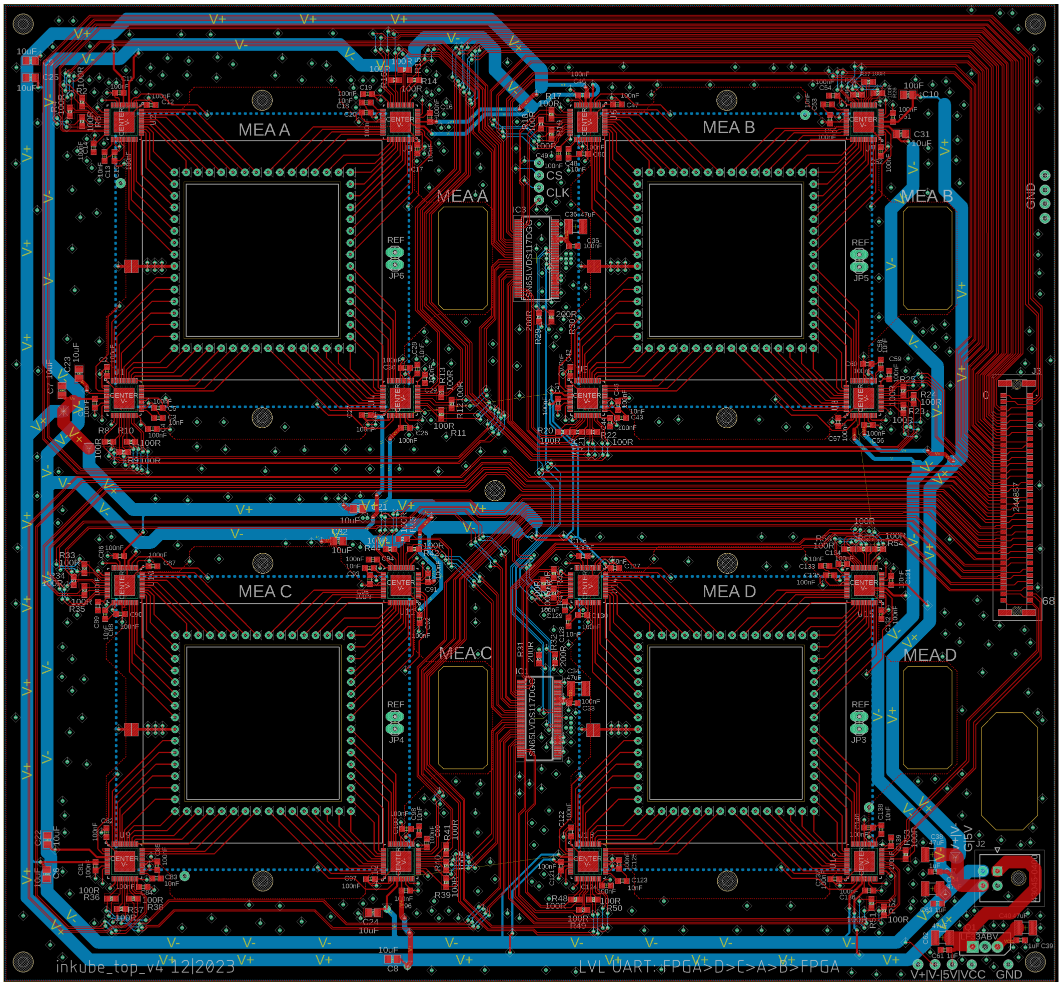


Figure S14: PD1_023_inkube_top_v4

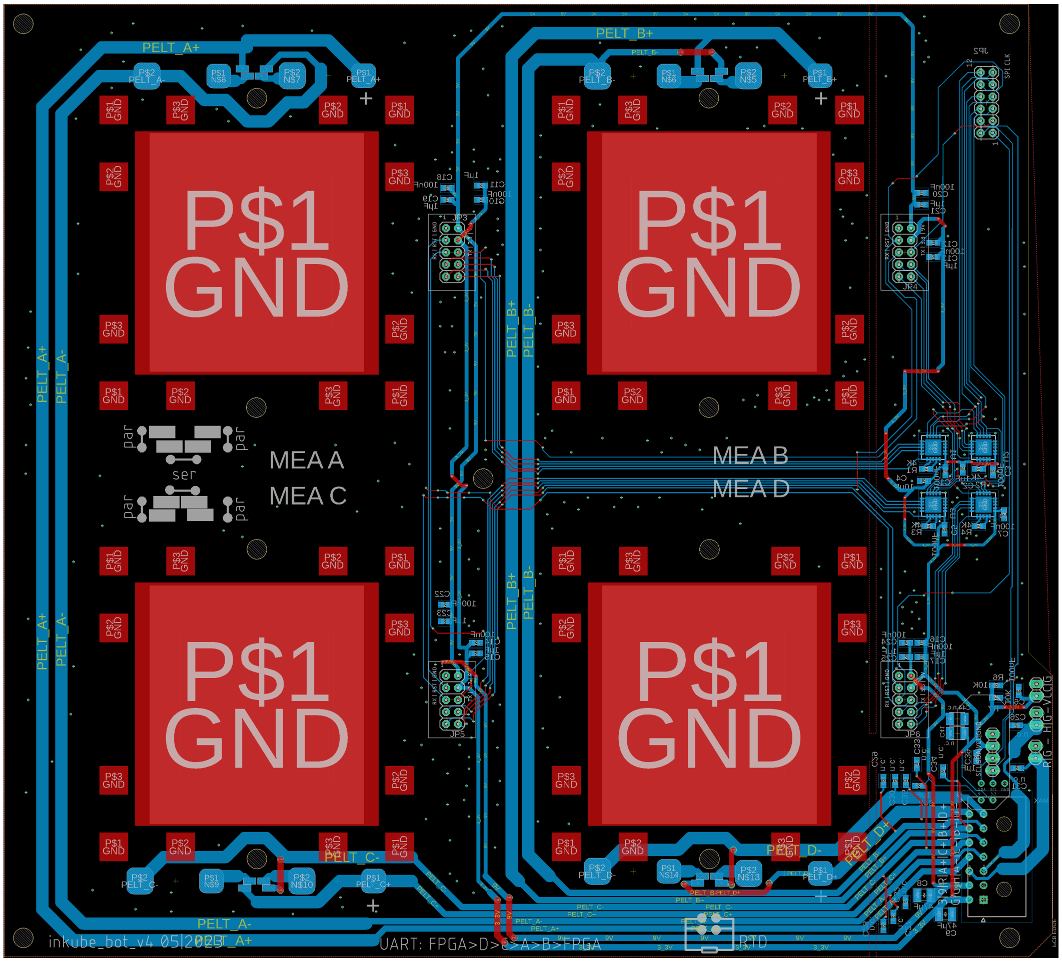


Figure S15: GD1_022_inkube_bot_v4

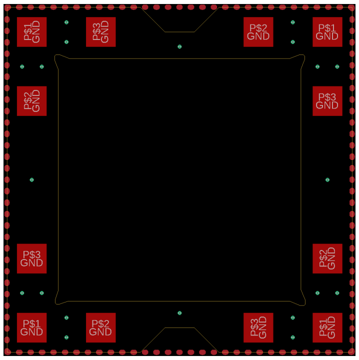

Figure S16: GD1_028_inkube_mea_v4

Here the boards for the fluidic system are listed.

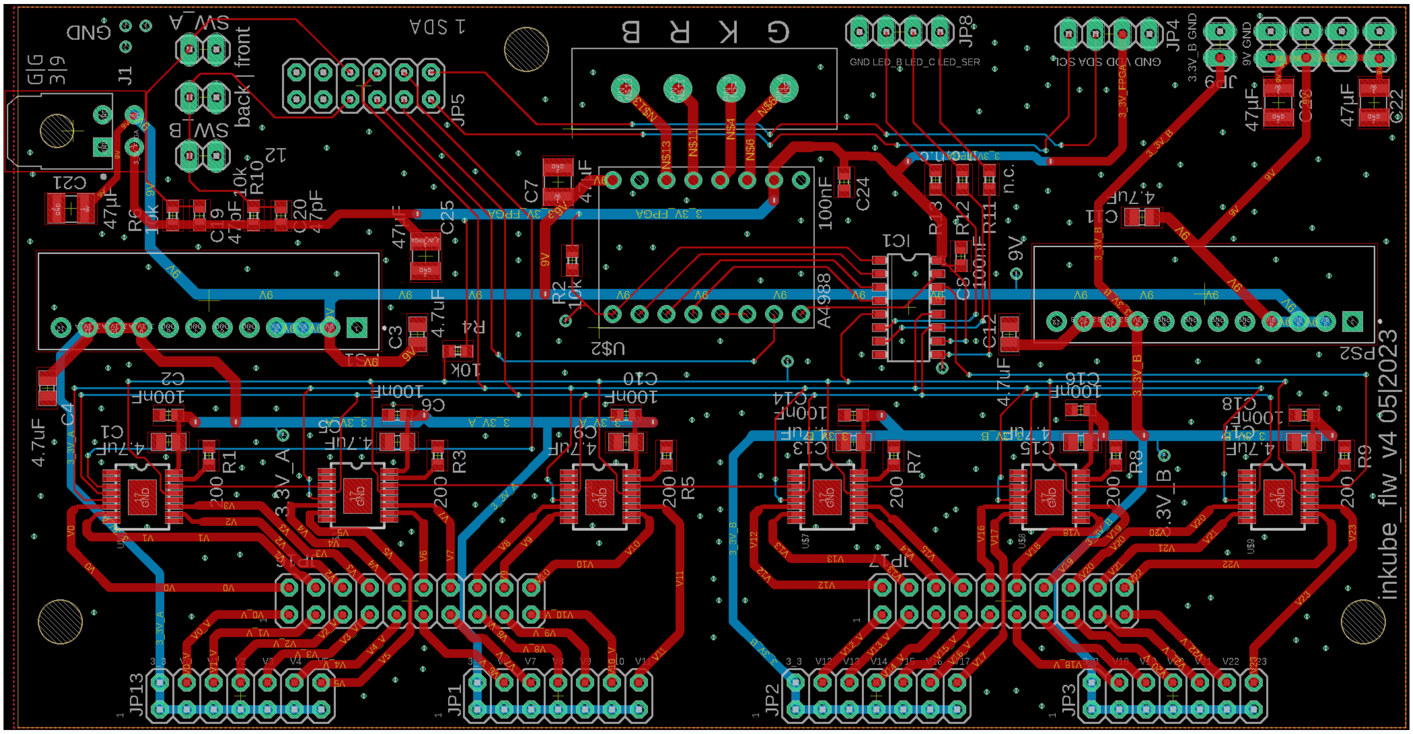


Figure S17: FD1_024_inkube_flw_v4

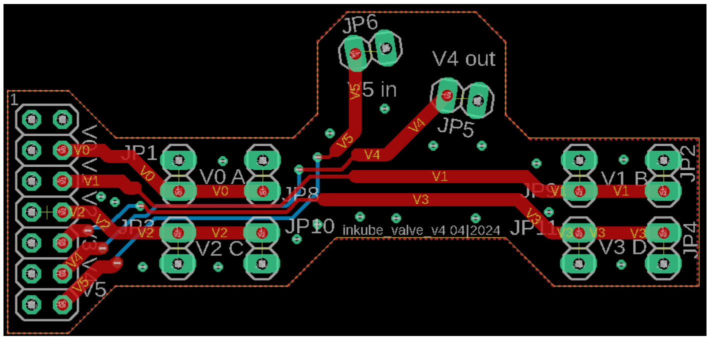


Figure S18: FD1_026_inkube_valve_v4

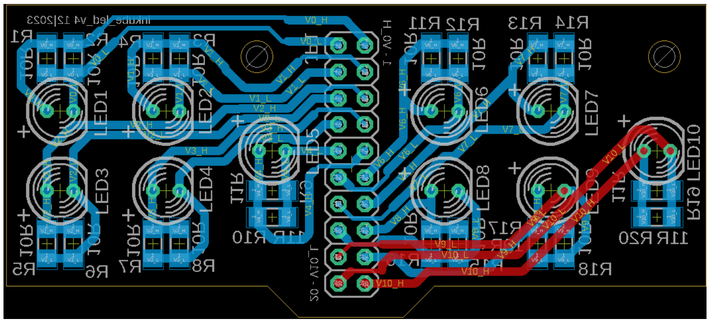


Figure S19: FD1_025_inkube_led_v4

Inkulevel is the only additional board required for volume feedback.

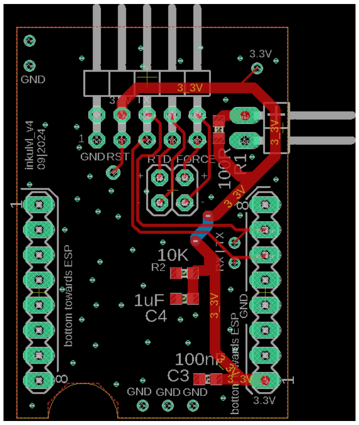


Figure S20: VD1_027_inkube_lvl_v4

#### A.4 Housing

The assembly instructions are provided here for the general components, the housing, and the SoC shield.

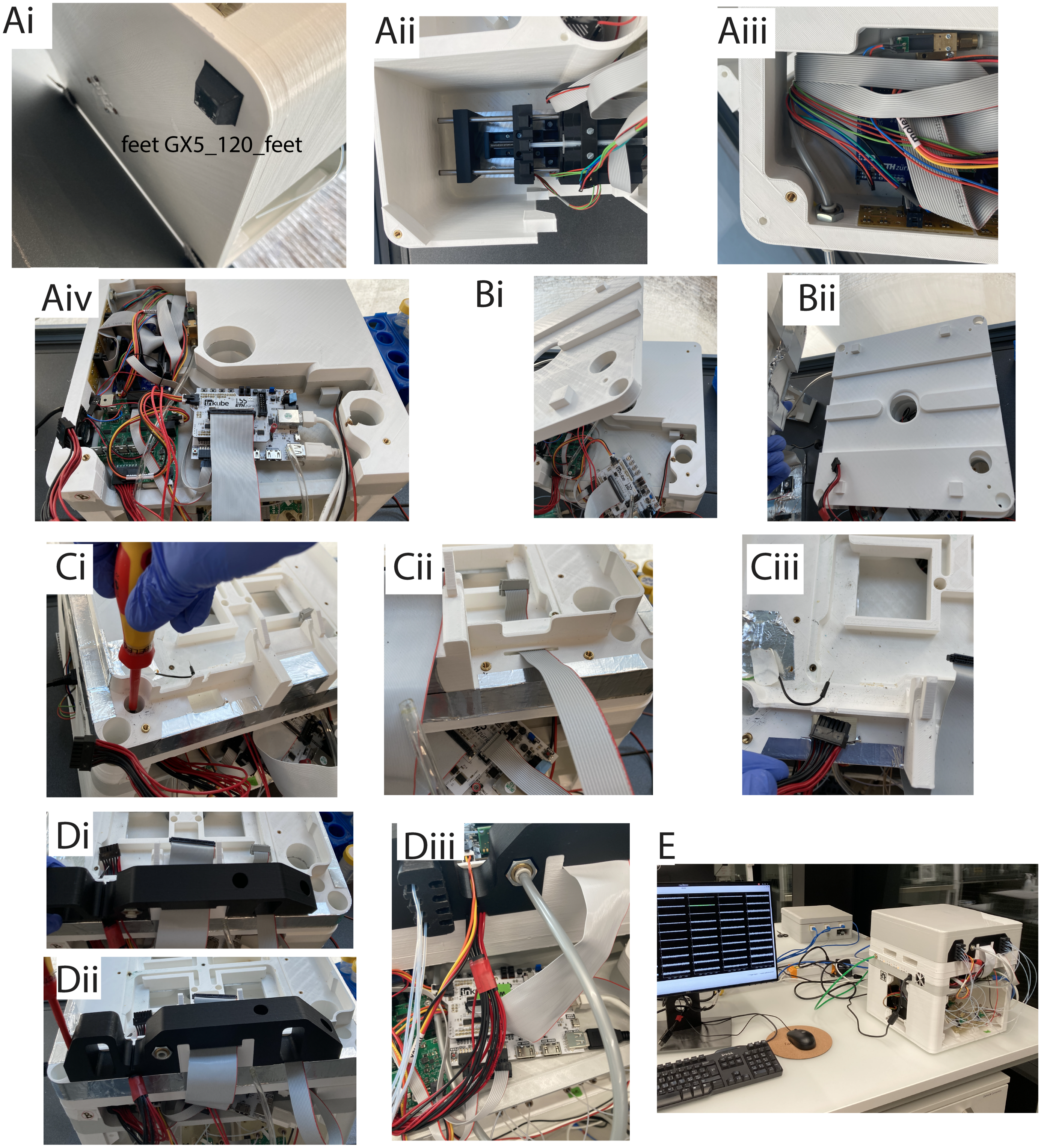


Figure S21: Housing assembly

To assemble the housing, prepare all layers with thread inserts and screws as shown in Fig. S8 to Fig. S11. The order in which to assemble the parts is listed here.

- Attach the feet as shown in Fig. S21Ai
- Assemble the fan as described in Fig. S25
- Assemble the dry bath as described in Fig. S33
- Assemble the pump as described in Fig. S28
- Place the pump as shown in Fig. S2 and Fig. S21Aii
- Mount the electronics layer and fix it with the outer screws
- Place the power board
- Place the inkuflow board
- Mount the electronics fan as described in Fig. S26
- Mount the $CO_2$ valve as described in Fig. S37
- Lead the cables for pump, dry bath, and fan up from the fluidics layer
- Attach the cables for the valve multiplexers and lead them down as shown in Fig. S21Aiii
- Attach all cables connecting the boards within the electronics layer
- Attach the cables to the host PC
- Attach the 2 power and 2 data cables leading up to the incubation layer as shown in Fig. S21Aiv
- Mount the ventilation layer as shown in Fig. S21Bi and Bii
- Mount the incubation layer and fix it with the inner screws as shown in Fig. S21Ci
- Lead the data and power cables for the bottom board through the gaps in the layer as shown in Fig. S21Cii and Ciii, respectively
- Lead the data cable for the top board through the gap
- Mount the incubation layer wall as shown in Fig. S21Di and Dii on top of the cable connections with the 3 inside screws
- Mount the base board and spacers as described in Fig. S23
- Mount the electrophysiology board as shown in Fig. S27
- Mount sensor board as shown in Fig. S23C
- Mount the $CO_2$ tube and the top board power cable as shown in Fig. S21Diii
- Mount the bottom layer valve multiplexers as shown in Fig. S2
- Mount the tube fittings as shown in Fig. S34
- Mount the lid fan as described in Fig. S39

To start an experiment complete these final steps:

- Remove the top board
- Place the MEAs as shown inFig. S27Di
- Mount the top board as shown in Fig. S27Dii and Diii
- Mount the inkulevel as shown in Fig. S35
- Connect the lid cables as shown in Fig. S39B
- Mount the lid (Fig. S21E)

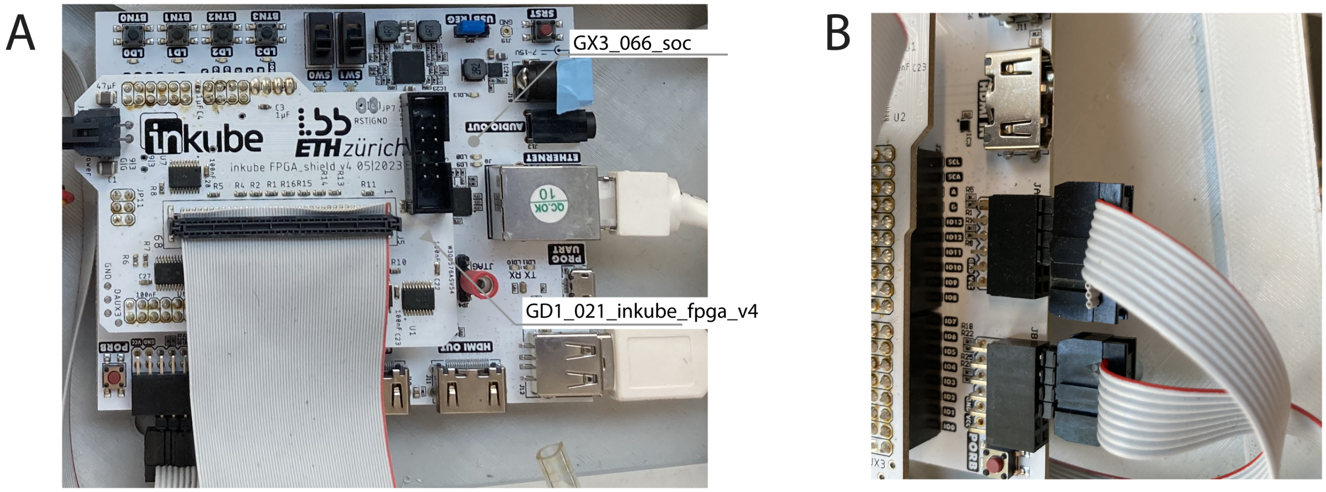


Figure S22: **A** Mount the shield and insert the PC cables, the data cable, and the power board cable **B** Insert the data cables for the power board and inkuflow. Pin 1 of the cables always goes to the top right pin of JA and JB.

#### A.5 MEA Temperature Control

In this section the components for temperature control on the MEAs are listed.

##### A.5.1 Peltier mounting

The thermoelectric Peltier elements are mounted on the bottom board. The steps are shown in Fig. S23A.

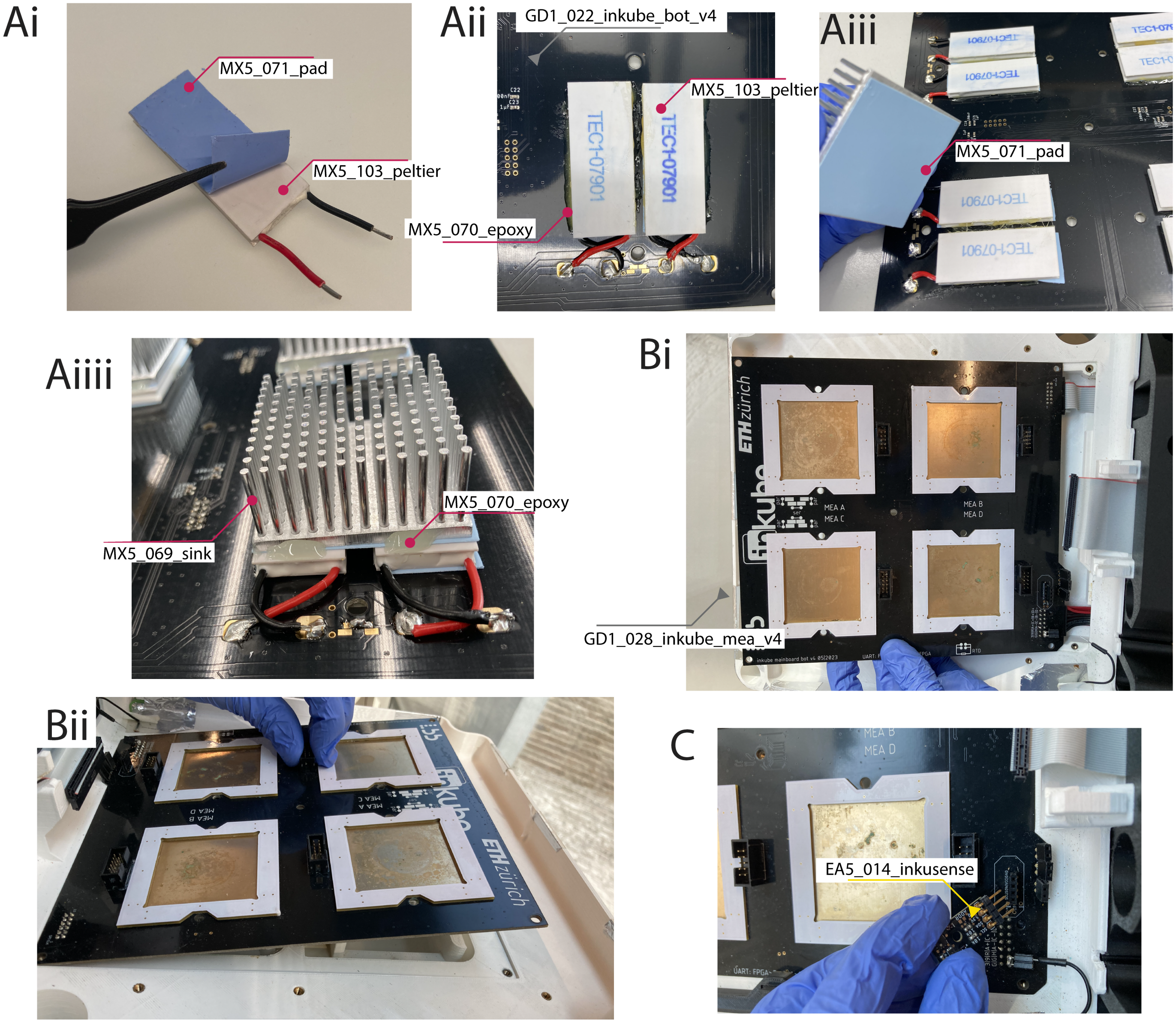


Figure S23: **A**i) cut the cables to about 40 mm. Attach the thermal gap pad to the not-labeled side of the Peltier element. ii) Place Peltier elements on the markings of the bottom board. When the writing points up, connect the red cable to the pad marked with '+' and the second device with similar polarity. For standard operation, shorten the two closer pads horizontally as shown on the bottom board print to use the thermoelectric devices in serial connection. Use the epoxy glue to fix them. iii) mount the heat sink with another piece of the gap pad. iv) Use the epoxy for fixation. **B**i) connect the data and power cable. ii) Slide the board in place from the side opposite of the cable connections. **C** Connect sensor board.

##### A.5.2 RTD assembly

The PT1000 temperature sensor is wrapped in a biocompatible tubing and connected in a 4-point measurement. The steps are shown in Fig. S24.

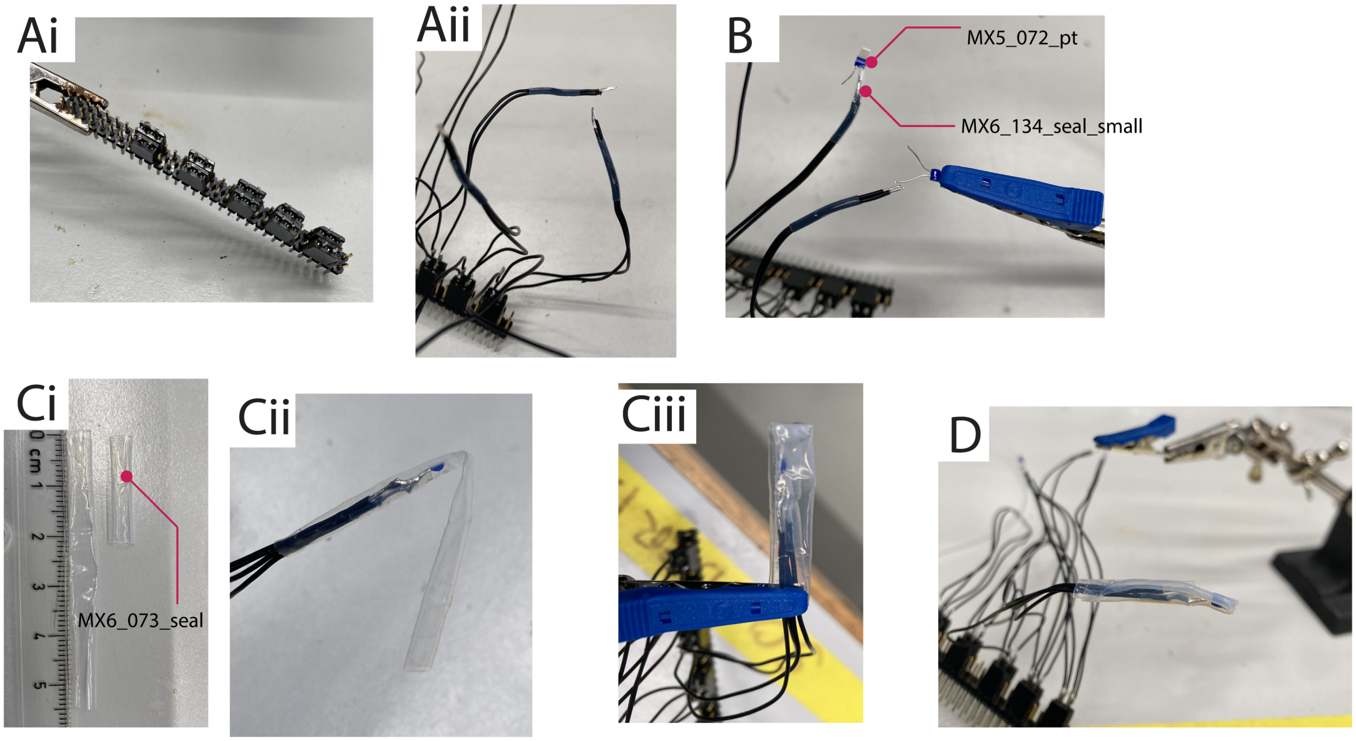


Figure S24: **A**i) Cut a standard 2.54 mm spaced 2 by 2 female header. ii) Attach 4 wires to the the pins (small diameter and heat resistant PTFE insulation recommended). **B** Connect 2 of the wires to one contact of the PT1000, insulate it with shrink tubing, and connect 2 wires to the other contact. **C**i) Cut 2 pieces of biocompatible shrink tubing. ii) Push one half of the long tube over the sensor and fold it to the side. iii) Push the short tube over the folded tube to hold it in place. **D** Use a heat gun at 400 °C to shrink the tube. Take care to not melt the solder tin.

##### A.5.3 Fan for heat sink and Peltier driver cooling

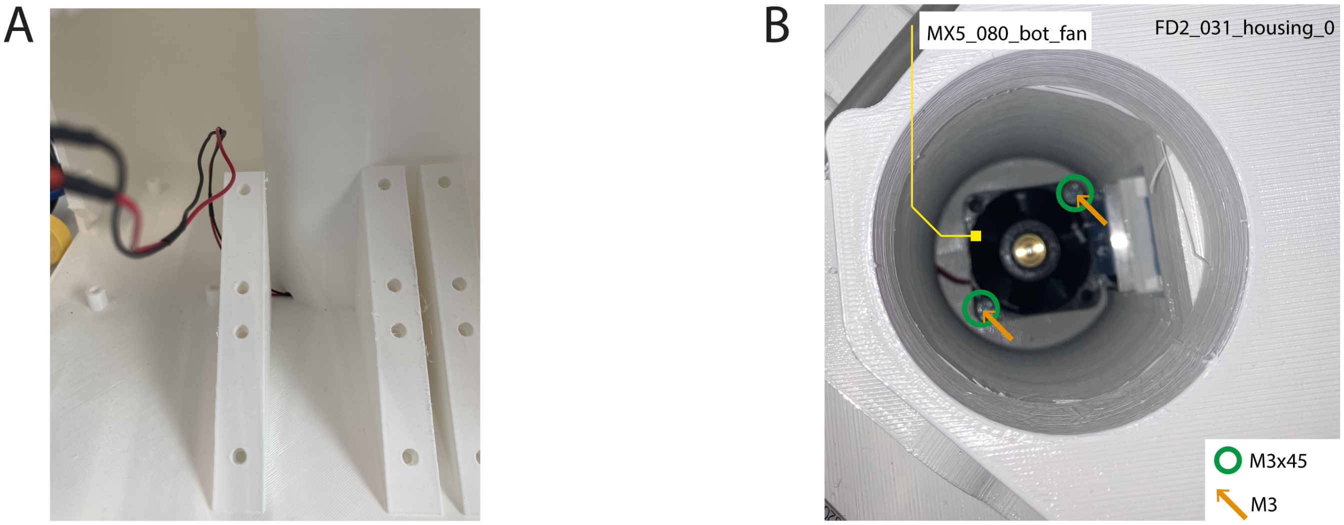


Figure S25: **A** Cut the fan wire and guide it out through the whole. Attach an extension to reach a 9 V connector on the power board. **B** Place the fan and insert the screws from the top. Counter the screws with a hex nut on the other side. Ideally use a magnetic screw driver to bring the screws in place.

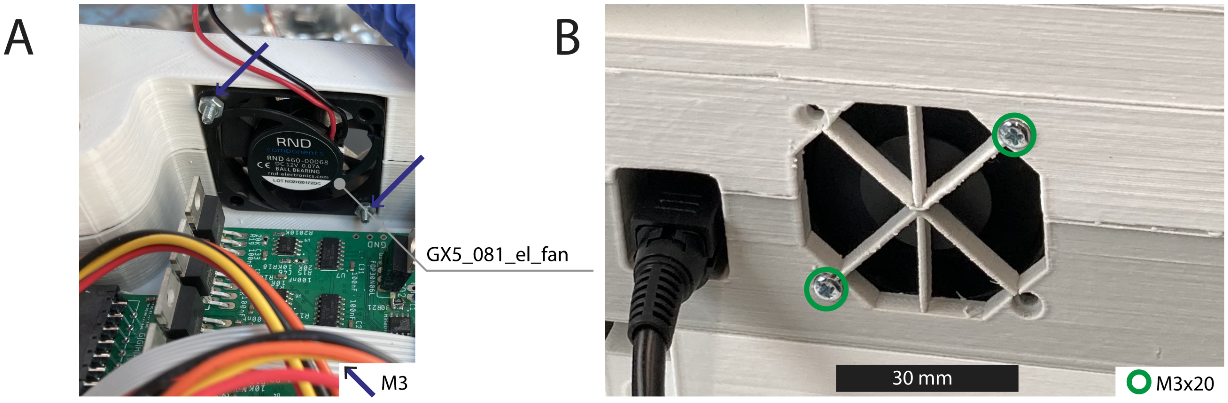


Figure S26: **A** Place the fan and attach a connector to the cables to plug it in to some 9 V power connector on the inkuflow or power board. **B** Insert the screws and use a hex nut as shown in **A** to counter it.

#### A.6 Electrophysiology

##### A.6.1 Electrophysiology board

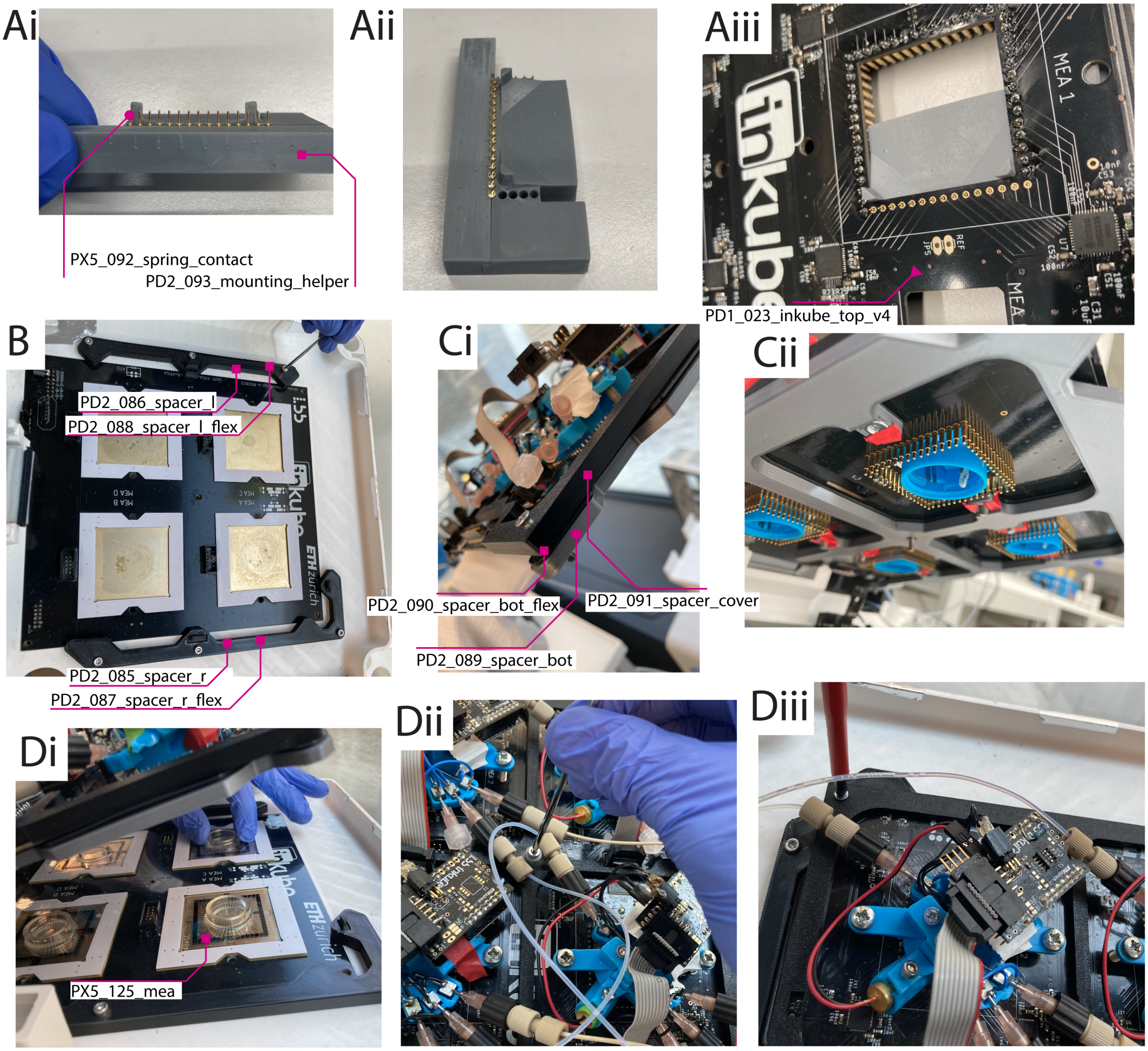


Figure S27: **A**i) Assemble the two parts of the mounting helper. ii) Use tweezers to place all pins such that the top rim sticks out. iii) Place the helper under the top board and solder the pins. Use low temperatures for soldering (depending on the solder tin 300 °C). **B** Mount the spacers on top of the bottom board. **C**i) Mount the spacers to the top board. ii) Insert the hex nuts to the top board spacer. Depending on printing tolerance hold them in place with some tape. **D**i) Slide the top board into the tilted slit on the side of the cable connections. Place the MEA. Take care of the reference electrode location. ii) Place the top board. Start tightening the center screw. iii) Continue with the corner screws.

#### A.7 Fluidics

##### A.7.1 Pump

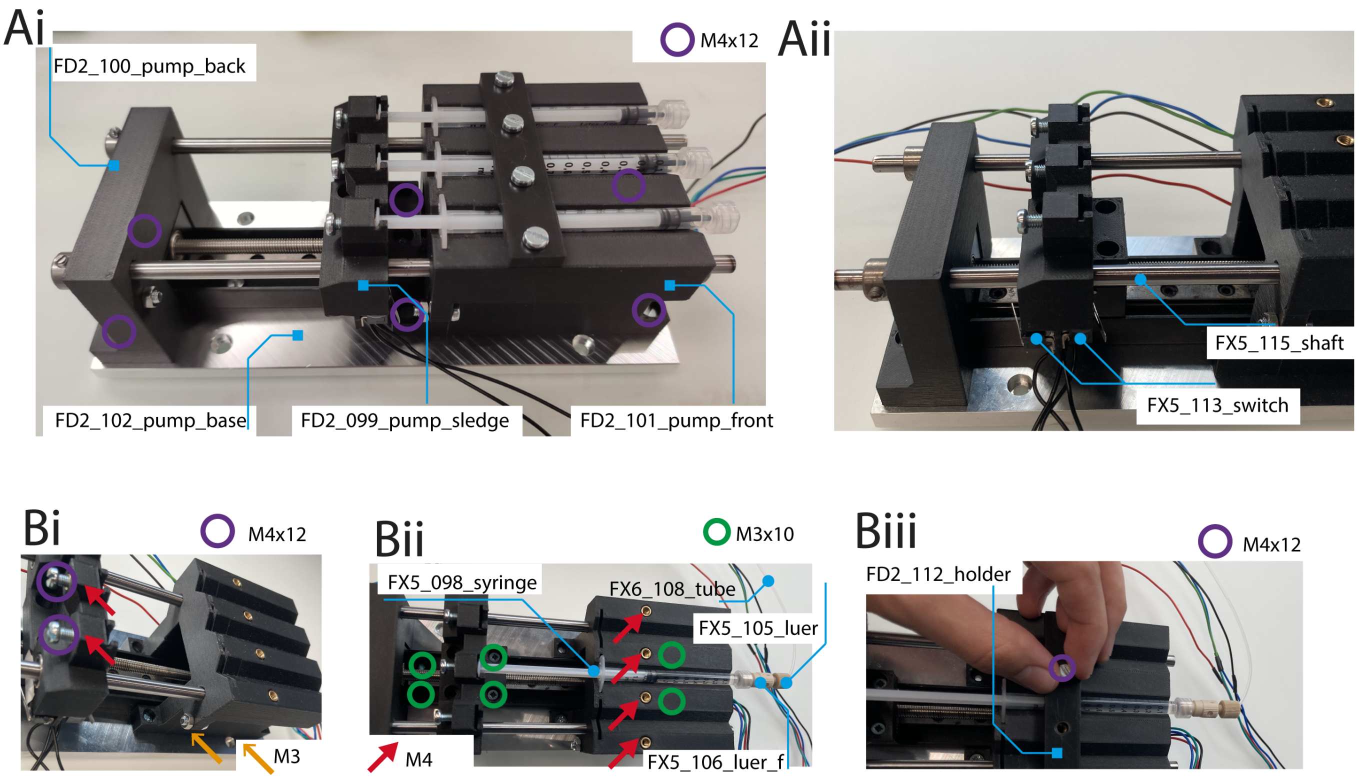


Figure S28: **A**i) Mount linear motor stage to base plate. ii) Mount the front part. Glue the safety switches to the sledge. Mount the sledge. Mount the back part. Insert shafts. **B**i) Mount screws to fix syringes. ii) Insert syringe. iii) Mount holder to hold syringes in place.

##### A.7.2 Multiplexing scheme

In Fig. S29 the multiplexing scheme of inkuflow is described. For each liquid a new path consisting of a syringe, a multiplexer, and a reservoir is required and for every MEA an extra port on the multiplexer is needed. Every multiplexer has one port for the syringe, which is connected through one valve to a reservoir, and an arbitrary number (in this case 4) valves with ports for the MEAs. The multiplexers always have a single valve open while the pump is being moved.

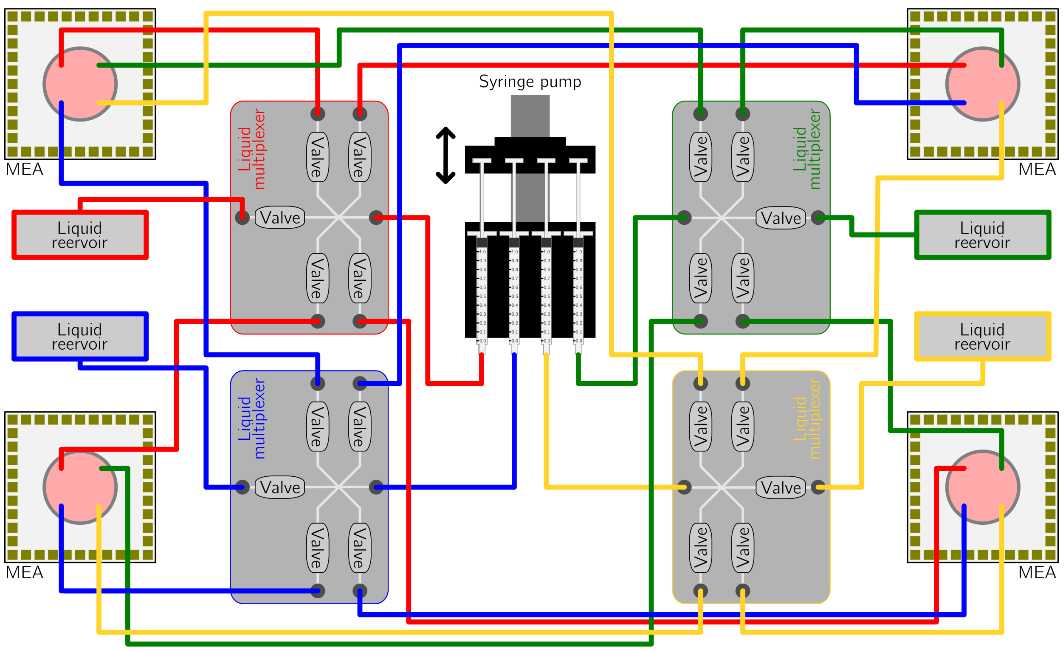


Figure S29: **Detailed block diagram of the inkube perfusion.** The perfusion system of inkube consists of 4 liquid reservoirs, 4 liquid multiplexers, and a single syringe pump. Pathways corresponding to the same liquid are shown in the same color.

##### A.7.3 Valve multiplexer

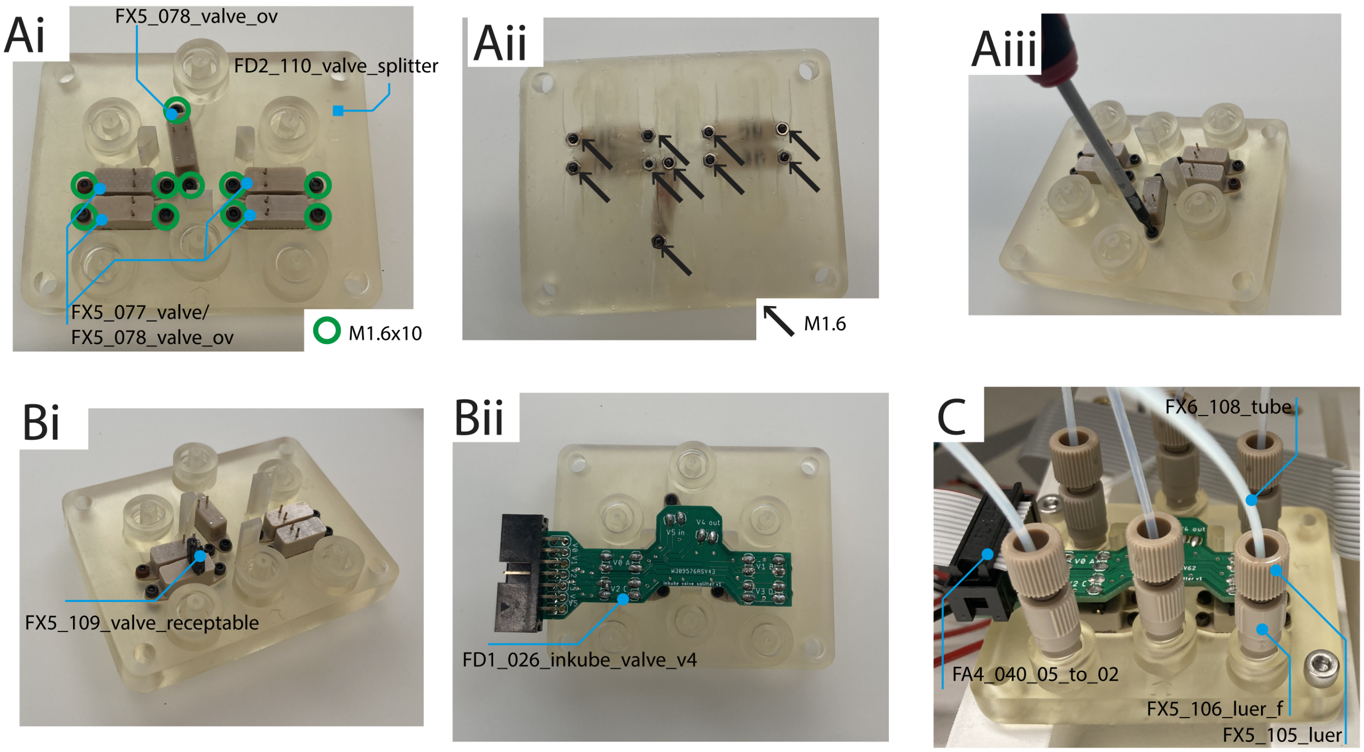


Figure S30: **A**i) Place valves on multiplexer. ii) and iii) Insert screws and counter with hex nuts. **B**i) Place preci pins (from valve PCB). ii) Mount PCB and solder connections. **C** Connect tubing through Luer locks.

##### A.7.4 Driver board and status LEDs

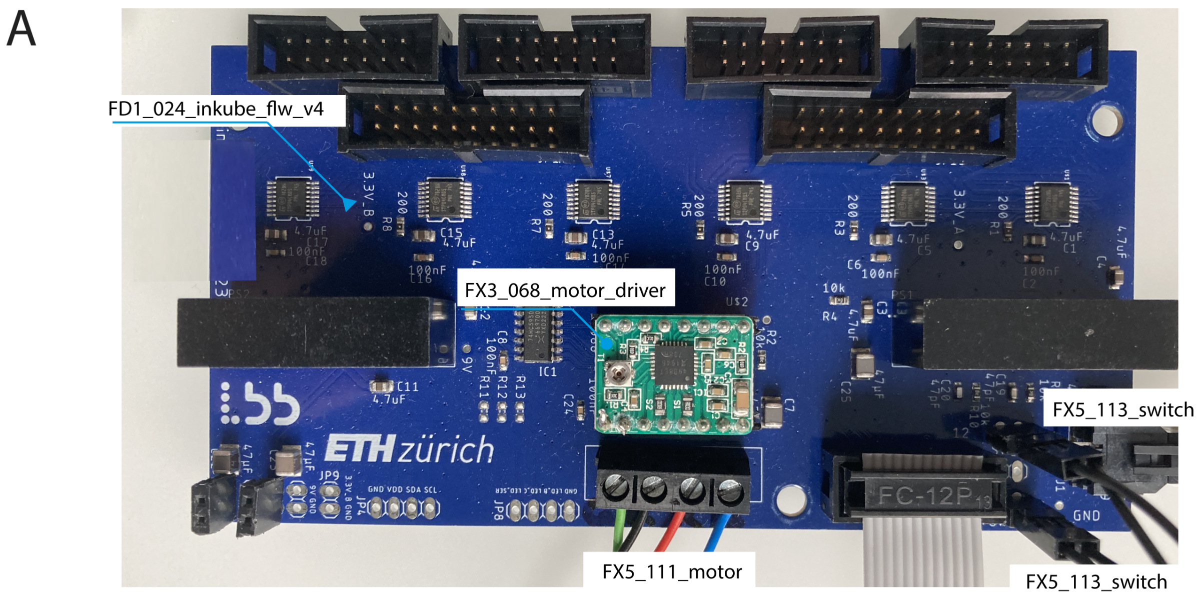


Figure S31: inkuflow board. Mount the motor driver PCB. Screw in the cables for the motor according to the labeling on the PCB. Connect safety switches for front and back according to labeling.

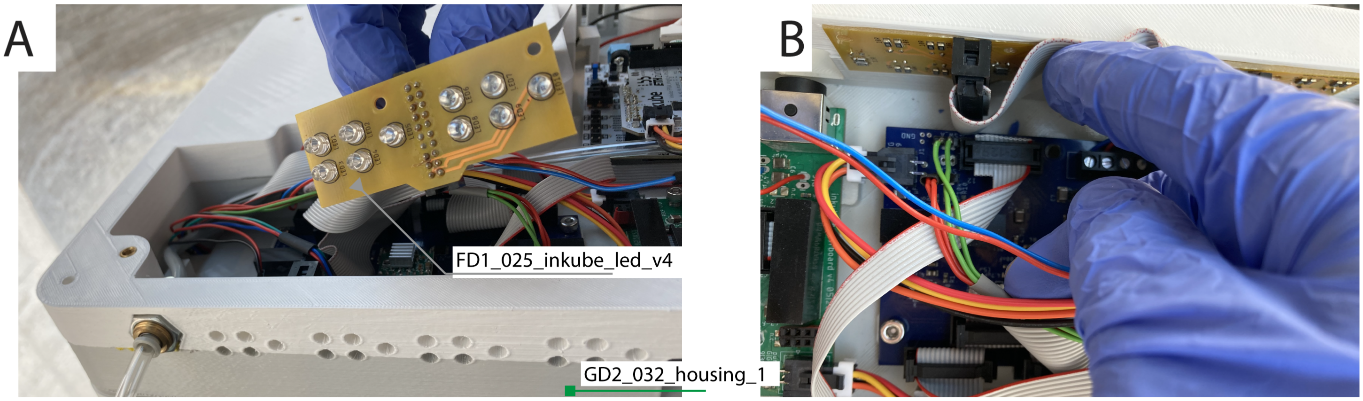


Figure S32: **A** Connect cable to back of assembled PCB. **B** Push PCB with LEDs into the wall of the electronics layer.

##### A.7.5 Liquid reservoir

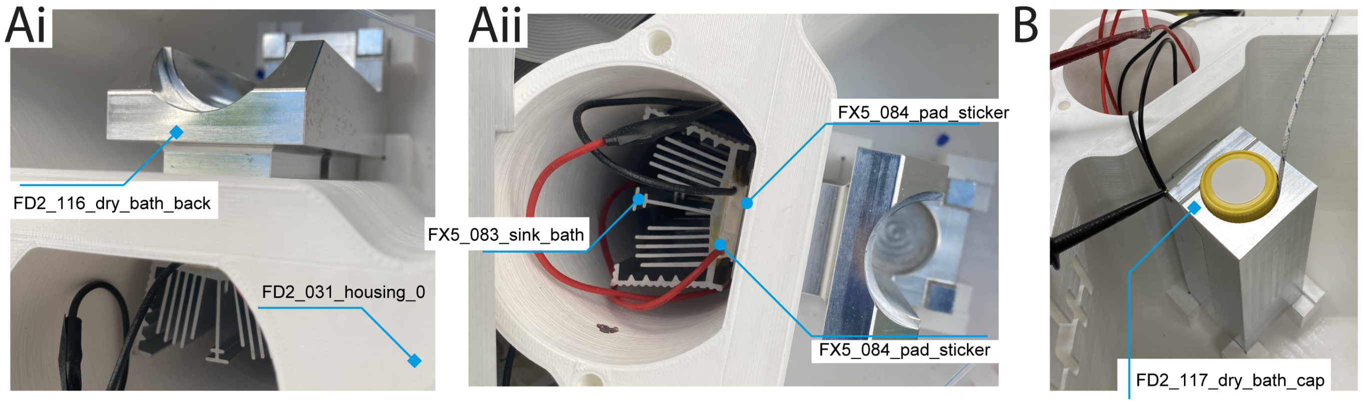


Figure S33: **A**i) Attach stickers to dry bath back. Insert back part through holes to ventilation channel in the fluidics layer. ii) Mount a second set of stickers on the Peltier elements. Mount the Peltier elements on the stickers of the dry bath back with the labeled side and the cables pointing upwards. **B** Mount the 50 mL falcon tube and place dry bath cap. Optionally add the insulation shield (FD2_130_dry_bath_cover) to prevent condensation.

##### A.7.6 Tubing and connection

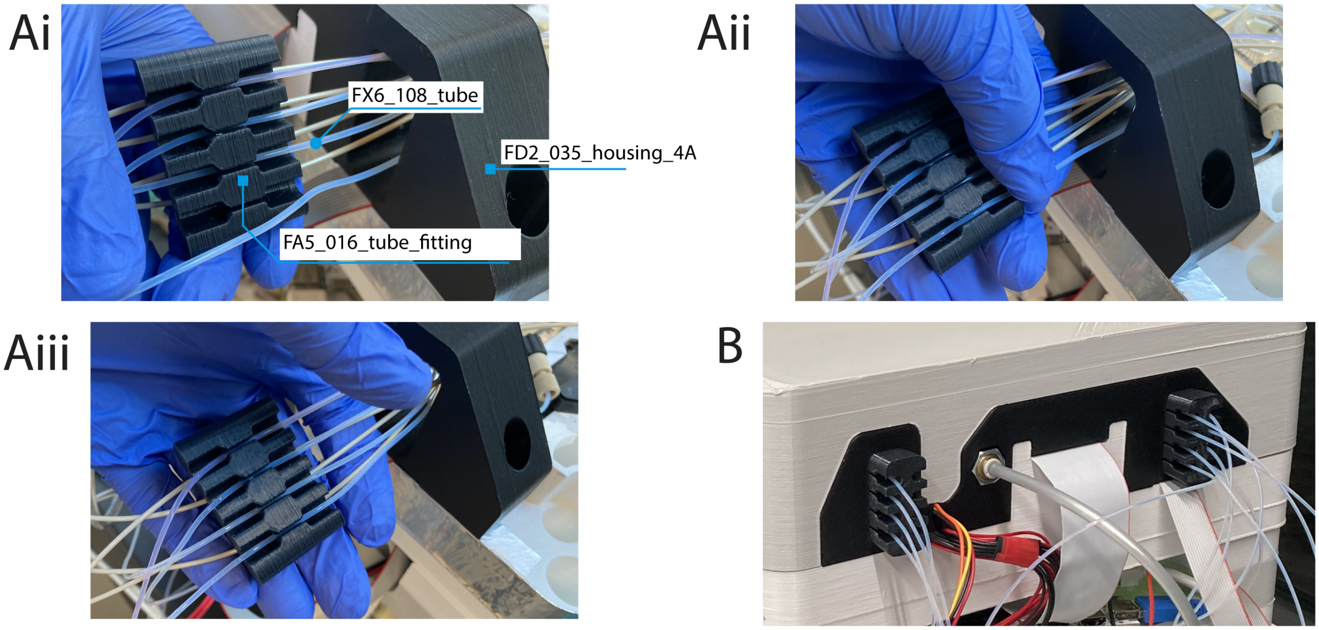


Figure S34: **A**i) Lead all tubes through the gaps in the incubation layer wall. ii) and iii) Bend and clip in the tubes. **B** Push the fittings in. Pay attention to not blocking the tubes, which can happen when pushing the tube fitting in too far.

#### A.8 Volume measurement

##### A.8.1 inkulevel

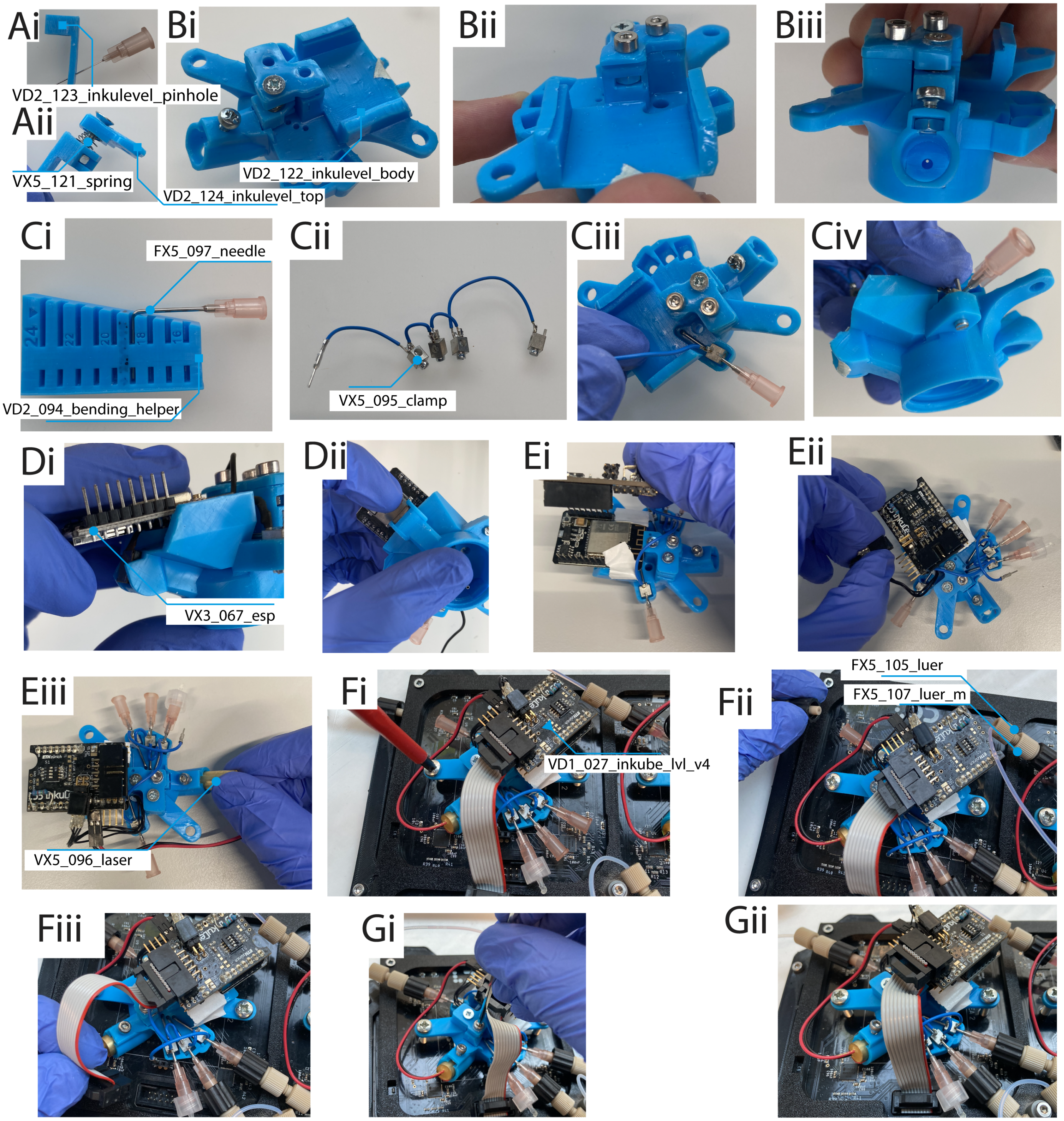


Figure S35: **A**i) Ensure the correct pinhole size by inserting a 0.45 mm syringe needle. ii) Mount the pinhole to the inkulevel top with a spring, a screw, and a M3 hex nut. **B**i) Insert this into the slit in the inkulevel body. ii) and iii) Insert hex nuts to screw the top part into place. **C**i) Bend the needles to the desired length. To insert the needle into the liquid 19 mm are recommend and used throughout this work. ii) Connect the clamps for grounding. iii) Insert the needle through the clamp. Place it on the inkulevel body. iv) Screw it into place. **D**i) Slide the ESP board in from the side. ii) attach the camera to the body. It should nicely click in. **E**i) Mount the shield. ii) Insert the temperature sensor and connect it according to the label on the shield. iii) Insert and connect the laser. **F**i) Screw inkulevel into place on top of the MEA. Insert the clamp cable to the reference connection on the top board. When removing the top board to exchange MEAs, these can be kept in place. ii) Connect the liquid tubes. iii) Make the bottom board connection. **G**i) Adjust the laser height until the reflection appears on the camera sensor. ii) Final setup.

#### A.9 Environment control

##### A.9.1 Inkusense board

Use the sensor board as previously published with [1]. Solder a pin header to the GND, $V_{SS}$, SDA, and SCL pin and connect as shown in Fig. S23C.

##### A.9.2 Reservoir resistive heater

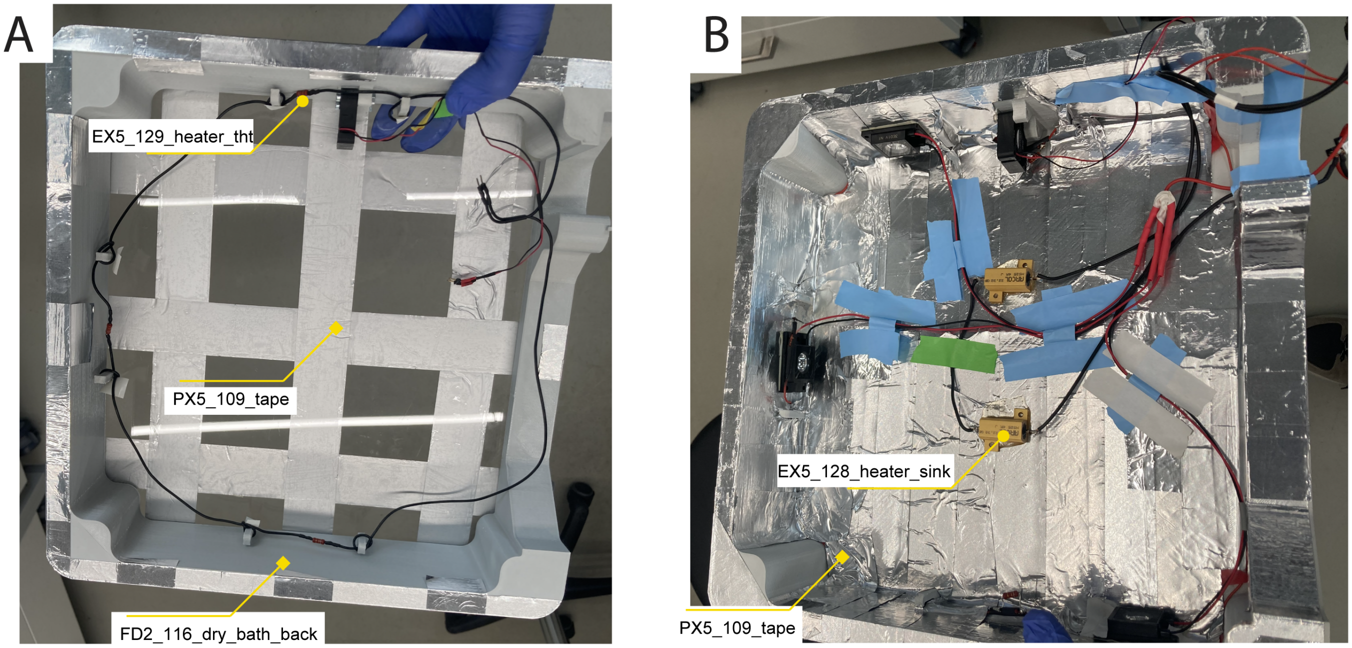


Figure S36: **A** Connect 3 of the 4 Ω resistors in series. Wrap the cable around the hooks to hold them in place. **B** Additionally, the resistors with heat sinks can be placed in the center and glued with epoxy.

##### A.9.3 $CO_2$ valve assembly

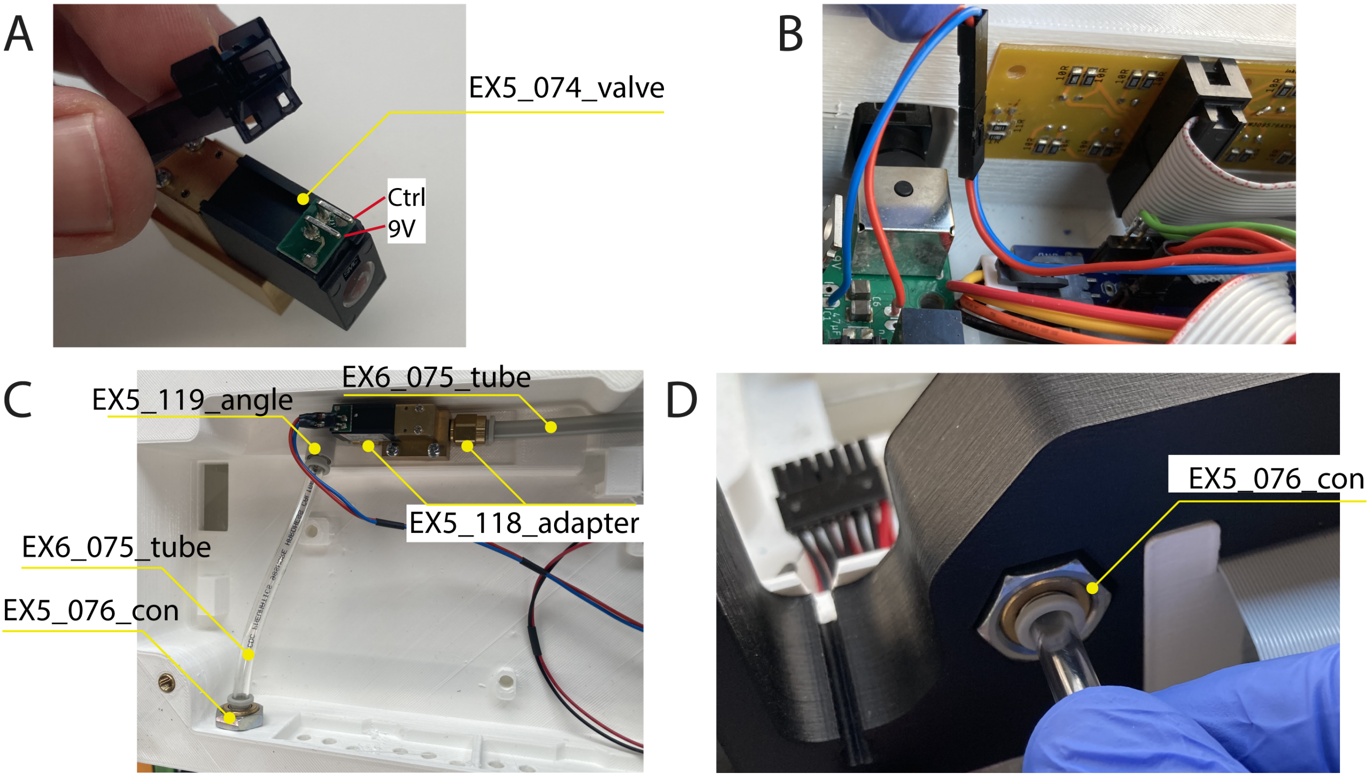


Figure S37: **A** Remove the clip on top of the connector pins of the valve. **B** Connect the valve to 9 V and the $CO_2$ control pin on the power board. **C** Screw in the tube adapters on both ends. Attach the angle to the input end. Mount the valve with the screws as shown in Fig. S9. Cut a piece of tube and connect the input with the wall connector in the electronics layer. Cut a second longer piece, connect it to the output, and lead it out the electronics layer. **D** Connect the output tube to the wall connector in the incubation layer.

##### A.9.4 Humidity heater

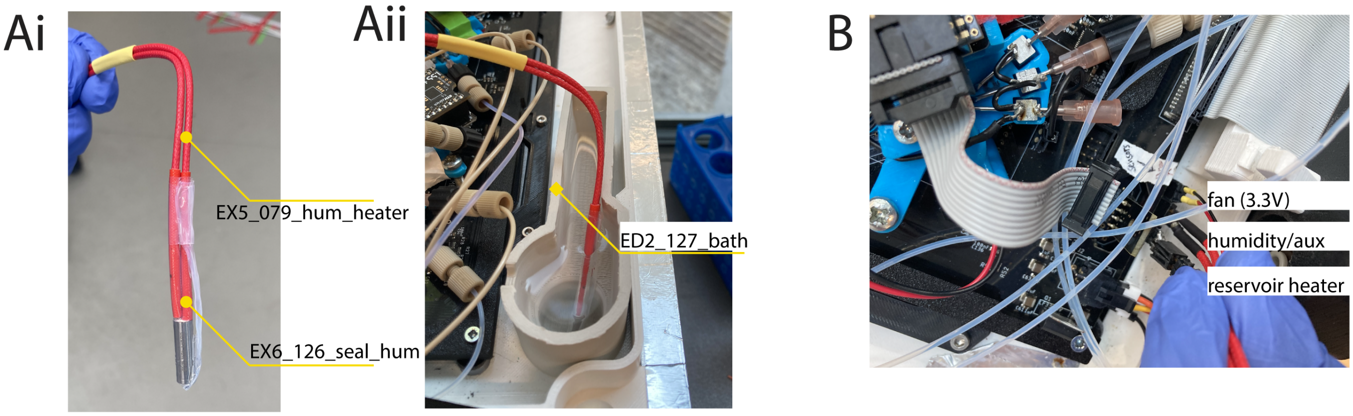


Figure S38: **A**i) Cut the cable of the heating cartridge to about 25 cm. Solder a pin header to the cable ends. Cut 2 pieces of PTFE shrink tube. Push the first piece half over the cartridge, bend it and hold it in place with the second piece. Use a heat gun at 400 °C to shrink the tube. ii) Fill the bath and insert the heater. **B** Connect the heater to the base board.

##### A.9.5 Fan for stable control

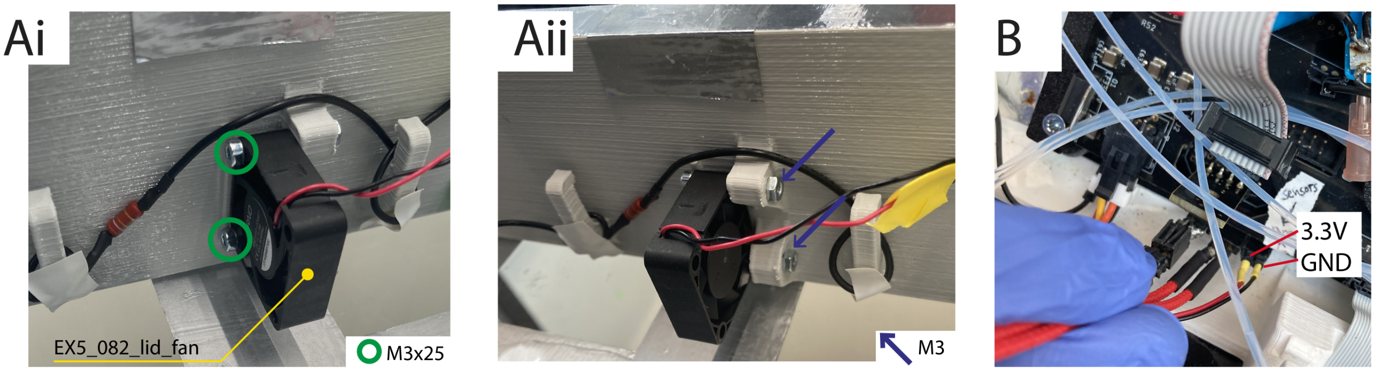


Figure S39: **A**i) and ii) Use 2 screws to mount the fan. **B** Connect the fan to the base board.

#### A.10 Equipment and unspecified components

- soldering iron and reflow oven
- LCD resin printer (Sonic Mini 4K, PHROZEN TECH CO., LTD., Hsinchu City, Taiwan), resins: PowerResinEu SG (Surgical Guide) (PowerResinsEU, Istanbul, Turkey), Phrozen Aqua Resin 4K PHR-RS1000AQG4K (PHROZEN TECH CO., LTD., Hsinchu City, Taiwan)
- extrusion printer for large parts (Ender 5 Plus, Shenzhen Creality 3D Technology Co Ltd, Shenzhen, China)
- extrusion printer for finer parts (Prusa i3 MK3S, Prusa Research a.s., Prague, Czech Republic)
- screwdrivers
- shrinking tube
- Crimping tool

- UART FTDI programmer or ESP32-CAM-MB Micro USB Programmer with CH340G Serial Chip (for flashing ESP32)
- screws as described layer figures
- M1.6, M2.5, M3 hex nuts
- M3, M4, M5 thread inserts
- cables and crimp contacts
- SD card for SoC

### B Software

An overview of the software architecture of inkube is provided in Fig. S40. It can be subdivided into 3 main parts: the software performing the low-level hardware communication on the SoC, the user interaction and electrophysiology analysis on the host PC, and the point detection on inkulevel. All parts and interfaces will be described in the following sections.

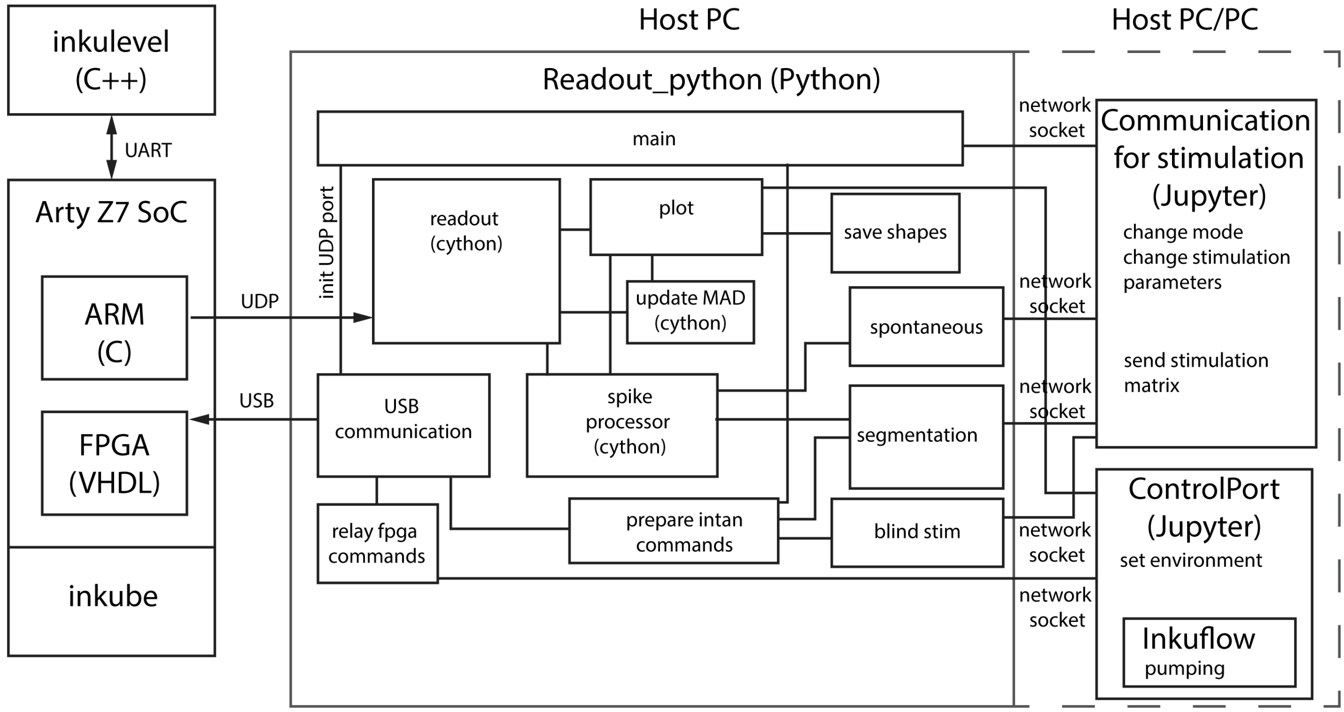


Figure S40: Overview block diagram of the software for the SoC, the host PC, and inkulevel.

#### B.1 SoC

inkube is built around the Arty Z7-20 (Digilent, Pullman, WA, USA), which contains a ZYNQ-7000 SoC (XC7Z020-1CLG400C, Xilinx, San Jose, CA). The SoC is programmed in VHDL and C. The block diagram of the SoC is given in Fig. S41.

The CPU of the ZYNQ SoC (see Fig. S41) is running at 650 MHz, while the DDR is running at 525 MHz. The Programmable Logic fabric clock runs at 100 MHz. The system uses two different types of Advanced eXtensible Interfaces

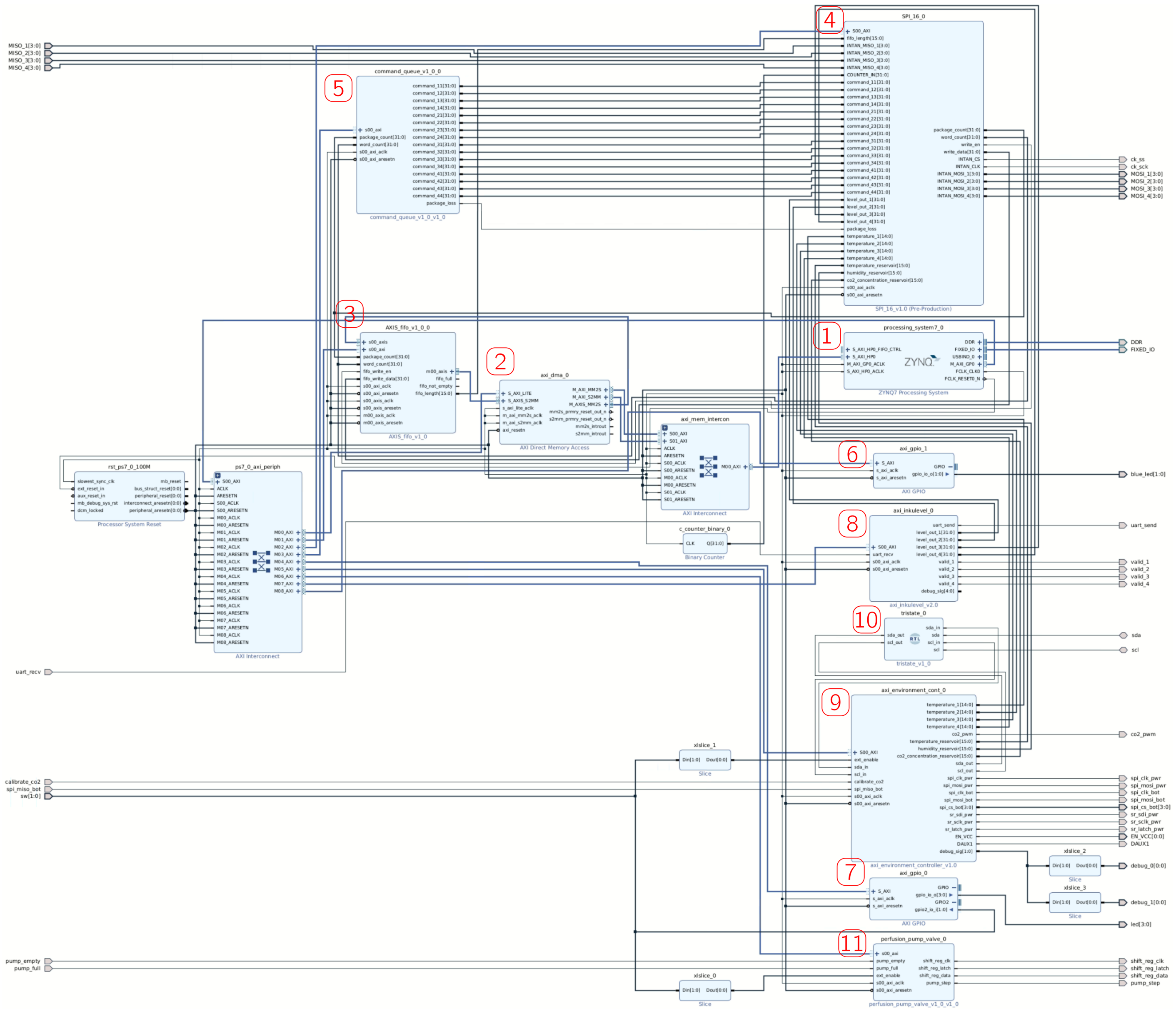

Figure S41: **Block diagram of the SoC.** **(1)** Hard core of the system. **(2)** Direct memory access to send data via the UDP protocol to the PC. **(3)** FIFO that is transforming the data from the INTAN chips into an AXI4-Stream. **(4)** Interface to control the INTAN ICs. **(5)** Interface to control the commands to be used for the INTAN ICs. **(6)** Interface to control the blue channels of the 2 Arty Z7 RGB LEDs. **(7)** Interface to interact with the switches and the mono-color LEDs of the Arty Z7 board. **(8)** Interface used to communicate with the 4 inkulevels. **(9)** Interface to measure and control the environment parameter. **(10)** Tristate Interface to implement allow for a high-impedance input for the $I^2C$ interface. **(11)** Interface for the perfusion system.

(AXIs) for inkube. Most of the interfaces use AXI4-Lite (AXI4L), except for the datastream sent from inkube to the PC, which uses an AXI4-Stream (AXI4S). The address ranges of different AXIs are given in Fig. S42.

| Name | Interface | Slave Segment | Master Base Address | Range | Master High Address |
|---|---|---|---|---|---|
| Network 0 | | | | | |
| /axi_dma_0 | | | | | |
| /axi_dma_0/Data_MM2S (32 address bits : 4G) | | | | | |
| /processing_system7_0/S_AXI_HP0 | S_AXI_HP0 | HP0_DDR_LOWOCM | 0x0000_0000 | 512M | 0x1FFF_FFFF |
| /axi_dma_0/Data_S2MM (32 address bits : 4G) | | | | | |
| /processing_system7_0/S_AXI_HP0 | S_AXI_HP0 | HP0_DDR_LOWOCM | 0x0000_0000 | 512M | 0x1FFF_FFFF |
| Network 1 | | | | | |
| /processing_system7_0 | | | | | |
| /processing_system7_0/Data (32 address bits : 0x40000000 [ 1G ]) | | | | | |
| /axi_dma_0/S_AXI_LITE | S_AXI_LITE | Reg | 0x4040_0000 | 64K | 0x4040_FFFF |
| /axi_environment_cont_0/S00_AXI | S00_AXI | S00_AXI_reg | 0x43C3_0000 | 64K | 0x43C3_FFFF |
| /axi_gpio_0/S_AXI | S_AXI | Reg | 0x4120_0000 | 64K | 0x4120_FFFF |
| /axi_gpio_1/S_AXI | S_AXI | Reg | 0x4121_0000 | 64K | 0x4121_FFFF |
| /axi_inkulevel_0/S00_AXI | S00_AXI | S00_AXI_reg | 0x43C5_0000 | 64K | 0x43C5_FFFF |
| /AXIS_fifo_v1_0_0/s00_axi | s00_axi | reg0 | 0x43C0_0000 | 64K | 0x43C0_FFFF |
| /command_queue_v1_0_0/s00_axi | s00_axi | reg0 | 0x43C2_0000 | 64K | 0x43C2_FFFF |
| /perfusion_pump_valve_0/s00_axi | s00_axi | reg0 | 0x43C4_0000 | 64K | 0x43C4_FFFF |
| /SPI_16_0/S00_AXI | S00_AXI | S00_AXI_reg | 0x43C1_0000 | 64K | 0x43C1_FFFF |

Figure S42: **Address ranges of different AXIs of inkube.** While large 64K address ranges are reserved for all AXI4Ls, not all of these addresses are actually used.

The AXI4S is controlled through a direct memory access (DMA, see Fig. S412), which packages data directly into UDP data. The UDP/IP headers and checksum are created by the processor of the ZYNQ SoC. For that, the IPv4 (192.160.10.10) and MAC (00:0a:35:01:02:03) address of inkube are hard-coded but can be changed in the C code. Similarly, for simplicity the PC connecting with inkube must have the IPv4 address of 192.168.10.1. In order to connect to inkube, an ARP request with these IP addresses must be sent to inkube. Inkube is capable of responding to ARP requests. However, ICMP is not implemented for inkube and cannot be used. Furthermore, to build up a connection with inkube, it also must be connected to the PC. Both the Ethernet and USB connection must be correctly set up for inkube to be capable of communicating with the PC to interact with inkube. The AXI4S is controlled by AXIS_fifo (see Fig. S413, which is containing a FIFO of width 32 bits and a depth of 32,768 entries. The status of the FIFO can be read out with an additional AXI4L (address range: 43C0XXXXh). The FIFO predominantly receives its data from the SPI_16 interface.

The SPI_16 interface (see Fig. S414 is communicating directly with the INTAN ICs through a Serial Peripheral Interface (SPI) with a shared chip select and clock signal, but 16 independent data channels; one for each IC. The SPI is running at 12.5 MHz. The whole interface is controlled by an AXI4L (address range: 43C1XXXXh). This interface also initializes the INTAN ICs at startup and sends data to the AXIS_fifo interface, which is then transformed into UDP packages through the DMA. Each UDP package also contains the parameters of the inkube environment such as $CO_2$ concentration, temperatures, humidity, and inkulevel values, which are also sent to the FIFO. Finally, this interface is also sending commands to the INTAN ICs. With each measurement cycle, where all 16 electrodes of an INTAN are measured and read out, 4 commands can be sent to each INTAN IC in addition. However, inkube currently only adds the INTAN responses of the first two commands to the FIFO. However, all commands are fed into the FIFO as a read back and to allow for frame perfect timing reconstructions. The commands are controlled by the command_queue interface. The SPI_16 interface is creating global counters “word_count” and “package_count”. The package_count counter is counting once for every measurement frame (32 bit, cycles approx. every 90 sec), while the word_count counter is used as a synchronization tool within a measurement frame.

The command_queue interface (see Fig. S415 is responsible for sending out the commands to the SPI_16 interface at the requested time points. The data is sent to the interface through a FIFO of a width of 2080 bit and a depth of 32. Each entry is a set of 64 commands (4 for each INTAN IC). These are the 4 commands used for a single measurement cycle. Since each command has 32 bit, this requires a total of 2048 bit. The remaining 32 bit decide on the time when the command shall be executed, which is based on the package_count counter. Since this counter cycles every 90 s, the commands need to be added at the appropriate time to hit the correct cycle. In addition, only the oldest element in the FIFO is considered at any time. As a consequence, commands need to be added in the order that they need to be executed. The FIFO itself does not sort the commands by time. The commands are sent to the command_queue interface through an AXI4L (address range: 43C2XXXXh) from the processor, which in turn received commands from the PC through the USB connection.

inkube is using the switches and LEDs that are connected to the ZYNQ on the ARTY-Z7. The blue channels of the two RGB LEDs of ARTY-Z7 are controlled by the axi_gpio_1 interface (see Fig. S416), controlled by an AXI4L (address range: 4121XXXXh). If the two blue channels are switched off, this means that inkube has received data through the USB connection. In addition, the axi_gpio_0 interface (see Fig. S417) is connected to the two switches and the 4 mono-color LEDs. The 4 LEDs encode (from 0 to 2) the state of the left switch, whether the command queue is full, and whether the command queue is not empty. Finally, the 3$^{rd}$ LED flashes every time an Ethernet package is received. The two switches are enabling environment control and are discussed further in later paragraphs.

The 4 inkulevels communicate with the ZYNQ SoC through the axi_inkulevel interface (see Fig. S418). It is controlled with an AXI4L (address range: 43C5XXXXh). Specifically, a daisy-chain has been implemented through a universal asynchronous receiver-transmitter (UART) protocol. The measured level values are sent to the SPI_16 interface, while the validity of the measurement of the 4 levels is visually described by the color of the red and green channels of the 2 RGB LEDs.

To control the MEA temperatures, as well as the temperature, $CO_2$ concentration, and humidity levels of the reservoir, inkube is using the axi_environment_cont interface (see Fig. S419). This interface is controlled with an AXI4L (address range: 43C3XXXXh) and controls an adjustable PI controller. The controller needs to be enabled by the Arty Z7 switch 1. This way, the environment control can be easily switched off while the rest of inkube continues running. In addition, the controllers can be switched on and off through the AXI4L. The axi_environment_cont interface is communicating with the temperature, humidity and $CO_2$ sensors as well as the environment actuators. For that, both an $I^2C$ interface and SPI has been implemented. An $I^2C$ interface is requiring a tristate input, which is provided by the tristate interface (see Fig. S4110).

The perfusion system is controlled through the perfusion_pump_valve interface (see Fig. S4111). An AXI4L (address range: 43C4XXXXh) controls this interface, too. The controller needs to be enabled by the Arty Z7 switch 0. Both the syringe pump and the valves of the liquid multiplexers are controlled through shift registers, except for the output signal, which controls the step count of the pump. The interface also receives end-of-range information from two switches integrated into the syringe pump.

#### B.2 Python

The Python software can be divided into the readout process and 2 classes for interacting with the process, for example within a jupyter notebook. The readout process is started by executing `main.py`. This will establish communication with the SoC via USB and UDP. Subsequently, the child processes are started. The inkube software can run in 4 different modes, which can be set through the `ControlPort` interface.

0 idle: standard mode of operation, no spike times are recorded and no stimulation is executed

1 spontaneous: mode for spontaneous recording, spike times are continuously recorded and sent out in milliseconds since package ID reset in chunks of up to 1.5 s to the `Communication` instance

2 closed-loop stimulation: mode for stimulating the network and record spikes for a pre-defined period after stimulation as latency after recording start, spike times are sent to the `Communication` instance and a new stimulus is received. Stimulation can be applied at up to 8 different timepoints before recording start.

3 blind stimulation: mode for only stimulation, no spike times are recorded. Stimulation pulses are sent in chunks, which circumvents the limit of 8 timepoints

`readout` The data stream from the SoC arrives at a frequency of 17,361 Hz. One data frame consists of the package ID, received commands counter, status information, sensor data, and the data of the electrophysiology chip (see Supplementary Data 02_Code/03_Additional_Code_Documentation/package_format.ods). It is received through the `socket` class in the process `readout`, which is written in Cython. The package ID and the counter for the received commands are copied into shared variables. The status and sensor data is copied from the frame into shared arrays with the plot process. The voltage traces are processed in a parallelized for loop. There, the filtering is applied and spike detection is performed. The filtered data, the data for thresholding, and the spike times are then copied into shared arrays with the plot process. Additionally, there is the option to extract the spike waveforms and send them to the plot process. The detected spikes are also put on a pipe to the spike processor.

`update MAD` The threshold is computed on the data shared with the plotting process. Furthermore, the noise levels are computed here. Update rate and sampling can be adapted in the process call. The threshold update can be enabled or disabled through a checkbox in the GUI.

`plot` The plot process runs multiple threads that update the data for plotting. In the same thread, temperature, level, and spike shape data can be saved in chunks. The main thread hosts the application with the GUI, based on the PyQt6 library. The GUI is organized into different tabs, which can be selected on the top left.

The first tab is the signals tabs, where the voltage traces of 60 electrodes are plotted. In the standard settings the last 500 ms are shown in oscilloscope mode at an updated rate of 4 Hz. When spikes are detected, a triangle is plotted at $y = 0$ on the voltage trace. An example plot is shown in Fig. S43.

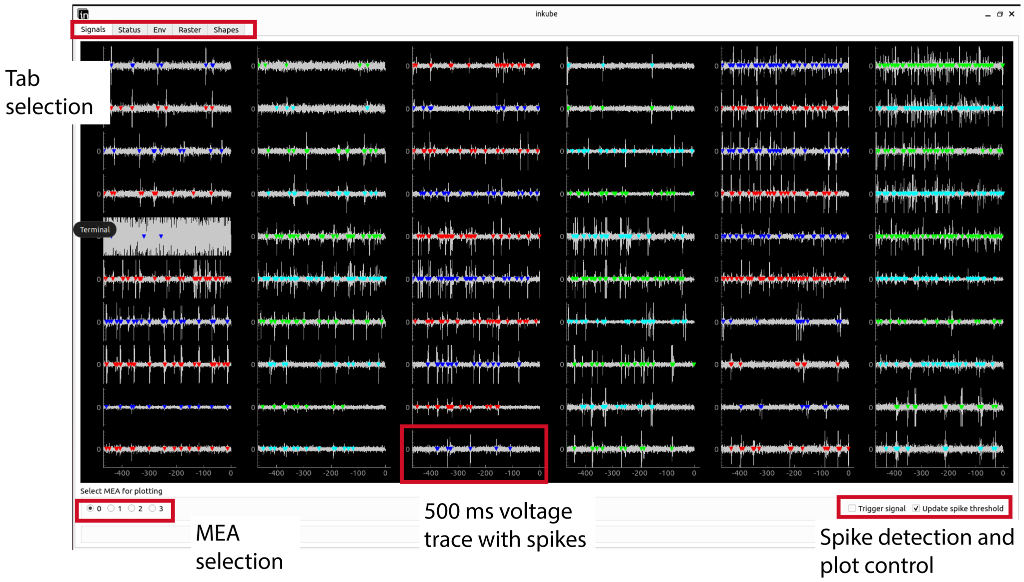


Figure S43: Signals tab of the inkube GUI showing the past 500 ms of the voltage signals of all 60 electrodes of an MEA. Detected spikes are marked with triangles. The y-axis is plotted in $\mu$V.

In the status tab depicted in Fig. S44, selected status data is shown over the past 500 ms. The first plot always shows the incrementing package ID. Additional information can be extracted from the aforementioned UDP package documentation.

The third tab shown in Fig. S45 provides information about the culture environment, the medium volume, and the noise levels of the electrophysiology data. In the update thread of this tab, the sensor data of the culture environment and medium volume can be saved in chunks of roughly 5 min. Additionally, the data can be shared utilizing a queue accessible through a network interface with a `ControlPort` instance to enable feedbacked control of all parameters.

The raster tab can be seen in Fig. S46. In case of stimulation, the post stimulus time histogram is shown for a single network. Spikes on the different electrodes are depicted in the respective color as a single dot at the measured latency in milliseconds after the recording start. The applied stimulation up to 10 ms before the recording start is plotted at a negative latency. The y-axis comprises 90 s of data and is then reset.

The spike shapes tab depicted in Fig. S47 shows cutout spike waveforms of roughly 2 ms of a full MEA. The shapes can be saved together with the respective package ID to enable spike sorting.

`spike processor` The spike processor process receives the spike time and electrode through a pipe. It can operate in 3 different modes. For mode 0 and 3, spikes are read from the pipe interface and discarded. For mode 1, all spikes are sent to the spontaneous readout process. For mode 2, spike times are compared to the recording start and discarded, if they lie outside the recording window (usually 20 ms after stimulation). Once the first spike past the window is detected or an external segmentation signal indicates that the window is over, all spike times are translated into latencies after window start and passed to the segmentation process.

`spontaneous` The spontaneous thread receives spike times from the `spike processor` and shares them through the network interface with the `Communication for stimulation`. The spontaneous thread utilizes the `ElectrodeMapping` class, which translates the channel IDs into network and MEA position. The spike times are shared as tuples for each electrode in a numpy array of dimension $network \times electrode$.

`segmentation` The segmentation process is used for the closed-loop stimulation (mode 2). It divides the recording time, base don the package ID, into periods of usually 250 ms. At the start of a period, spike times are recorded during the

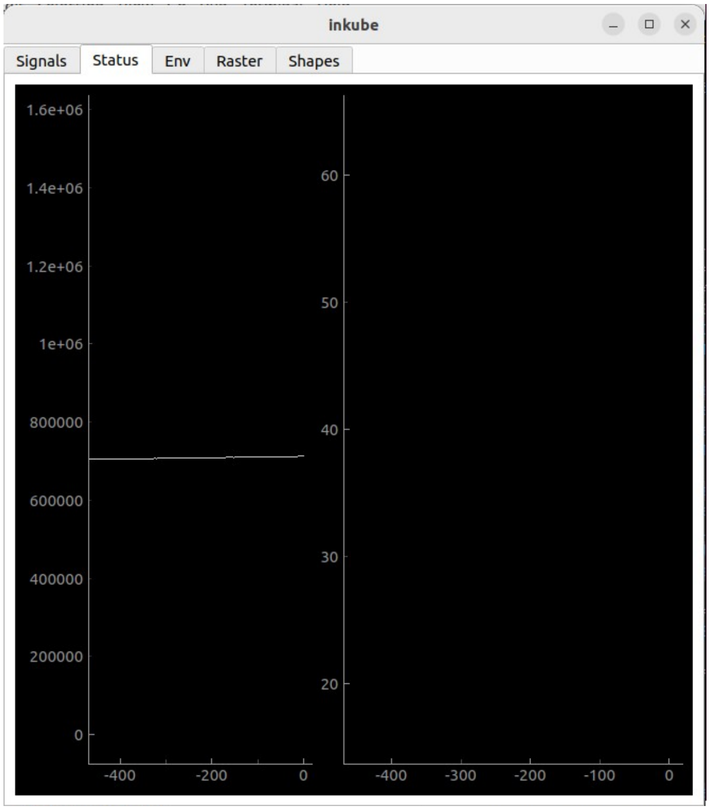


Figure S44: Status tab of the inkube GUI. Different status words can be plotted here over the past 500 ms. The left plot shows the linearly increasing package ID.

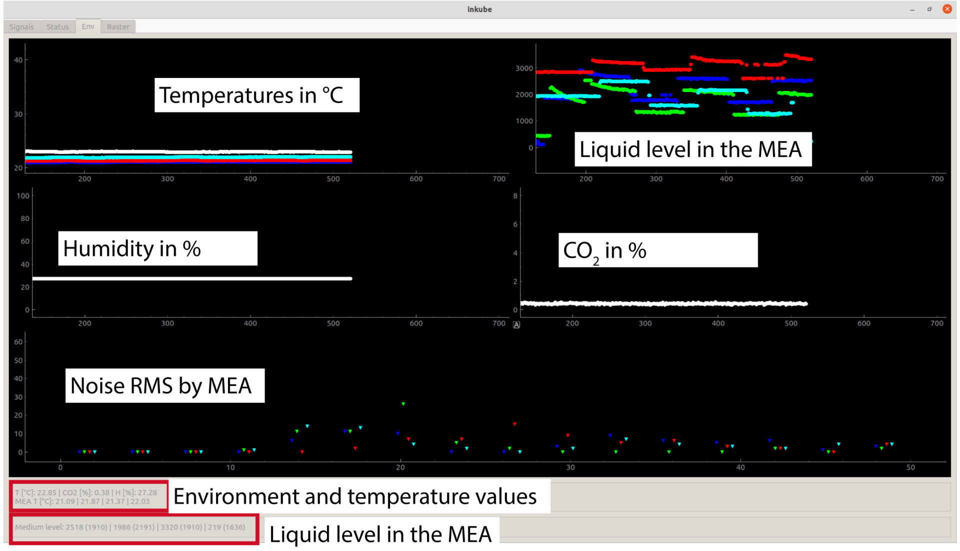


Figure S45: Env tab of the inkube GUI. The top 4 plots show data of the culture environment for up to 1 h. The x-axis unit is time in minutes. The top left plot shows the temperature of the reservoir in white and the 4 MEAs in the MEA colors (blue, green, red, cyan). The top right plot shows the inkulevel values for the 4 MEAs. The bottom left plot shows the relative humidity in the reservoir in % V/V and the bottom right plot show the $CO_2$ concentration in % V/V. All current values are also shown in the text fields in the bottom. The bottom plot shows the RMS noise for each MEA in the respective color with the bins in µV on the x-axis and the electrode count on the y-axis.

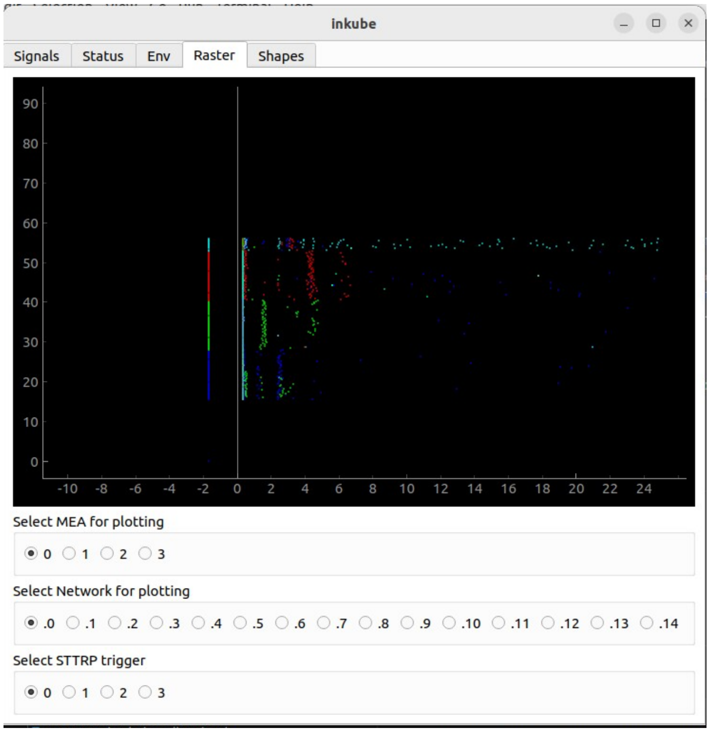


Figure S46: Raster tab of the inkube GUI. During stimulation (mode 2), the post-stimulus raster plots are shown, with the applied pattern up until 10 ms before recording start

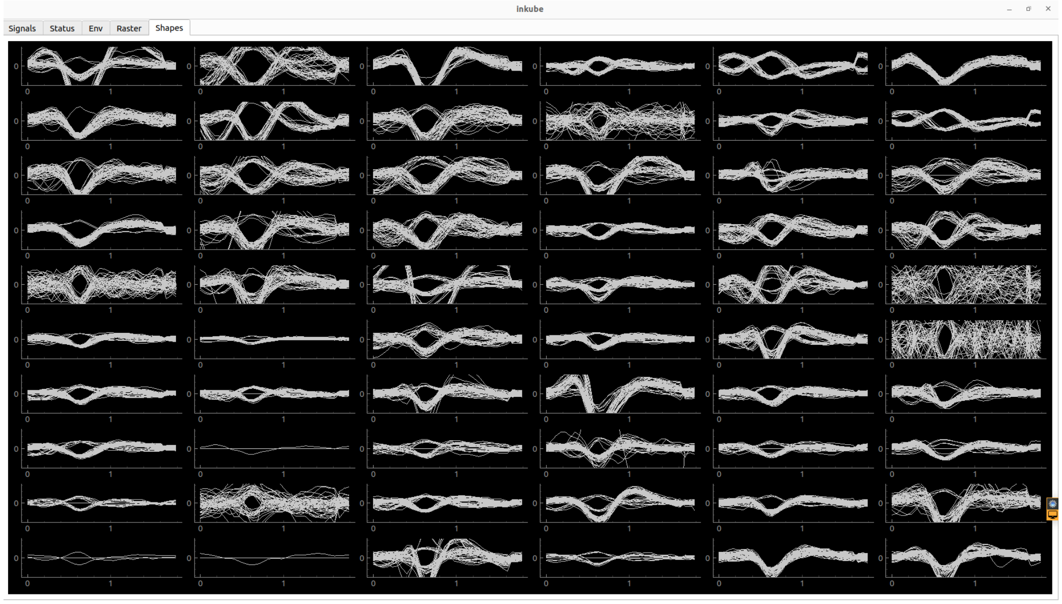


Figure S47: Shapes tab of the inkube GUI. When a spike is detected, about 2 ms are stored of the voltage signal and plotted on top of each other.

recording window (usually 5 to 25 ms). Afterwards, the recorded spikes are organized by network and electrode, similar to the spontaneous thread, and translated into latencies with respect to the window start. The spike matrix is subsequently sent out to the `Communication for stimulation` instance together with information about the period: a flag whether a stimulus has been received and executed for this period, the executed stimulation matrix, and the ID of the period). The process then waits for a stimulation command from the `Communication for stimulation` instance, up until either a stimulation command is received with the period ID corresponding to the following period or the maximum wait time is over. If a valid stimulation command has been received, the stimulation is sent to the `prepare intan commands` process. The stimulation time points are received as relative values in delay in samples before recording start and translated into the corresponding package IDs for the commands. Once the stimulation is forwarded, the period ID is incremented and the corresponding updated recording window times are passed to the `spike processor`.

`blind stimulation` The blind stimulation process is an adapted version of the segmentation process. Here, no spikes are recorded and therefore no synchronization with the spike processor has to be achieved. The information sent to the `Communication for stimulation` instance is reduced to the period ID, stimulation flag, and stimulation matrix. Stimulation commands are however not executed in the current period before recording start but in the following period. In case the stimulation has to be performed at more than 8 different timepoints, the stimulation matrix is chunked and sent dynamically to the `prepare intan commands` process.

`prepare intan commands` This process receives the electrode IDs and timepoints as package IDs where a stimulation pulse should be sent. Additionally, a flag is received whether this is a first or last pulse of a stimulation sequence. For the first and last pulse, commands are sent to all chips for fast settle and high-pass filter pole shift in order to reduce the stimulation artifact. From the electrode IDs the chip ID is derived. The stimulation pulses are sent to the respective electrodes. An active discharge is performed after the pulse on all electrodes of a whole chip of which at least one electrode has been stimulated. The commands are then passed to the `USB communication` process.

`relay fpga commands` This process receives commands for the pump, valve multiplexers, and environment set values from a `ControlPort` instance and sends them to the `USB communication` process.

`USB communication` This process receives commands in the final format together with an ID indicating the command type and sends them to the SoC via USB, optionally with a handshake. The handshake is performed by including the package ID into the first byte of the command after the package preamble and synchronizing it withg the received packages counter in the UDP package sent by the SoC. Commands can be of following types:

1 send port for UDP communication, no handshake

2 send intan commands, with receive package id (handshake), directly forwarded to ASIC by the FPGA

3 send fpga commands, with receive package id (handshake), these are register writes directly on the SoC for e.g. pumping

4 send reset command that sets receive package counter to 0, with receive package id (handshake)

`ControlPort` Set values for the environment controller can be set through an instance of the ControlPort class. Furthermore, the sensor data can be received for closed-loop control. Pumping and switching of the valves can be performed through the `Inkuflow` class, of which an instance is a member of the `ControlPort` class. A logging file is created by the instance with the sent commands and timestamps.

`Communication for stimulation` The `Communication for stimulation` instance receives spikes either from spontaneous recording (mode 1) or from the recording window during closed-loop stimulation (mode 2). Stimulation pulses are sent back as a matrix of the format $n \times 3$, where n corresponds to the number of pulses. The columns correspond to the delay of the stimulation pulse in samples before the recording window start, the network ID, and the electrode ID. The ID of the period for execution is derived by incrementing the last received period.

#### B.3 inkulevel

inkulevel is built around an ESP32 microcontroller, which can be programmed in C++ using the arduino IDE. The code is based on the arduino ESP webserver example: `https://github.com/espressif/arduino-esp32/tree/09a6770320b75c219053aa1` `libraries/ESP32/examples/Camera/CameraWebServer` [2] The main functionality is the detection of the laser reflection on the sensor area. UART communication is used to change the settings of the camera, the point detection, and send out the data. The details for extraction and communication can be extracted from the code documentation.

### C Installation and operation

This section contains information on how to flash the embedded devices, enable USB and network communication of the SoC and the PC, and install and run the software.

#### C.1 SoC

The compiled SoC code can be found in the inkube git under `ArtyZ7_SoC/BOOT.bin`. Move the file to a micro SD card and insert the card into the Arty Z7 board. Set the jumpers: JP4 to SD and JP5 to REG.

#### C.2 PC connection

In order to enable the readout from the SoC, the USB and network communication with the PC have to be established. For this, power the SoC and follow the instructions in this section. Initially, the 2 RGB LEDs of the SoC should light up in blue. They indicate the status of the ethernet and USB connection.

##### C.2.1 Ethernet

Connect the ethernet cable to the PC. Adapt the settings of the port as shown in Fig. S48. It might be necessary to reset the SoC afterwards. Once communication has been established, one blue LED turns off. Proceed with the next section.

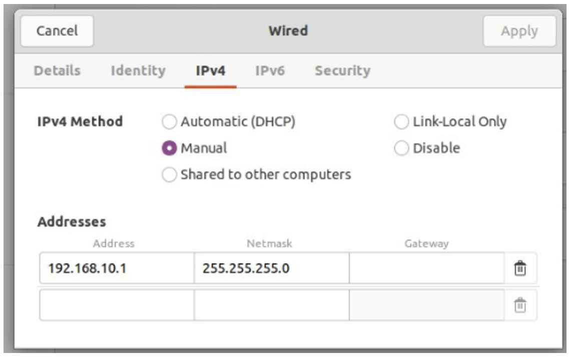


Figure S48: Ethernet settings to connect to FPGA.

##### C.2.2 USB

Connect the USB cable to a USB port of the PC. Follow the instructions below to allow communication of the PC with the SoC. The instructions can also be found in `USB_communication.py`.

- Open a terminal and execute the command: `lsusb`
- The result should contain a line with: `Bus 00x Device 00y:  ID 33ff:1234 Inkube This is the downlink USB connection of Inkube` with `00x` being the bus ID and `00y` the device ID.
- Show the permissions by executing the command: `ls -l /dev/bus/usb/00x/00y`
- if no access rights add the rules file `01-inkube_usb_permissions.rules` with content: `SUBSYSTEM=="usb",ATTR\{idVendor\}=="33ff",ATTR\{idProduct\}=="1234",MODE="0666",GROUP="plugdev"` to `/etc/udev/rules.d` by executing the command: `sudonano/etc/udev/rules.d/01-inkube_usb_permissions.rules`
- you might have to reboot the PC for the changes to take effect
- once the communication has been established the second blue LED should turn off
- as a sanity check you can execute the `USB_communication.py` file, this should make the LEDs on the FPGA blink (LD0 – LD3)

#### C.3 System startup

In order to use the GUI and control class interface, the Python readout process has to be running. For this follow the steps below.

- clone the inkube git repository
- install pip requirements
- build cython code for spike readout by executing in the directory `Readout_python`: `pythonsetup_filter.pybuild_ext--inplace`
- Execute `pythonReadout_python/main.py` to start readout and open the GUI. You can now use the example jupyter notebooks to change the environmental parameters, use the closed loop stimulation, and use the fluidic system.

#### C.4 Inkulevel

To receive feedback about the volume inside the MEA, inkulevel has to be flashed. For this one of the options listed in can be used together with the arduino IDE . With the FTDI chip, the GPIO 0 has to be pulled low. If the device is correctly flashed, the red LED should be constantly on once the board is powered. In case of a blinking red LED, an error occurred upon booting.

It is recommended to start by adjusting the laser while the image is streamed via bluetooth with the `bt_all_script_integral.py` script, which can be found in the inkube git under `inkulevel/Python_bt`. Be aware that this requires additional libraries. Then the settings can be tweaked for optimal detection. Standard parameters and values can be found in the example jupyter notebook `Experiments/02_Fluidics_Environment_Control.ipynb` in the inkube git.

### D System characterization

This section contains additional information that shows the performance of inkube. The limits of the actuators are tested. Additionally, the controller performance is tested by demonstrating the response to a change of the set value. Subsequently, the electrophysiology is covered with showing the spike detection and stimulation.

#### D.1 Actuator and controller performance

##### D.1.1 Reservoir temperature

The reservoir heating is tested with 8 Ω parallel to 12 Ω resistive heaters. Additionally, the water bath is heated with a 14.4 Ω heating cartridge to hold the relative humidity between 60 and 70%.

##### D.1.2 Humidity

##### D.1.3 $CO_2$ calibration and transient behavior

The $CO_2$ valve is supplied with 100% $CO_2$ at approximately 1.5 bar. The $CO_2$ sensor has an initialization routine, which is automatically performed at system start-up and a calibration routine, where the current $CO_2$ measurement is set to 0%. Due to the temperature dependence of the sensor, it is recommended to call the calibration routine once the reservoir has been successfully heated up to 37 °C.

##### D.1.4 MEA heating and cooling

A class B (according to EN 60 751) PT1000 is used for measuring the liquid temperature, as temperature stability is prioritized over temperature accuracy. Class B implies an absolute accuracy of the impedance of ±0.12%, which limits the absolute accuracy of the temperature to roughly 360 mK at 37 °C. Drifts due to self heating are not expected to have a noticeable effect with respect to the quantization error.

#### D.2 Fluidics

The stepper motor based syringe pump is characterized in the following section. Measuring the pumped volume yielded 108.025 µL for 10,000 steps ($\sigma$=0.255 µL, n=5). The linearity of the pump across the range of the stepper motor is tested by performing pumping of 100, 500, 1,000, 5,000, and 10,000 steps on the motor, starting at 20,000 and 60,000 steps. The

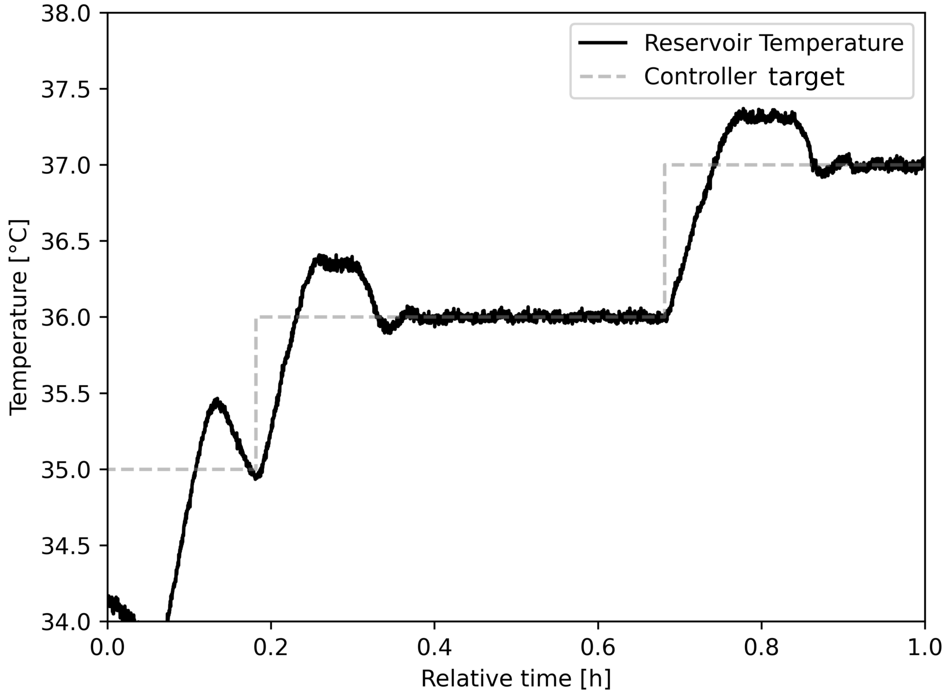


Figure S49: Reservoir temperature when changing the set values (dashed line).

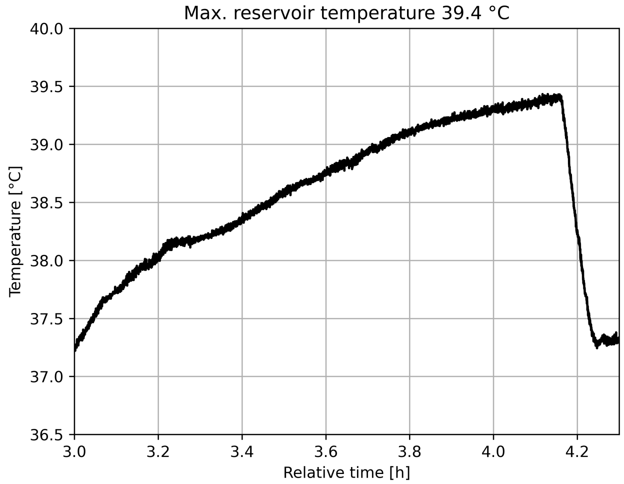


Figure S50: Maximum reservoir temperature achieved with the MEA temperatures at 37 °C and the reservoir set to maximum heating.

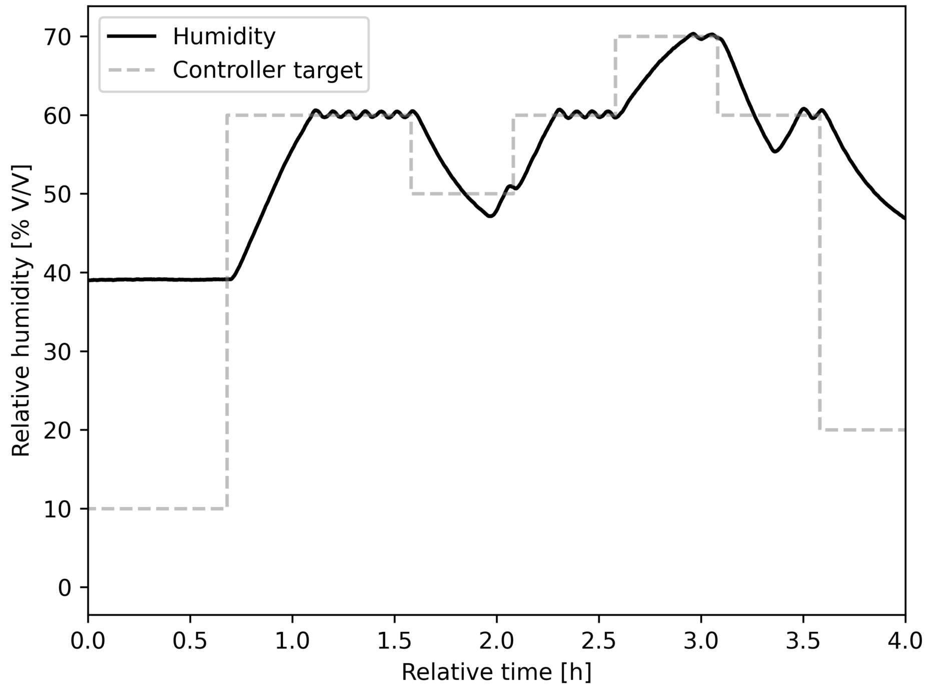


Figure S51: Reservoir relative humidity when changing the set values (dashed line).

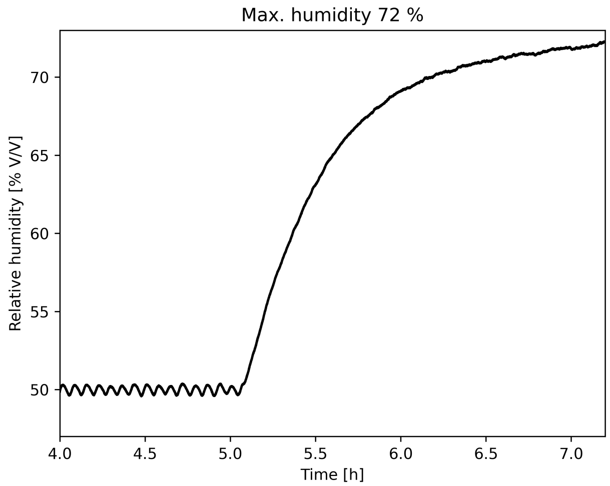


Figure S52: Maximum reservoir relative humidity achieved with the MEA and reservoir temperature at 37 °C and the bath heater set to maximum heating.

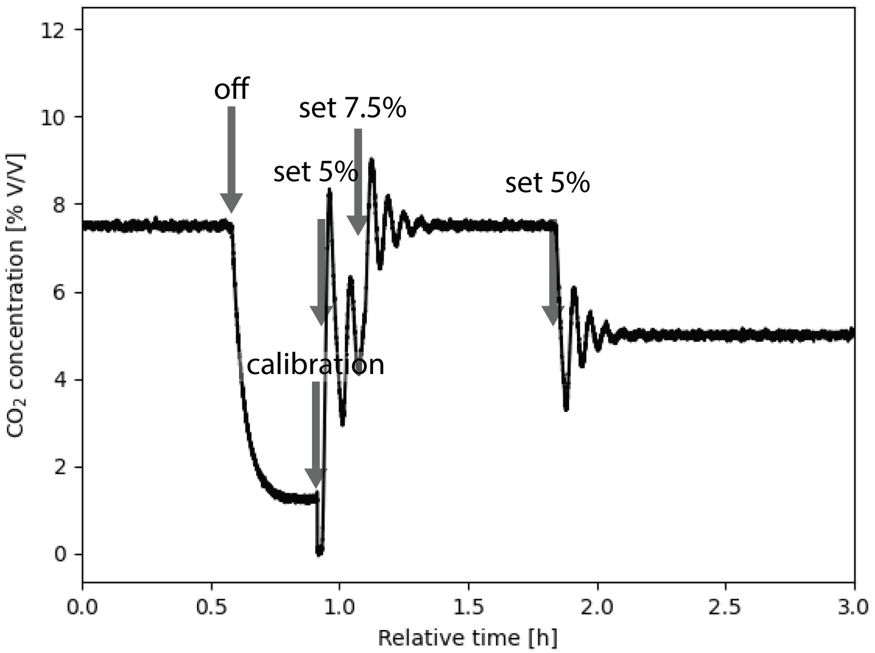


Figure S53: $CO_2$ value upon turning the valve off, calling the calibration routine, turning it on again at a set value of 5%, increasing the set value to 7.5%, and reducing it to 5% again. Time points are indicated with grey arrows.

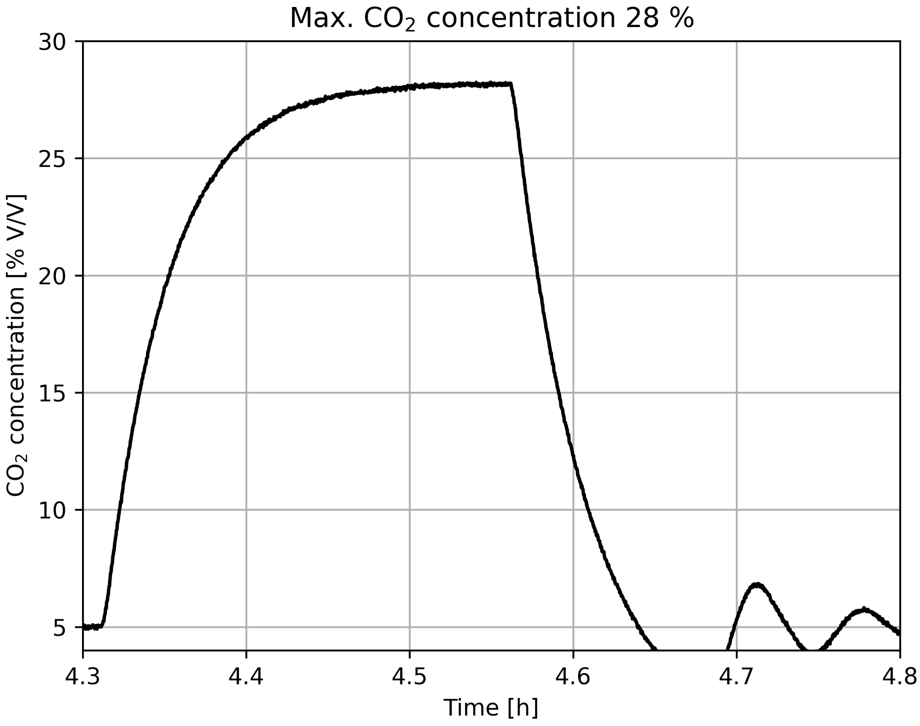


Figure S54: Maximum $CO_2$ value with the reservoir at 37 °C. The pulse width modulation of opening the valve is limited in order to increase the resolution at smaller values.

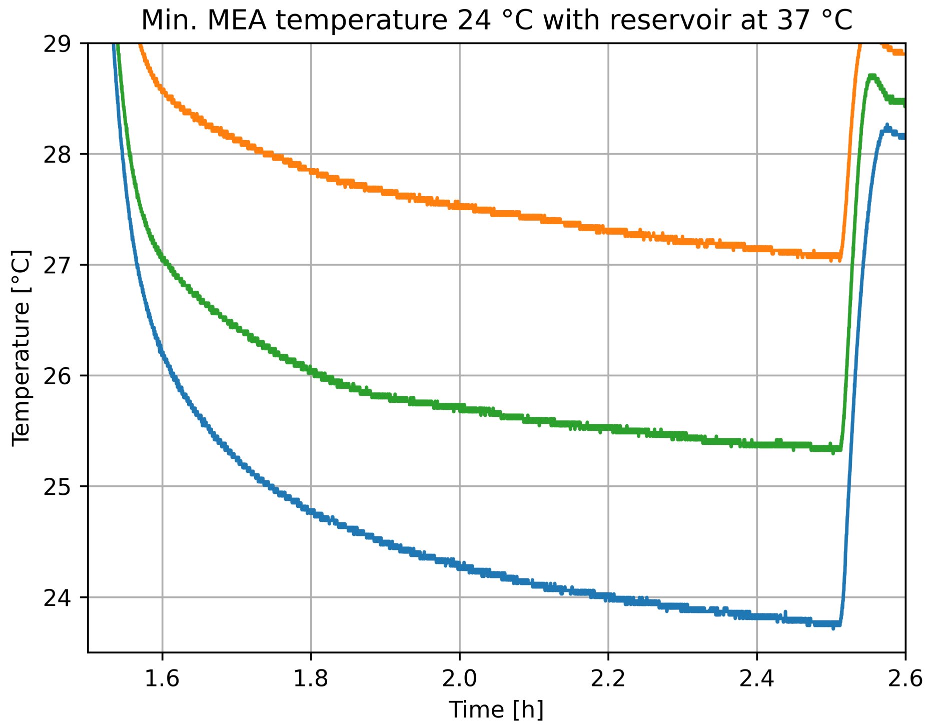


Figure S55: Minimum medium temperature achieved with the reservoir temperature at 37 °C and the thermoelectric devices set to maximum cooling.

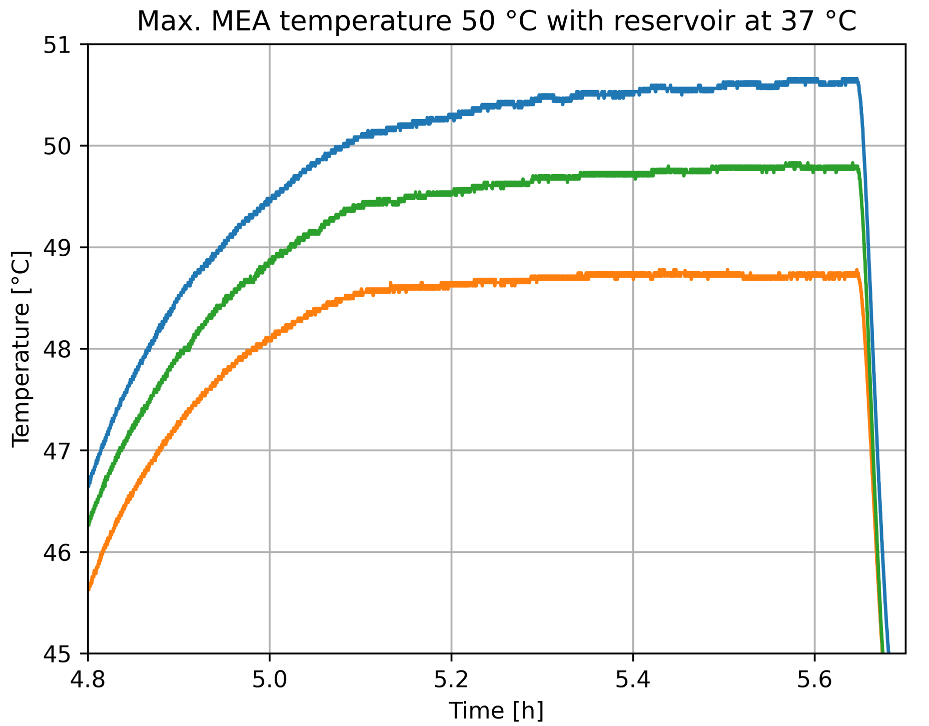


Figure S56: Maximum medium temperature achieved with the reservoir temperature at 37 °C and the thermoelectric devices set to maximum heating.

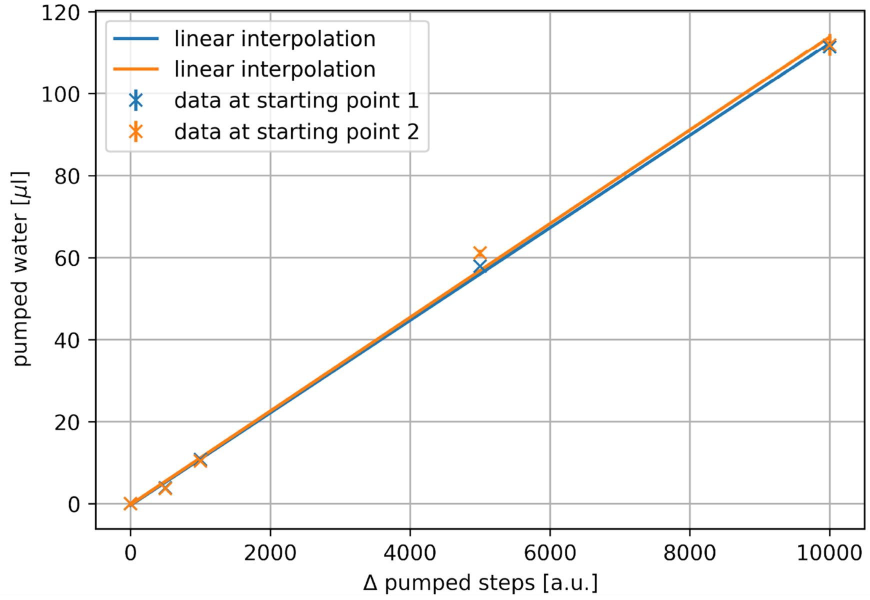


Figure S57: Linearity measurements on the pump, with starting point 1 at 20'000 steps and starting point 2 at 60'000 steps on the motor (n=3).

measurement was repeated 3 times with negligible variations between iterations. The influence of the starting position can be neglected. The results can be seen in Fig. S57.

The hysteresis, the difference between adding and retrieving liquid, of the pump was tested by pumping liquid with 2,500 and 5,000 steps in both directions. The results are shown in Fig. S58 and should be considered when operating the pump without feedback.

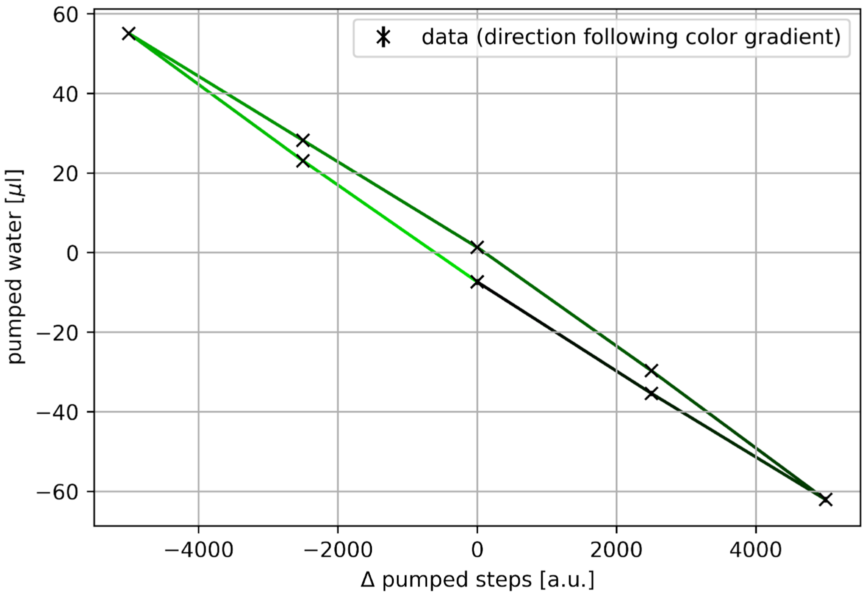


Figure S58: Hysteresis measurement on the pump, direction of pumping and measurement follows the color gradient from dark to light green (n=3). The measurement points are marked with an 'x'.

The minimum dry bath temperature was tested by connecting the Peltier elements to 9 V, while the reservoir was heated to 37 °C. The results are shown in Fig. S59.

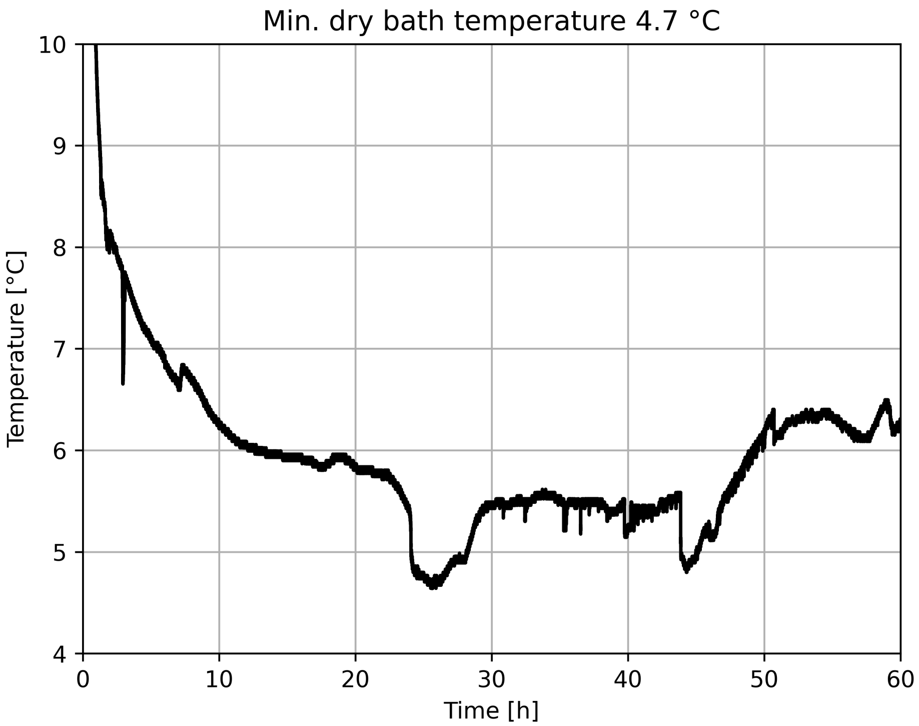


Figure S59: Measurement of the temperature of the liquid in the dry bath falcon tube measured by the liquid temperature sensor usually used for the MEAs.

#### D.3 Spike shapes

Example spike shapes of a single electrode are provided in Fig. S60.

#### D.4 Stimulation

An example trace of 16 evenly spaced pulses at 100 Hz is shown in Fig. S61. To reduce the impact of the stimulation artifact, an active discharge is performed on all electrodes of a stimulated chip. The discharge time is usually set between 0.5 and 1 ms.

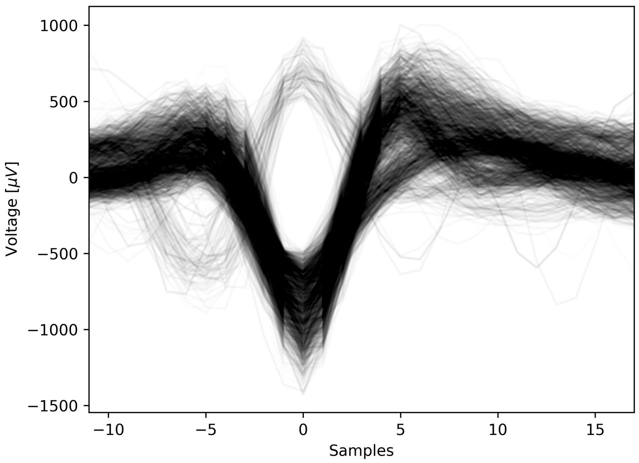


Figure S60: Example spike shapes of a single electrode, detected on the filtered signal with a threshold of 6. All shapes are aligned to the detected maximum.

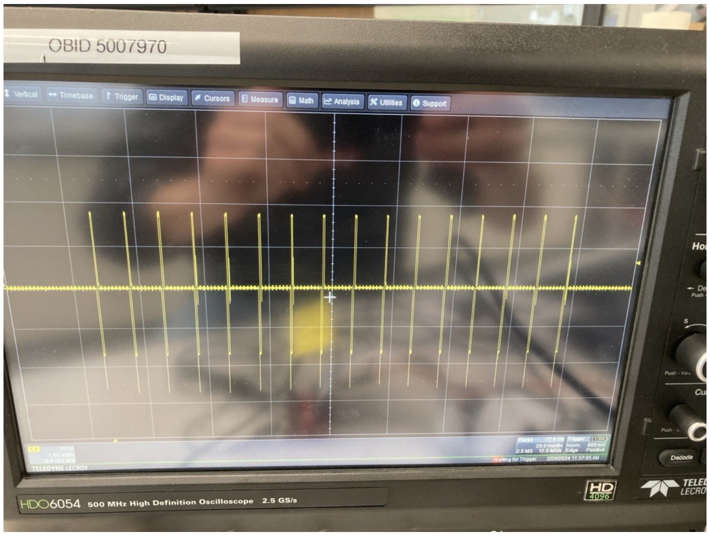


Figure S61: Stimulation pulses.

### E Additional experimental data

Additional experimental data of the experiments shown in the main text are provided in this section.

#### E.1 Commercial system temperature characterization

The performance and stability of the temperature control of inkube is compared with a commercial system (MEA2100-Mini-60-System and MEA2100-System, Multi Channel Systems MCS GmbH, Reutlingen, Germany). The temperature is measured with the inkube PT1000 sensor inside the liquid. To create similar conditions to a mounted a inkube, the PT1000 is attached to a membrane cap reducing evaporation (ALA MEA-MEM, Multi Channel Systems MCS GmbH, Reutlingen, Germany). When the temperature control of the MEA2100 headstage is set to 37 °C, the liquid temperature settles at roughly 3.5 °C below at a ambient temperature of about 23 °C Fig. S62. The start-up is shown in Fig. S63 with an exponential fit.

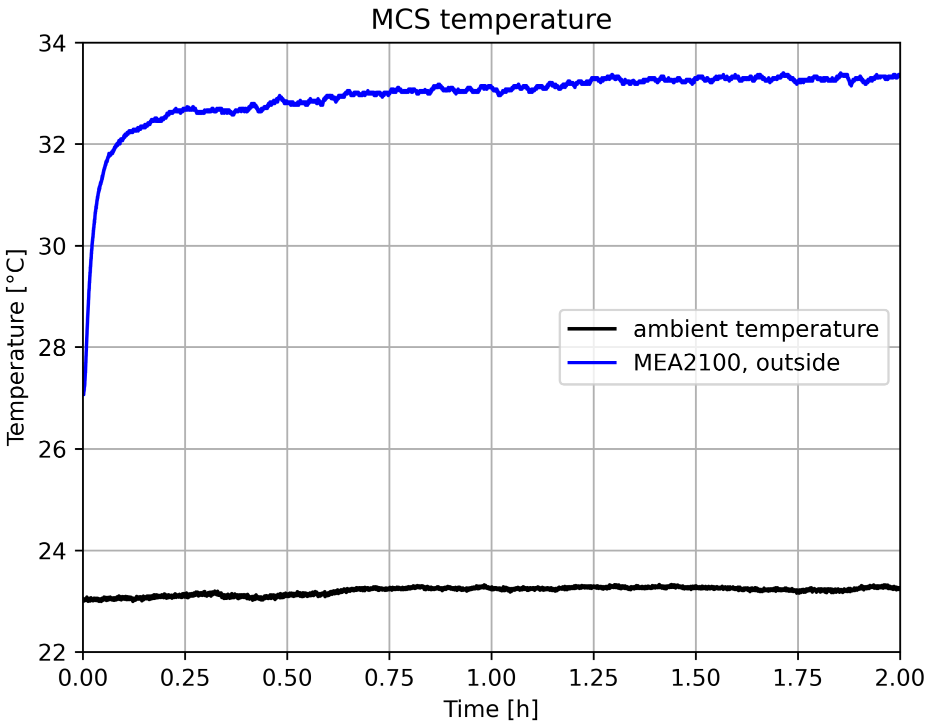


Figure S62: Ambient temperature and liquid temperature of a MEA inside the MEA2100 headstage, with the controller set to 37 °C.

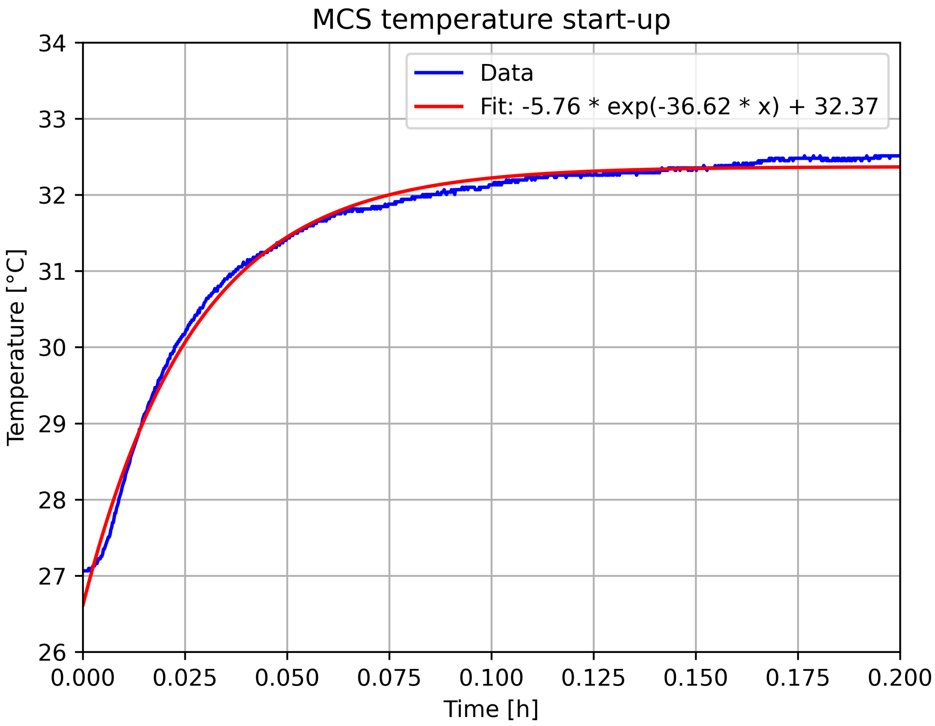


Figure S63: Exponential fit to the start-up of Fig. S62.

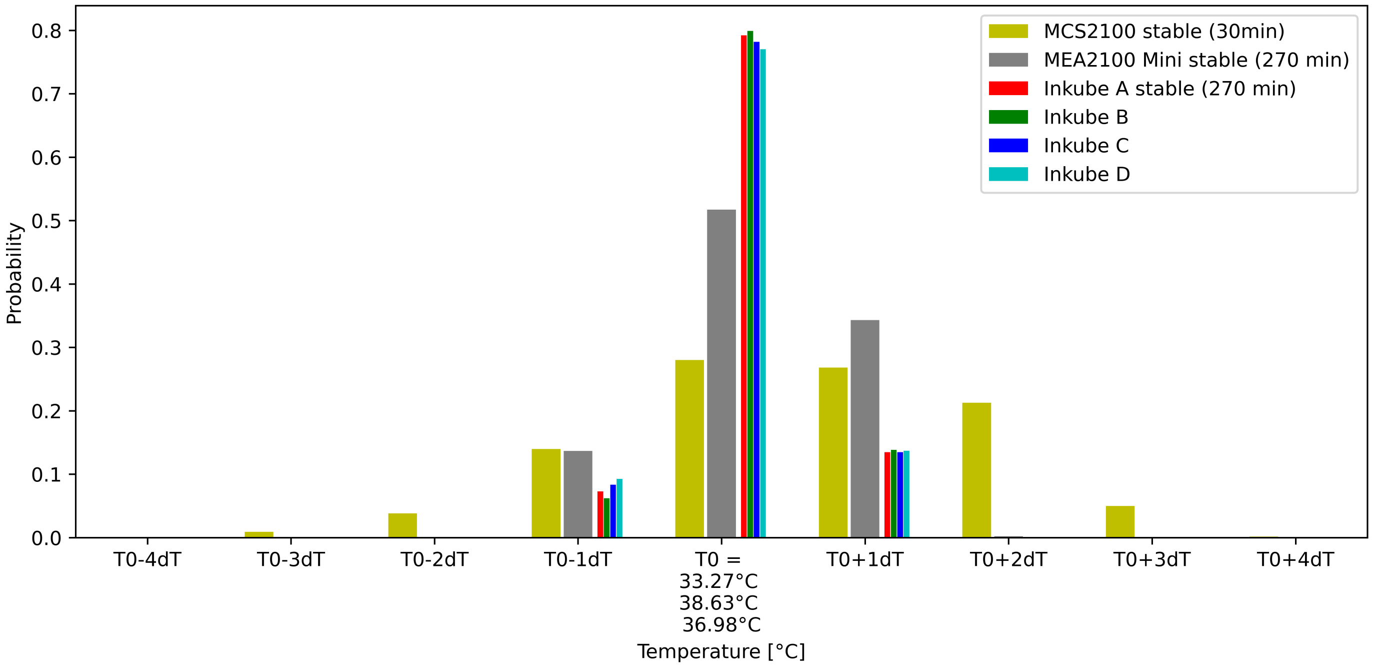


Figure S64: Temperature stability for 3 different devices. MCS2100 is the histogram of Fig. S62 between 90 and 120 min. MCS2100 Mini is the liquid temperature inside an MEA placed in a MEA2100 Mini with set temperature at 37 °C in an incubator (CB 170, BINDER Inc., Bohemia, NY, USA) with set temperature at 35 °C at steady state. The inkube MEAs are a replicate of Fig. 3A for direct comparison. The T0 for the 3 different devices in the x-axis label is listed in order of the histograms from left to right. dT corresponds to 31.7 mK.

#### E.2 Unordered data with temperature steps

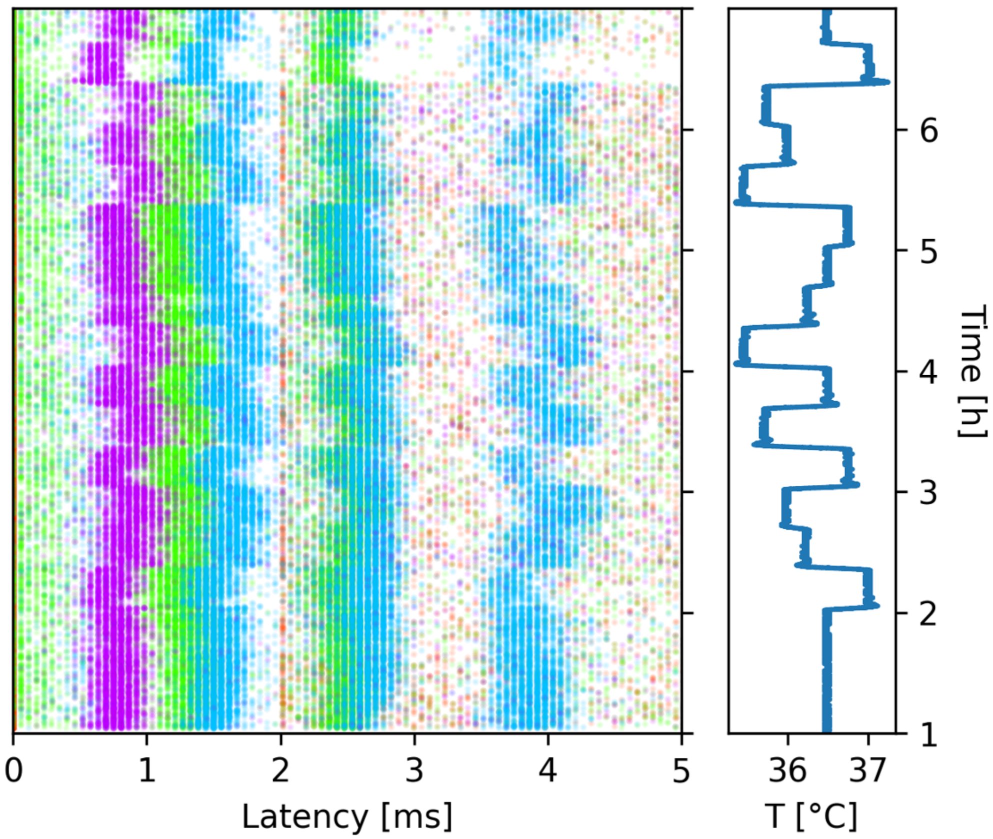


Figure S65: Unordered data from Fig. 4C.

#### E.3 Analysis for additional bands in Fig. 5C.

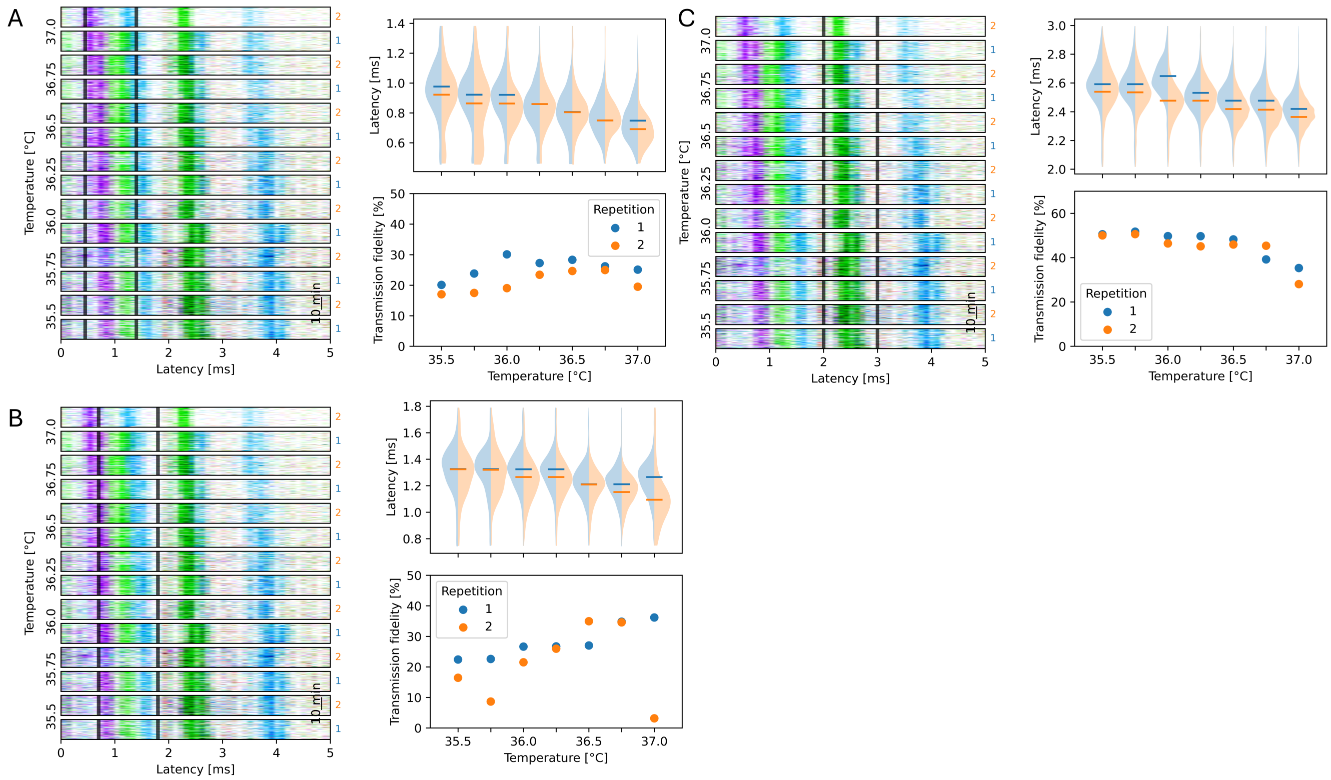


Figure S66: **(A)** Observation electrode 3 (starting at 0), band limits [0.45, 1.4] **(B)** Observation electrode 1 (starting at 0), band limits [0.7, 1.8] **(C)** Observation electrode 1 (starting at 0), band limits [2, 3]

#### E.4 Histogram of network responses and spontaneous recordings at different Magnesium concentrations

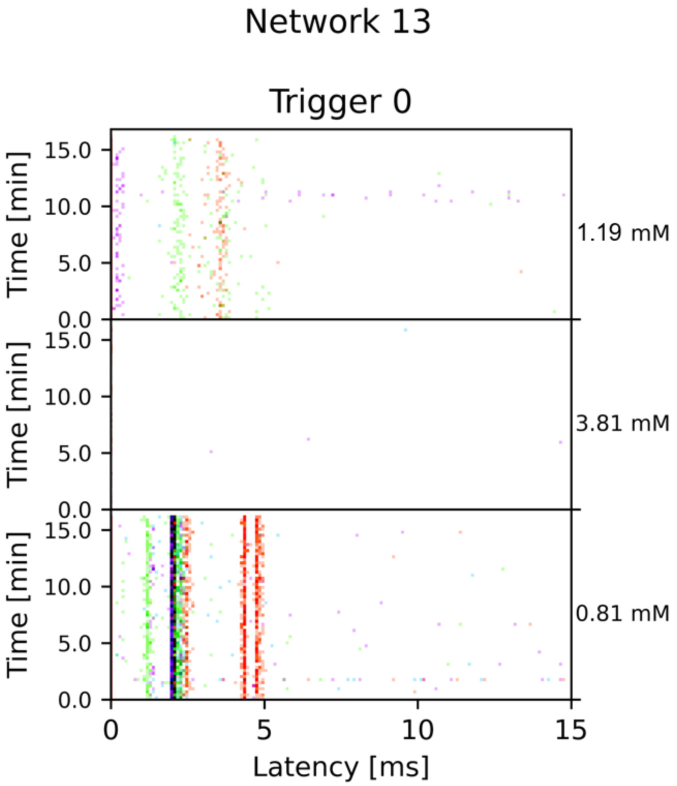


Figure S67: STTRPs of experiment in Fig. 6 at 0.81, 3.81 and 1.19 mM (second measurement after wash-out) before stimulus.

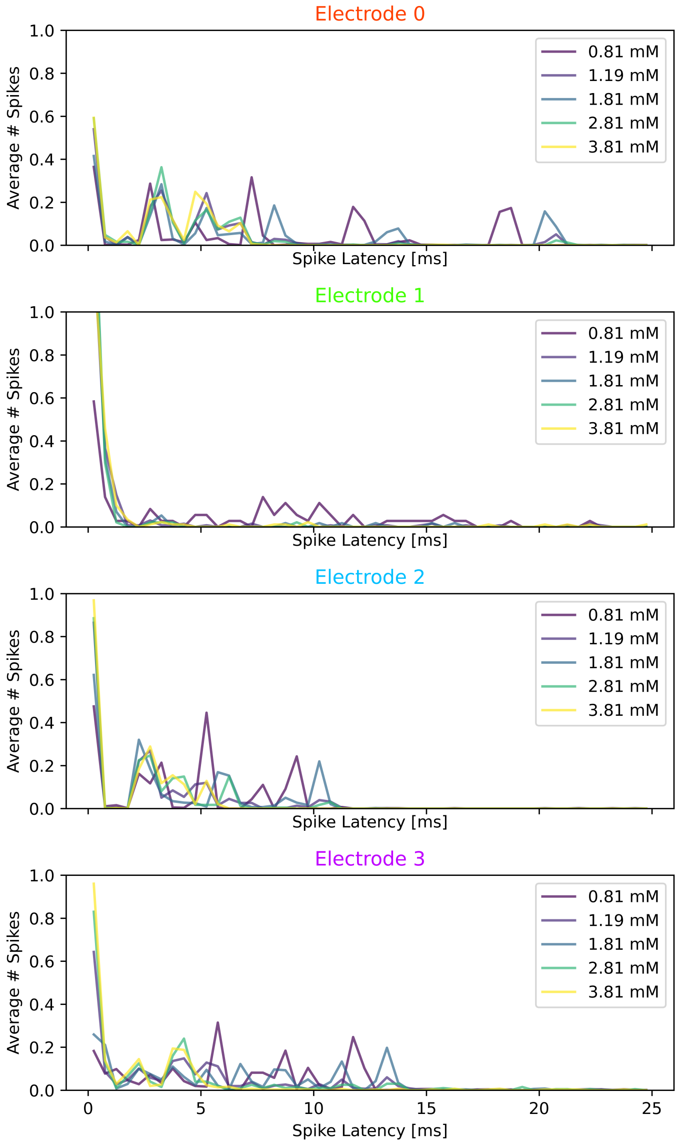


Figure S68: Spike count post stimulus with data from Fig. 6B binned to 0.5 ms.
